# Confounding effects of inferring gene co-expression networks from pooled data from different biological populations

**DOI:** 10.64898/2026.06.23.734063

**Authors:** Rogini Runghen, Tina Eliassi-Rad, Daniel I. Bolnick

## Abstract

Weighted Gene Co-expression Network Analysis (WGCNA) is routinely applied to pooled datasets from multiple biological populations, genotypes, or treatment groups, implicitly assuming a shared module structure across groups. While the distortion of pairwise correlations by pooling heterogeneous groups is well established statistically, three aspects of this problem have received little systematic attention in the context of co-expression network analysis: the extent to which pooling disrupts the discrete module-level community structure inferred by WGCNA; whether this disruption is detectable from the global topology metrics researchers routinely report; and how prevalent the pooling practice is in published multi-group WGCNA studies. Using analytical toy examples and a four-scenario simulation framework, we address all three questions. Module preservation *Z*_summary_ scores declined progressively with between-population divergence, from full preservation under identical populations (mean median *Z*_summary_ = 25.2 ± 3.3, 95% interval 19.0–30.7 across 20 simulation replicates) to substantial disruption when both network structure and mean expression differed (mean median *Z*_summary_ = 11.9 ± 1.0, 95% interval 10.2–13.5). This disruption was undetectable from global topology metrics: modularity and clustering coefficient remained stable across all scenarios, while edge density was sensitive but non-specific. These findings were corroborated in an empirical reanalysis of divergent lake and stream stickleback transcriptomes, where merged analysis collapsed 26 lake-specific and 59 stream-specific modules into only 19 merged modules. A survey of 100 publications found that 78.7% (95% CI 69.4–87.9%) of multi-group WGCNA studies with sufficient methodological reporting used a single merged analysis. Results were robust across network sizes of 250–1,000 genes and rewiring rates of 10–50%. We provide concrete recommendations including module preservation testing in both directions, population-specific baseline networks, and consensus WGCNA as a principled alternative.

## 1 Background

Over the past decade, Weighted Gene Co-expression Network Analysis (WGCNA) has been widely used to analyse gene expression data [Zhang and Horvath, 2005, Langfelder and Horvath, 2008]. The main objective of WGCNA is to infer gene regulation within organisms by identifying clusters (or modules) of genes with similar expression profiles. To achieve this, WGCNA assumes that genes which are co-regulated or functionally related will have correlated transcript abundance, and vice versa. That is, if two genes share regulatory control or contribute to the same biological process, their mRNA levels are expected to co-vary across a sample of varying individuals: individuals with high expression of one gene will tend to have high (or low, in the case of negative co-regulation) expression of the other. Covariation between genes is, in turn, used to infer potential co-regulation between those genes. This covariation may be constitutive—reflecting stable, sample-to-sample differences in the activity of shared regulatory programmes—rather than necessarily implying dynamic temporal fluctuation. Because biological functions usually involve multiple genes acting in concert, pairwise correlations cluster into larger sets called modules. A module is a group of genes more strongly correlated with one another than with genes outside it, and is inferred to share regulatory control, biological function, or both [Langfelder and Horvath, 2008]. Biologists often wish to identify such modules because they can reveal sets of interacting genes—such as the members of a gene regulatory cascade or the components of a shared biological pathway—whose coordinated expression would not be apparent from examining individual genes in isolation.

In the case where data are collected from multiple groups of samples, how should one proceed with using WGCNA and interpreting the outcome? Samples may be grouped based on their exposure to different treatments or environments, or they may come from distinct populations that experienced different environments or had distinct genotypes. Should the two groups be merged to infer a single WGCNA network and set of modules? Merging biological groups for WGCNA assumes that there exists one true module structure that applies to all groups within the dataset, and that merging groups correctly estimates this module structure. When this assumption is valid, the larger sample size enabled by analysing all individuals together provides greater power for accurately estimating covariances and hence network structure. Alternatively, one could split the data and apply WGCNA separately to each group, inferring distinct network architectures, or use consensus WGCNA [Langfelder and Horvath, 2008] to identify modules that are robust across groups.

The merged approach dominates current practice. A survey of 100 randomly selected publications applying WGCNA to multi-group expression data from 2018 to 2025 found that 78.7% (59/75 papers with sufficient methodological reporting, 95% CI 69.4–87.9%) pooled all samples into a single merged analysis; only 14.7% applied WGCNA separately per group, and 6.7% used consensus WGCNA. This prevalence is unsurprising given that the canonical WGCNA tutorial workflow [Langfelder and Horvath, 2008] demonstrates a single pooled network, and researchers following the default example would naturally adopt the merged approach. Nevertheless, the near-universal adoption of this design is concerning because it implicitly assumes a single shared co-expression architecture across all groups—an assumption that is rarely tested and, as we show here, readily violated.

In the context of WGCNA, one assumes that *only* genes which are correlated to one another are co-expressed in the inferred gene network, such that all genes sharing similar expression profiles belong to the same module. However, two important factors must be considered when researchers correlate genes across distinct biological groups. First, the covariance between two genes reflects how they vary together among individuals *within* a biological group; this covariation may differ substantially across populations, genotypes, or treatment conditions [Zhang et al., 2018, Hu et al., 2025]. Second, among-group differences in mean expression can induce spurious correlations between genes that have no direct functional relationship: if two genes are both upregulated in one group relative to another, they will appear correlated in the merged dataset simply because of their shared response to group membership, not because of any shared regulatory mechanism. This form of confounding by group membership is the statistical phenomenon underlying Simpson’s paradox [Simpson, 1951, Pearl, 2009] and has been documented as a major source of spurious associations in pooled omics analyses more broadly [Leek et al., 2010, Johnson et al., 2007]. Numerous genes can differ in mean expression between populations, genotypes, or treatment groups for independent reasons. These coincidental among-group correlations can dominate the merged co-expression structure, displacing genuine within-group regulatory signals.

These two sources of distortion—changes in within-group covariance structure between groups, and spurious among-group correlations—interact in ways that are difficult to diagnose from the merged network alone. In this paper, we use analytical toy examples and a four-scenario simulation framework to quantify these module-level consequences—specifically, the disruption of discrete co-expression modules and its invisibility to standard global network diagnostics. We validate simulation findings using published transcriptome data from genetically divergent lake and stream stickleback populations, survey 100 published multi-group WGCNA studies to characterise current practice, and conclude with concrete recommendations.

## 2 Toy example 1: Populations with the same mean expression but varying type of gene–gene correlation

For the first toy example, we held the mean expression of Gene A and Gene B constant across the T1 and T2 populations and varied only the type of association between Gene A and Gene B. That is, the correlation between Gene *A* and Gene *B* varied from positive through none to negative in the respective populations (see Figures 1 and 2).

**Figure 1:**
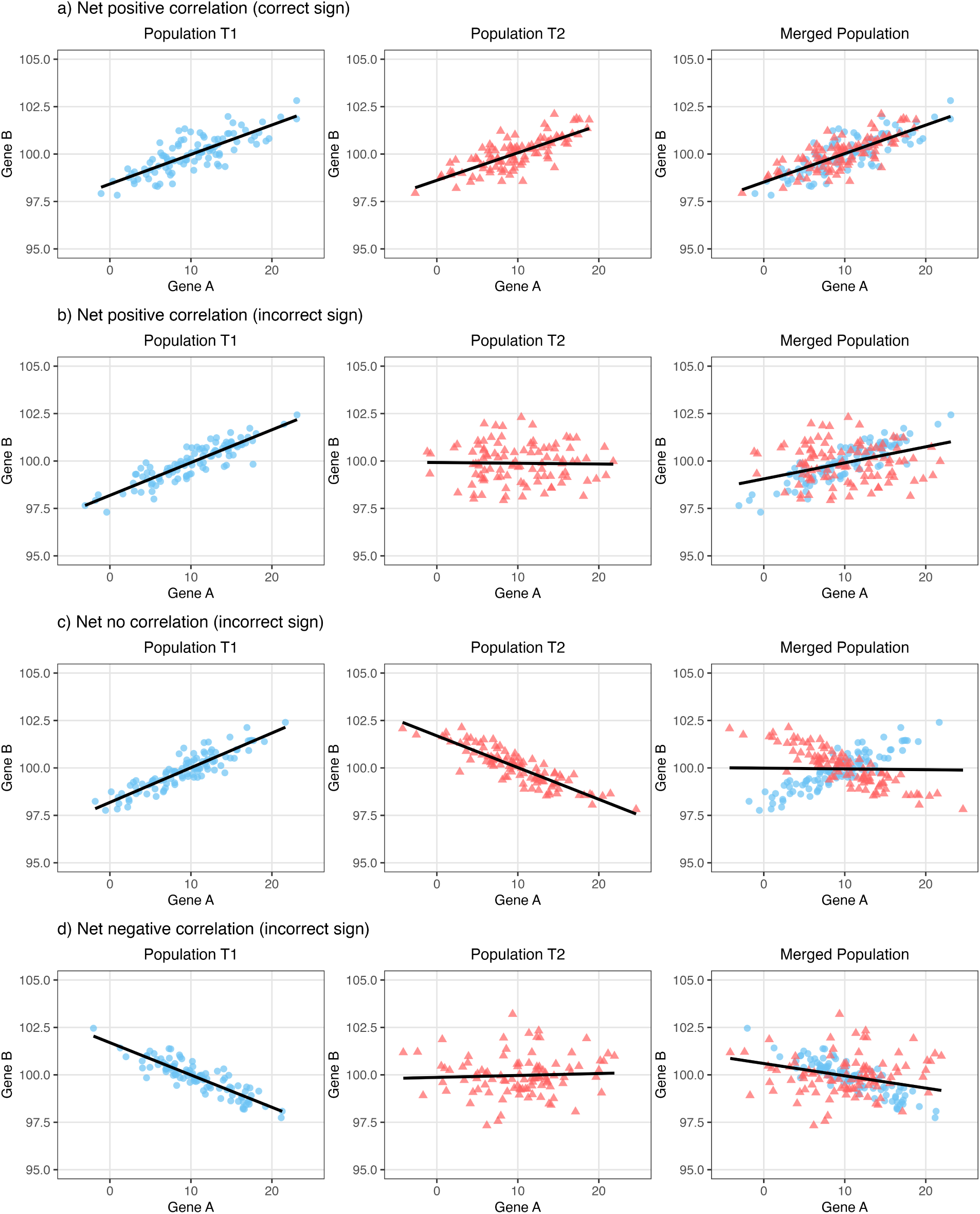
Impact of pooling samples from Populations T1 and T2 on gene–gene correlations, despite equal mean expression. (a) Case where pooled populations and within-population correlations between Gene *A* and *B* remain similar. (b)–(d) Cases where merging populations masks true population-specific relationships and produces distinct correlations.

**Figure 2:**
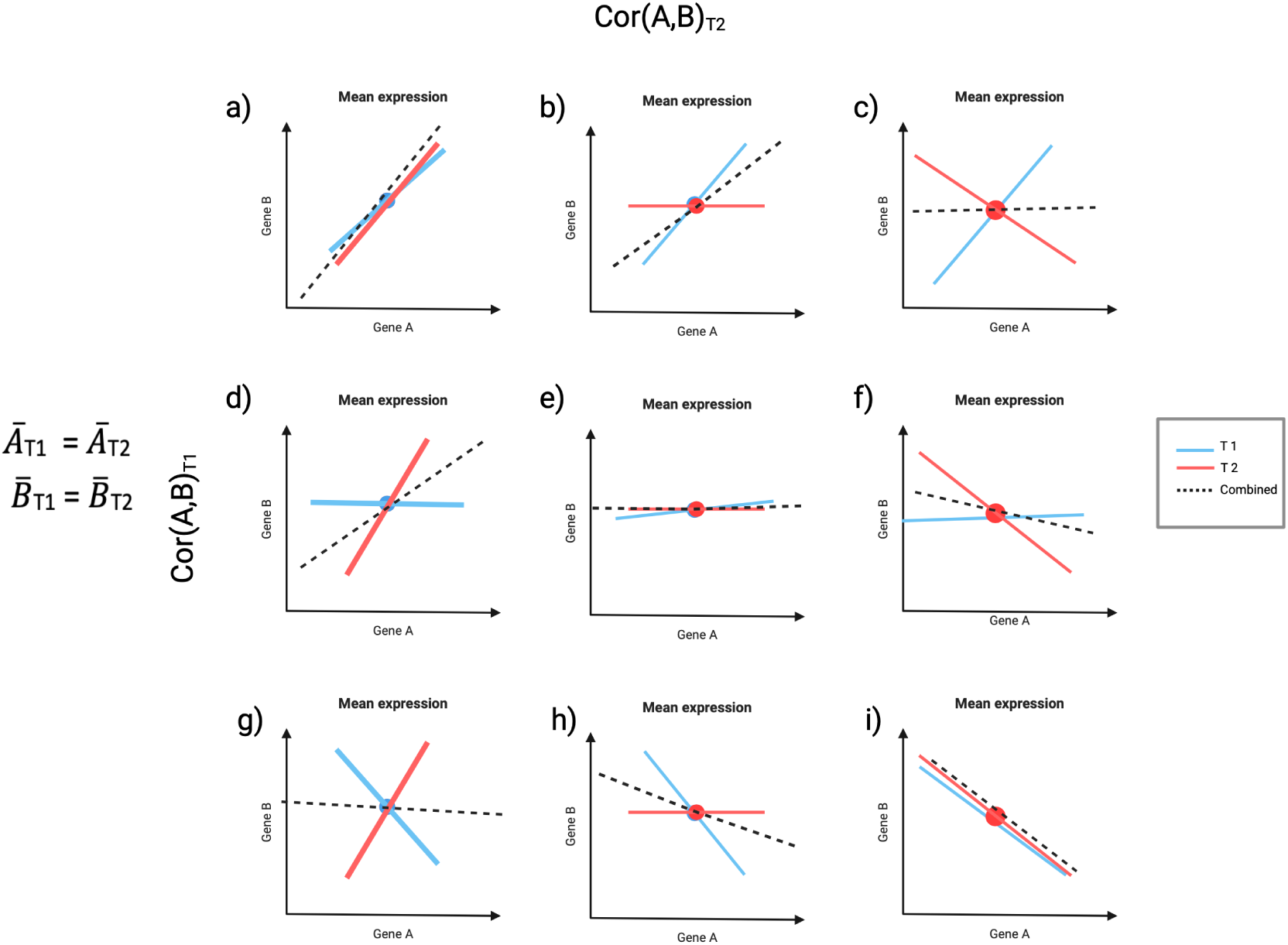
Impact of merging data from two populations with equal mean expression on gene–gene correlations. Panels (a)–(i) show nine possible correlation scenarios between Gene *A* and *B*. Blue and red lines represent correlations in Population T1 and T2 respectively, while dotted lines show correlations in the pooled data. Diagonal panels (a), (e), and (i) demonstrate accurate correlation detection when both populations share similar correlation patterns. Other panels illustrate how ignoring population structure can lead to misidentified correlations when populations exhibit dif-ferent correlation patterns.

As shown in Figure 1, when we merge the samples from two populations with the same mean expression, and same correlation structure, the merged analysis correctly detects the correlation between gene *A* and *B*. In general, because the sample size is made larger by merging, we will have greater power to detect the gene–gene correlation. However, when two populations have different correlations (one strong, one weak, or opposite signs) the merged correlation will be unlike either within-population correlation, as shown in the red box in Figure 1. In these cases the merged analysis correlation will consistently be weaker than one population (and stronger than the other), or appear uncorrelated. In this sense the merged analysis is ‘conservative’, tending to underestimate correlations, but this can result in erroneous gene co-expression network inferences. If sample sizes are unequal for the biological groups, the inferred gene–gene correlation will be biased towards the trend found in the larger sample size. As highlighted in Figure 2, even if samples from two populations have the same mean expression, in some instances, if the type of correlation is different (i.e. positive/negative, none/positive, none/negative or any of the different combinations) in either population, when merging samples from both populations, the net correlation can be incorrectly detected (see Figure S1).

## 3 Toy example 2: Populations with varying mean expression and type of gene–gene associations

In the second toy example, we additionally allowed mean expression to differ between populations: Gene *A* is higher in T1 and Gene *B* is higher in T2, generating a negative between-population association. This scenario is not merely theoretical—between-group differential expression is a defining feature of the contexts in which WGCNA is most commonly applied, affecting hundreds to thousands of genes in disease versus control, genotype, or treatment comparisons [Soneson and Delorenzi, 2013, Schurch et al., 2016, Love et al., 2014, Robinson et al., 2010]. The bias introduced by pooling can be characterised analytically. Consider two equally sized populations (*n* samples each) with within-population Pearson correlations *ρ*_1_ and *ρ*_2_, equal within-population variances *σ*^2^, and population mean differences *δ_A_* = *µ_A,_*_T1_ − *µ_A,_*_T2_ *>* 0 and *δ_B_* = *µ_B,_*_T1_ − *µ_B,_*_T2_ *<* 0. The merged sample covariance between Genes *A* and *B* decomposes as:

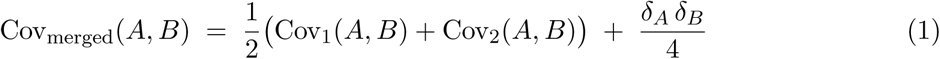

Since *δ_A_ >* 0 and *δ_B_ <* 0, the leverage term 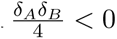 shifts the merged covariance in the negative direction independently of *ρ*_1_ and *ρ*_2_. When |*δ_A_δ_B_*| is large relative to *σ*^2^, this term dominates and the merged correlation reflects between-population mean differences rather than within-population co-expression. Figure 3 illustrates all nine combinations of within-population correlation across the two populations: the pooled correlation (dotted line) is systematically pulled negative relative to the within-population correlations, and only recovers the correct sign when both within-population correlations are themselves sufficiently negative to align with the between-population term. The key consequence is that pooling can (i) recover correct correlations, (ii) create spurious correlations of the wrong sign, or (iii) obscure genuine within-group correlations—and because these pairwise biases form the basis of network inference, they propagate directly to module-level estimates.

**Figure 3:**
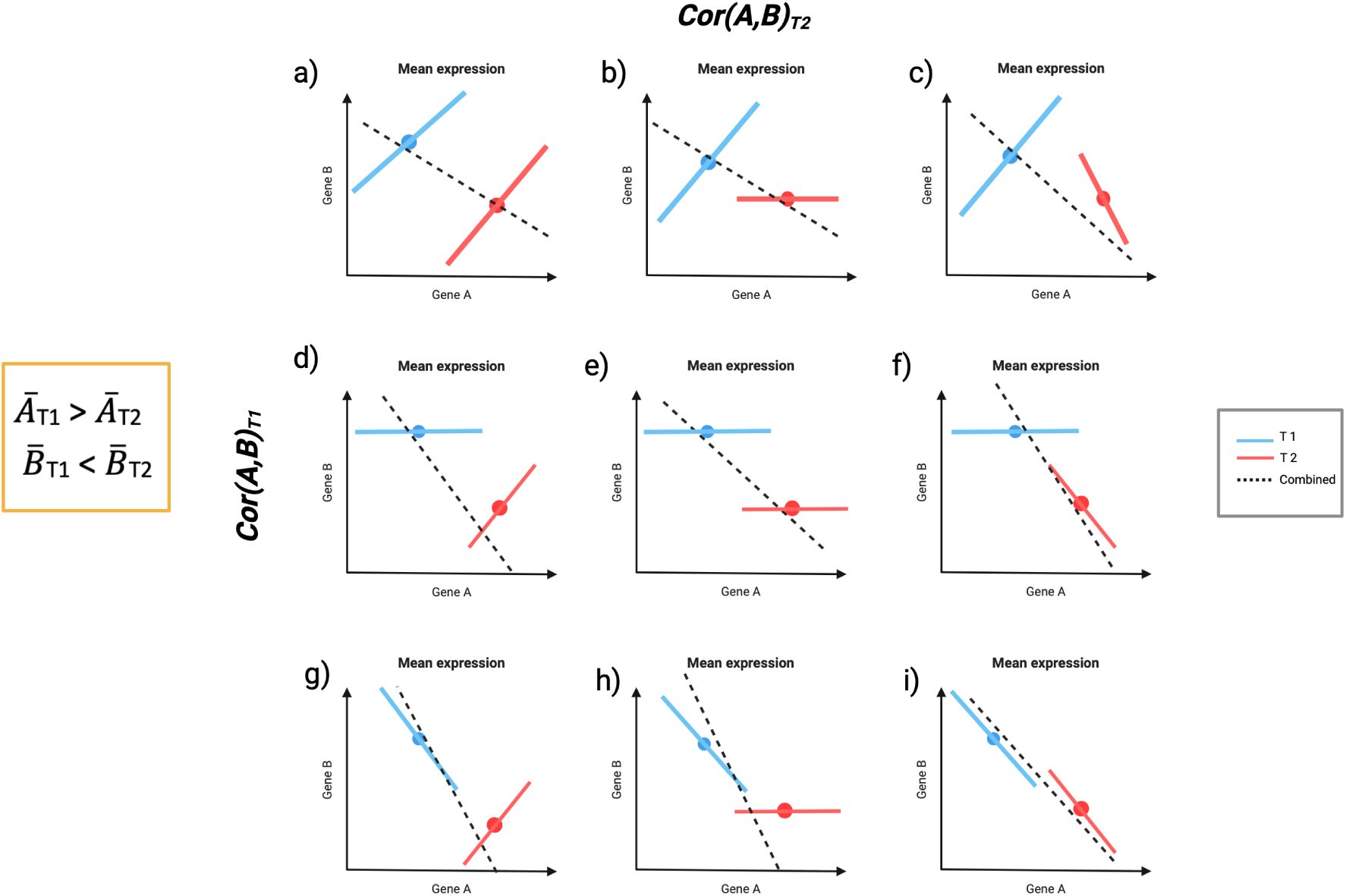
Impact of merging data from populations with different mean expression levels (Gene *A* higher in T1, Gene *B* higher in T2). Panels show nine correlation scenarios between genes, with blue and red lines representing correlations within Population T1 and T2 respectively, and dotted lines showing pooled correlations. When populations have different mean expression, pooled analysis typically misidentifies true gene–gene correlations, except when the within-population correlations are both in the direction of the between-population correlation.

**Figure 4:**
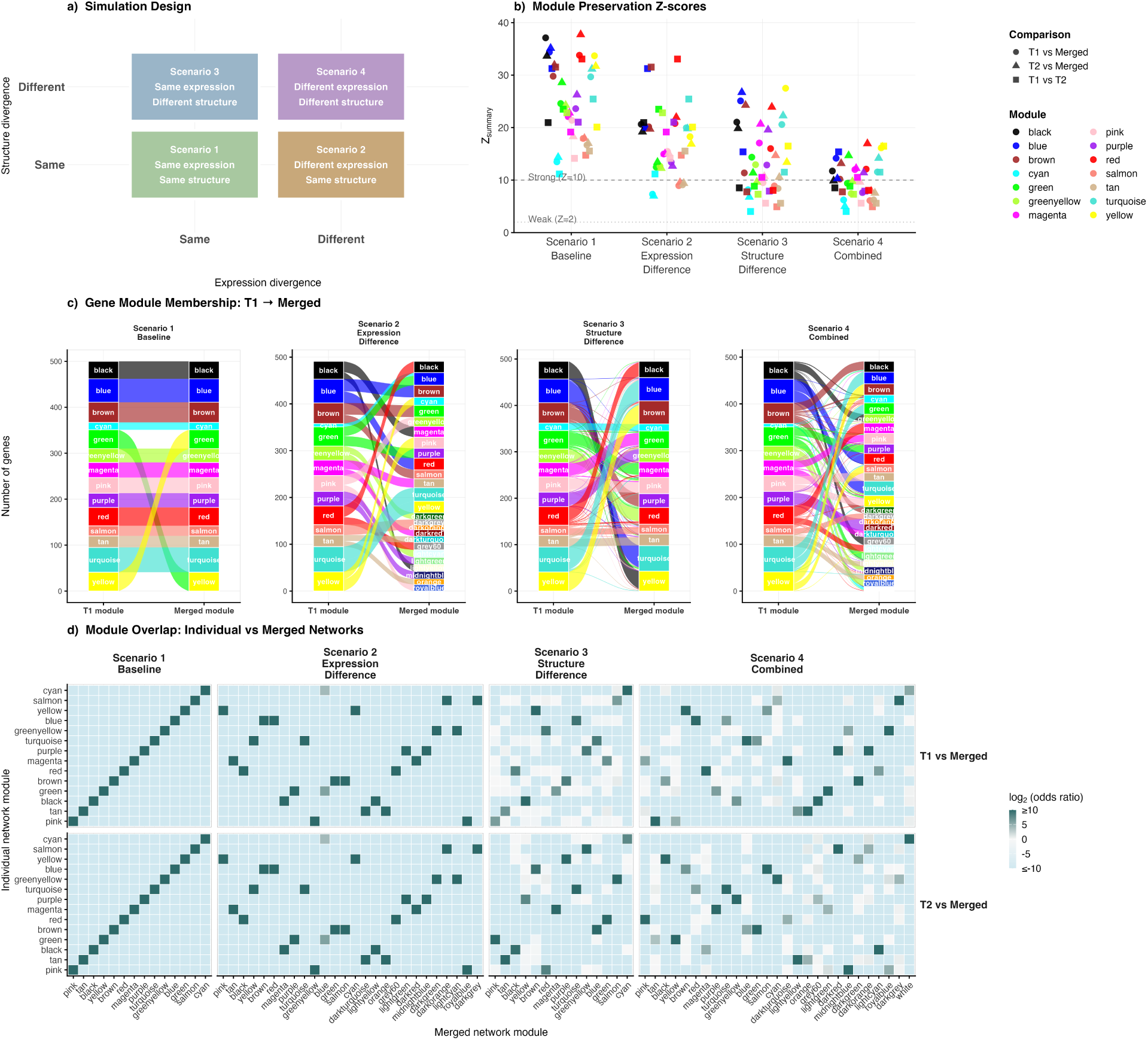
Module-level consequences of cross-population network merging. **a)** Simulation design: four scenarios varying whether populations T1 and T2 share mean expression profiles and co-expression network structure. **b)** Module preservation *Z*_summary_ for all 14 non-grey modules across scenarios. Point colour indicates WGCNA module; shape indicates comparison (circle: T1 versus merged; triangle: T2 versus merged; square: T1 versus T2 as a baseline for natural inter-population divergence). Dashed lines mark strong (*Z*_summary_ = 10) and no-preservation (*Z*_summary_ = 2) thresholds. **c)** Alluvial diagrams showing gene flow from T1 module assignments to merged network assignments. Crossing flows indicate membership reorganisation; flow width is proportional to gene count. **d)** Module overlap heatmaps (log_2_ odds ratio, Fisher’s exact test) for T1 versus merged (upper) and T2 versus merged (lower). Dark teal indicates strong enrichment; a near-diagonal pattern indicates clean module recovery.

**Figure 5:**
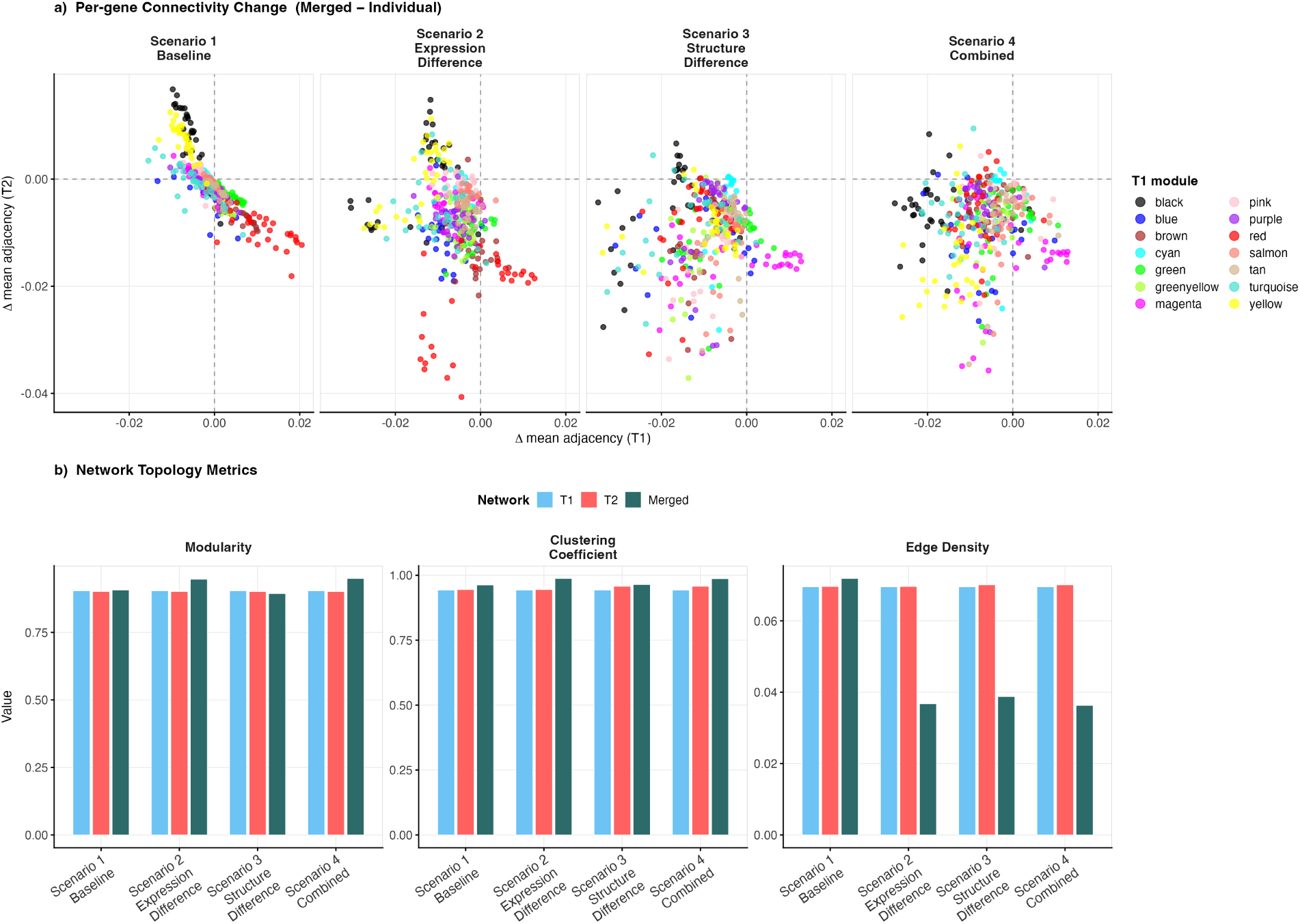
Network-level consequences of cross-population network merging. **a)** Per-gene connectivity change (Δ mean adjacency weight) between each individual and the merged network, for T1 (x-axis) and T2 (y-axis). Points are coloured by T1 module assignment; grey genes excluded. Displacement from the origin indicates genes gaining or losing co-expression partners upon merging. Scenario 1 shows minimal displacement. Scenarios 2–4 show increased spread, with Scenario 3 exhibiting the widest dispersion and strongest module-specific directionality. **b)** Global topology metrics—modularity, clustering coefficient, and edge density—for T1 (light blue), T2 (red), and merged (dark teal) across all four scenarios. Modularity and clustering coefficient remain stable across all scenarios even when module membership is substantially disrupted (cf. Figure 4). Edge density drops markedly in Scenarios 2, 3, and 4 (merged ∼0.036–0.039 versus ∼0.070 in each individual network for Scenarios 2 and 3; merged = 0.036 for Scenario 4), but only Scenarios 3 and 4 show module disruption, demonstrating that edge density is a sensitive but non-specific diagnostic that cannot distinguish expression-driven from structure-driven network changes.

## 4 Simulating gene co-expression networks and RNA-Seq data with varying network structure and mean expression

The gene–gene associations estimated by WGCNA, and the co-expression modules derived from them, are directly determined by the expression data provided as input. In this section, we show that merging samples from two different populations—with either varying network structures or varying mean expression (or both)—can affect the modules identified when using WGCNA. To do so, we simulated gene co-expression networks and their subsequent RNA-Seq data under four scenarios that systematically vary whether the two populations share the same network structure, the same mean expression profiles, both, or neither:

1. **Scenario 1**: Two populations with similar network structure and the same mean expression profiles. This is the situation assumed by many WGCNA users;
2. **Scenario 2**: Two populations with similar network structure but different mean expression profiles;
3. **Scenario 3**: Two populations with different network structure but similar mean expression profiles;
4. **Scenario 4**: Two populations with different network structure and different mean expression profiles, where approximately half of genes have higher mean expression in T1 and the remaining half have higher mean expression in T2 (bidirectional symmetric shift). This models the more typical pattern of differential expression observed in disease versus control or genotype comparisons, where both populations contribute upregulated and downregulated genes.

For each scenario, we simulated three networks: two population-specific networks (T1 and T2), each comprising 150 samples and 500 genes, and a third network derived from merging samples from both populations (300 samples total). The underlying network topology for each population was generated using a scale-free graph via the Barabási–Albert preferential attachment model [Barabási and Albert, 1999], with each new node attaching to *m* = 3 existing nodes. Edge weights were randomly assigned from a uniform distribution over [0.6, 0.9] with a randomly assigned sign (positive or negative), reflecting the mixture of activating and repressive gene–gene relationships commonly observed in biological networks.

For scenarios requiring different network structures (Scenarios 3 and 4), the second population’s network was modified by randomly removing 30% of edges and replacing them with new edges between randomly selected gene pairs, thereby altering the network topology while preserving the overall edge count. To simulate expression data with realistic co-expression community structure, gene communities were first identified in the Barabási–Albert graph using the Louvain community detection algorithm [Blondel et al., 2008], treating edge weights as absolute values. A block-diagonal covariance matrix was then constructed from these community assignments: gene pairs within the same community were assigned a within-community Pearson correlation of *ρ* = 0.85, while gene pairs belonging to different communities were treated as independent (*ρ* = 0). The covariance matrix was constructed from community membership rather than directly from BA edge weights for two reasons. First, translating arbitrary signed edge weights into a valid positive definite covariance matrix is non-trivial without regularisation that would introduce additional free parameters. Second, WGCNA module detection operates on the correlation structure of the expression data rather than on any underlying graph topology; the relevant quantity for evaluating the effects of pooling is therefore the covariance structure of the simulated expression data. The BA graph serves to motivate a community structure with realistic degree heterogeneity at the level of community sizes, while the block-diagonal covariance ensures well-defined, interpretable modules against which preservation can be assessed. This approach guarantees a positive definite covariance matrix (minimum eigenvalue = 1 − *ρ* = 0.15 *>* 0) without requiring regularisation; a safeguard to shift any negative eigenvalues was nonetheless included in the implementation (see Supplementary Material).

For Scenarios 3 and 4, two independent modifications were applied to the second population prior to covariance construction: 30% of edges in the BA graph were rewired (preserving edge count), and 30% of gene community memberships from the original Louvain partition were randomly reassigned to different communities. The covariance matrix for T2 was then constructed from these perturbed community assignments (see Supplementary Material for full details of the covariance construction and regularisation procedure).

RNA-Seq counts were drawn from a multivariate normal with gene-wise log-scale means ∼ *N* (8, 1^2^) and the community-derived covariance described above. Counts were obtained by exponentiation and rounding. In Scenarios 2 and 4, bidirectional mean expression differences were introduced between populations. For each gene, a direction (+1 or −1, each with probability 0.5) was independently assigned, and the magnitude of the shift was drawn from |*N* (2, 0.5^2^)|. Approximately half of genes therefore had higher mean expression in T2 and the remainder in T1, yielding near-identical overall mean expression across populations. The resulting median absolute pairwise gene–gene correlation was approximately 0.05–0.06 across all scenarios, providing a realistic but detectable co-expression signal.

The merged dataset was constructed by combining the expression matrices of T1 and T2 into a single pooled matrix (300 samples × 500 genes) and passing it directly to WGCNA, mirroring the common practice of treating multi-population datasets as a single cohort without population-aware pre-filtering. A separate correlation threshold network (|*ρ̂*| *>* 0.6) was constructed from the pooled data using the igraph package [Csardi and Nepusz, 2006] for visualisation purposes only and was not used in the WGCNA module detection analysis described below (see Supplementary Material).

## 5 Comparing networks and modules inferred from merged populations

To explore how merging gene co-expression networks from two different biological populations can impact the modules identified, we compared the modules identified in the population-specific networks (T1 and T2) with those identified in the network constructed from the pooled data, across all four scenarios. For each network, we first characterised global topology, then assessed the quality and coherence of the modules detected, and finally quantified the degree to which modules identified in the individual population networks were preserved—or reorganised—in the merged network.

### 5.1 Network Structure Analysis

For each network, we quantified three topology metrics using the igraph package [Csardi and Nepusz, 2006]. Modularity was computed using the Louvain community detection algorithm [Blondel et al., 2008] to assess the strength of community structure. The global clustering coefficient was calculated using the transitivity method to measure local network connectivity. Network density was computed as the ratio of observed to possible edges to evaluate overall connection strength. These metrics were computed from the WGCNA soft-thresholded adjacency matrices (see Module Detection section below) converted to igraph objects, for each of the three networks (T1, T2, and merged) across all four scenarios, to characterise how pooling samples alters global network topology independently of the module-level changes considered below.

### 5.2 Module Detection

For each network, modules were identified using WGCNA [Langfelder and Horvath, 2008]. Adjacency matrices were constructed using a signed hybrid network type with a soft-thresholding power of *β* = 6, set *a priori* to ensure a comparable scale-free fit across all three networks within each scenario. Topological Overlap Matrices (TOM) were then computed from the adjacency matrices using a signed TOM type, and genes were clustered using average linkage hierarchical clustering on the TOM-based dissimilarity. Modules were identified using the dynamic tree cutting algorithm [Langfelder et al., 2008] with default sensitivity (deepSplit = 2) and a minimum module size of 10 genes. Genes not assigned to any module were assigned to the grey (background) module and excluded from downstream comparisons.

### 5.3 Module Quality Assessment

Module quality was assessed using two complementary metrics. The variance explained by each module eigengene was calculated as the ratio of the eigengene variance to the total expression variance, defined as the sum of per-gene sample variances across all genes in the dataset, where the eigengene is the first principal component of the module expression matrix. Module density was computed as the mean pairwise WGCNA adjacency weight among all genes assigned to a given module, providing a direct measure of intra-module co-expression strength. Together, these metrics allowed us to evaluate whether pooling samples reduces the coherence of the modules detected, beyond any changes in their membership.

### 5.4 Module Preservation and Overlap Analysis

To quantify the degree to which modules identified in the individual population networks were preserved in the merged network, we employed two complementary approaches. Module preservation statistics were computed using WGCNA’s `modulePreservation` function [Langfelder et al., 2011], with T1 and T2 each used in turn as the reference network and the merged network as the test network, using 100 permutations with a signed hybrid network type. To distinguish confounding introduced by merging from the natural inter-population divergence, module preservation was additionally computed between T1 and T2 directly, providing a baseline against which any preservation losses attributable to the merging procedure could be assessed. The *Z*_summary_ statistic was extracted as the primary summary measure of module preservation, where *Z*_summary_ *>* 10 indicates strong preservation and *Z*_summary_ *<* 2 indicates no preservation[Langfelder et al., 2011].

Pairwise module overlap between networks was assessed using Fisher’s exact test applied to contingency tables of gene-to-module assignments, for each module pair across the T1-versus-merged and T2-versus-merged comparisons. This allowed us to determine whether the gene membership of a given module in the individual network overlapped with that of any module in the merged network more than expected by chance. *p*-values were corrected for multiple testing using the Benjamini–Hochberg procedure[Benjamini and Hochberg, 1995].

To quantify replicate-level variability in all reported statistics, each scenario was run across 20 independent simulation replicates using seeds 42 + 1,000*k* for *k* = 1*, . . .,* 20. For each replicate, module preservation statistics were computed as described above using 100 permutations. Key outcomes—median *Z*_summary_, number of strongly preserved modules, and edge density—are reported as means with standard deviations and 95% intervals across replicates. The single-seed results presented in the main figures use seed = 42; full replicate-level summaries are provided in the Supplementary Material (Table S4).

### 5.5 Consensus WGCNA comparison

To evaluate whether consensus WGCNA recovers individual population module structure more faithfully than a merged analysis, we additionally applied consensus WGCNA [Langfelder and Horvath, 2008] to the Scenario 4 data using blockwiseConsensusModules() with the same soft-thresholding power (*β* = 6), minimum module size (10 genes), and network type (signed hybrid) as the individual and merged analyses. Consensus WGCNA constructs a consensus topological overlap matrix by taking the component-wise minimum of the per-group TOMs, identifying modules that are consistently present across both T1 and T2. Module preservation statistics were then computed comparing the consensus modules against each individual network (T1 and T2 separately) and against the merged network, using the same permutation-based framework (*n* = 100 permutations) described above. The key comparison of interest was whether consensus modules showed higher preservation in the individual networks than merged modules did.

### 5.6 Empirical validation using lake–stream stickleback co-expression data

To assess whether our simulation findings generalise to empirically derived data, we applied the merged and split WGCNA approach to published TagSeq expression data from wild lake and stream threespine stickleback (*Gasterosteus aculeatus*) from Roberts Lake and Stream, Vancouver Island [Lohman et al., 2017]. These populations are genetically and phenotypically divergent, with documented differences in immune gene expression, parasite communities, and morphology [Lohman et al., 2017]. As such, the expression data from Lohman et al. [2017] provides a biologically grounded two-population comparison with both mean expression and structural co-expression differences, which is the empirical analogue of Scenario 4.

Wild lake (*n* = 17) and stream (*n* = 14) fish were analysed. Raw counts were normalised using variance-stabilising transformation in DESeq2 [Love et al., 2014] with batch correction (design: ∼Batch + Treatment), and genes with mean count below 1 were excluded, retaining 9,911 genes. WGCNA was applied separately to each population and to the merged dataset using the same parameters as the original analysis [soft-thresholding power = 6, minimum module size = 30, merge cut height = 0.20; Lohman et al., 2017]. Module preservation was assessed using modulePreservation() with 100 permutations and a signed hybrid network type.

## 6 Results

### 6.1 Scenario 1: Baseline Condition

Under the baseline condition, where both populations shared the same network structure and mean expression profiles, the merged network’s global topology closely matched that of the individual population networks (Figure S2). Modularity was high and comparable across all three networks (*Q* = 0.904 for T1, *Q* = 0.901 for T2, and *Q* = 0.907 for the merged network). Global clustering coefficients were similarly consistent (*T* 1 = 0.943, *T* 2 = 0.945, merged = 0.963), as were network densities (*T* 1 = 0.070, *T* 2 = 0.070, merged = 0.072). These results confirm that, in the absence of any between-population differences (in means or covariances), pooling samples provides accurate estimates of the overall connectivity structure of the network.

At the module level, 14 non-grey modules were identified in each of T1, T2, and the merged network. Module preservation was strong across all 14 modules in both comparisons: *Z*_summary_ ranged from 13.5 to 37.1 (median = 24.1) for T1 versus merged, and from 14.3 to 37.7 (median = 27.4) for T2 versus merged, with all 14 modules exceeding the strong preservation threshold in both cases. Across 20 independent simulation replicates, mean median *Z*_summary_ for T1-versus-merged was 25.2 ± 3.3 (95% interval 19.0–30.7; Table S4). The direct T1-versus-T2 comparison yielded similarly high preservation (range: 11.2–33.1, median = 21.0, all 14 modules strongly preserved), confirming that the high preservation in the merged network reflects genuine biological similarity between populations.

### 6.2 Scenario 2: Mean Expression Differences

When populations shared the same co-expression architecture but differed in mean expression, module preservation in the merged network was reduced relative to the baseline (Figure S3). For T1, 11 of 14 modules met the strong preservation threshold (*Z*_summary_ *>* 10; range: 7.3–20.8, median = 15.21), and the pattern was near-identical for T2 (11 of 14 strongly preserved; range: 7.0–22.0, median = 15.06). Across 20 replicates, mean median *Z*_summary_ for T1-versus-merged was 15.1± 1.2 (95% interval 12.8–17.2; Table S4). The direct T1-versus-T2 comparison confirmed strong preservation across all 14 modules (median *Z*_summary_ = 21.00), indicating that the reduction in merged network preservation is attributable to the pooling process rather than to inherent biological differences between populations.

Global topology metrics were largely maintained: modularity (T1 = 0.904, T2 = 0.901, merged = 0.947) and clustering coefficient (T1 = 0.943, T2 = 0.945, merged = 0.987) were stable or slightly elevated in the merged network. However, edge density dropped substantially relative to the individual networks (T1 = 0.070, T2 = 0.070, merged = 0.037), indicating that bidirectional expression differences reduce the number of strong gene–gene correlations without systematically distorting which genes are co-expressed. Edge density is therefore sensitive but non-specific: it signals that populations differ without distinguishing expression-driven density loss from structure-driven module disruption.

### 6.3 Scenario 3: Network Structure Differences

When the two populations differed in network structure but shared the same mean expression profiles, modularity remained broadly comparable (T1: *Q* = 0.904; T2: *Q* = 0.901; merged: *Q* = 0.894) and clustering coefficients were similar (*T* 1 = 0.943, *T* 2 = 0.957, merged = 0.964). However, the merged network exhibited a pronounced reduction in edge density (merged = 0.039) compared to both T1 (0.070) and T2 (0.070), representing an approximately 44% decrease (Figure S4).

At the module level, *Z*_summary_ for T1-versus-merged ranged from 8.2 to 27.5 (median = 13.7), with 10 of 14 modules strongly preserved; 4 fell into the moderate range (2 *< Z*_summary_ ≤ 10). For T2-versus-merged, preservation was somewhat higher (range: 6.8–26.7, median = 17.1, 12/14 strongly preserved). Across 20 replicates, mean median *Z*_summary_ for T1-versus-merged was 16.8±1.5 (95% interval 13.8–19.0; Table S4). The direct T1-versus-T2 comparison revealed substantially lower preservation (range: 4.0–16.5, median = 8.0, only 4 of 14 strongly preserved), confirming genuine structural divergence: the merged network represents a compromise between two structurally distinct populations rather than faithfully reproducing either.

### 6.4 Scenario 4: Combined Network Structure and Expression Differences

When both network structure and mean expression differed between populations, module disruption was greatest of all four scenarios (Figure S5). Global topology metrics remained stable (modularity: *T* 1 = 0.876, *T* 2 = 0.875, merged = 0.938; clustering coefficient: *T* 1 = 0.912, *T* 2 = 0.929, merged = 0.972), while edge density again dropped substantially (T1 = 0.071, T2 = 0.073, merged = 0.036), consistent with structural cancellation compounded by expression-level noise.

At the module level, *Z*_summary_ for T1 ranged from 6.1 to 16.1 (median = 9.99), with 7 of 14 modules strongly preserved. For T2, values ranged from 5.0 to 17.0 (median = 10.85), with 8 of 14 strongly preserved. Across 20 replicates, mean median *Z*_summary_ for T1-versus-merged was 11.9 ± 1.0 (95% interval 10.2–13.5; Table S4)—the lowest of all scenarios, with a 95% interval that does not overlap with Scenario 1 (19.0–30.7). The near-symmetry between T1 and T2 confirms that bidirectional expression differences do not introduce systematic directional bias toward either population’s architecture. The direct T1-versus-T2 comparison confirmed genuine structural divergence (range: 4.0–16.5, median = 7.97, 4/14 modules strongly preserved); adding expression differences to structural divergence therefore compounds disruption modestly—mean median *Z* declining from 16.8 ± 1.5 in Scenario 3 to 11.9 ± 1.0 in Scenario 4.

### 6.5 Consensus WGCNA recovers individual network structure more faithfully than merged analysis in Scenario 4

To provide an empirical evaluation of consensus WGCNA as a practical alternative to merged analysis, we applied blockwiseConsensusModules() to the Scenario 4 data—the scenario where merged analysis showed the greatest module disruption—and compared the resulting consensus modules against both individual networks. Consensus WGCNA identified 14 modules, identical in number to the individual population networks. These modules showed near-complete preservation in both individual networks: all 14 consensus modules were strongly preserved in T1 (median *Z*_summary_ = 24.74) and all 14 were strongly preserved in T2 (median *Z*_summary_ = 21.80). This stands in stark contrast to the merged analysis, in which only 7 of 14 T1 modules were strongly preserved (median *Z*_summary_ = 9.99) and 8 of 14 T2 modules were strongly preserved (median *Z*_summary_ = 10.85).

The preservation scores achieved by consensus WGCNA (median *Z* = 24.74 for T1, 21.80 for T2) are comparable to those observed in Scenario 1, where populations were identical and merged analysis itself produced strong preservation (median *Z*_summary_ = 24.1). This indicates that consensus WGCNA effectively eliminates the module disruption introduced by pooling populations with different co-expression architectures, recovering the shared biological signal that the merged approach fails to identify. However, these results should be interpreted with caution, as consensus analysis by design reflects only the shared network structure across populations and may therefore overlook biologically meaningful differences specific to one group.

### 6.6 Empirical validation: module disruption in lake–stream stickleback data

WGCNA applied separately to each population identified 26 lake modules and 59 stream modules. When applied to the merged lake and stream expression matrix, WGCNA identified only 19 modules, collapsing the richer population-specific structure. Lake module preservation in the merged network was moderate (median *Z*_summary_ = 10.25; 15/26 strongly preserved), and stream module preservation was lower (median *Z*_summary_ = 9.14; 26/59 strongly preserved). As a biological baseline, lake modules showed poor preservation in the stream network (median *Z*_summary_ = 5.23; 5/26 strongly preserved), confirming that these populations have genuinely divergent co-expression architectures (Figure S8).

### 6.7 Overall Patterns

Taken together, the simulation results reveal a clear and consistent pattern across all four scenarios. Global network topology—modularity and clustering coefficient—was largely preserved in the merged network across all scenarios, while edge density dropped substantially in Scenarios 2, 3, and 4 (merged network retaining approximately 36–39 edges per 1,000 possible pairs compared to approximately 70 in either individual network). Mean median *Z*_summary_ for T1-versus-merged declined progressively from 25.2 ± 3.3 (Scenario 1) to 11.9 ± 1.0 (Scenario 4), with Scenarios 2 and 3 showing comparable intermediate disruption (overlapping 95% replicate intervals) and Scenario 4 showing consistently greater disruption whose interval (10.2–13.5) does not overlap with Scenario 1 (19.0–30.7; Table S4).

Consensus WGCNA, applied to the Scenario 4 data, recovered all 14 modules from both populations with strong preservation (median *Z*_summary_ = 24.74 for T1; 21.80 for T2), demonstrating that the disruption introduced by merged analysis is not an inherent consequence of between-group differences but is specifically attributable to the pooling procedure. These simulation patterns were corroborated by the stickleback empirical data [Lohman et al., 2017]: merged analysis recovered only 19 modules compared to 26 and 59 in the population-specific networks, with moderate preservation in the merged network (median *Z*_summary_ = 10.25 and 9.14 respectively), closely mirroring the Scenario 4 results.

## 7 Discussion

Three aspects of the pooling problem have received little systematic attention in the context of co-expression network analysis: whether pooling disrupts discrete module-level community structure, whether this disruption is detectable from routinely reported global topology metrics, and how prevalent the merged approach is in the published literature. Our results address all three questions directly.

### 7.1 Global topology is resilient to merging; module membership is not

A consistent finding across all four scenarios was that global network properties—modularity and clustering coefficient—were largely maintained in the merged network, while module membership was often substantially reorganised. Edge density showed a more nuanced pattern, dropping substantially in Scenarios 2, 3, and 4, indicating that both bidirectional expression differences and structural divergence independently reduce merged network density, though through different mechanisms: expression differences add symmetric noise to gene–gene correlations without distorting community structure, while structural divergence appears to cause systematic cancellation of population-specific edges, consistent with the leverage effects illustrated in the toy examples.

This dissociation between global and local network properties is an important and underappreciated distinction. A researcher relying on modularity or network density as quality checks might conclude that pooling has not distorted the analysis—yet in Scenario 4, global topology was broadly maintained while only 7 of 14 T1 modules were strongly preserved (*Z*_summary_ *>* 10, median *Z*_summary_ = 9.99; mean across 20 replicates 11.9 ± 1.0, 95% interval 10.2–13.5). Metrics such as modularity or network density, which are routinely reported as quality checks in WGCNA studies, are therefore insufficient diagnostics for detecting the confounding effects of population structure on module membership.

This pattern was directly replicated in empirical data from genetically divergent lake and stream stickleback [Lohman et al., 2017]: merged analysis recovered only 19 modules compared to 26 and 59 in the population-specific networks, with moderate preservation of lake and stream modules in the merged network (median *Z*_summary_ = 10.25 and 9.14 respectively), despite the populations having genuinely divergent co-expression architectures (lake modules in stream network: median *Z*_summary_ = 5.23). It is important to note that Lohman et al. [2017] themselves recognised this risk. As a result, they generated population-specific networks alongside the merged network to verify that module–trait correlations were not confounded by population origin. This independent convergence on the same safeguard suggests the confounding effects induced by pooling data are practically recognised even when they are not formally documented, motivating the explicit guidance in Recommendations 3 and 4.

### 7.2 Structural differences drive module disruption; expression differences compound but do not redirect it

Comparing the four scenarios reveals that network structure differences and mean expression differences affect module organisation through distinct mechanisms. Structural differences (Scenarios 3 and 4) were the primary driver of module disruption, reducing strongly preserved T1 modules from 14/14 in Scenario 1 to 10/14 in Scenario 3 and 7/14 in Scenario 4; expression differences alone (Scenario 2) caused an intermediate reduction to 11/14. When expression differences were combined with structural divergence (Scenario 4), disruption increased modestly but symmetrically (T1: 7/14, T2: 8/14), confirming that expression differences do not introduce systematic directional bias—the merged network fails both populations approximately equally.

The stickleback data corroborate this in the context of Scenario 4: lake and stream fish differ in both mean expression and co-expression architecture, and merged analysis correspondingly failed to recover the population-specific module structure of either group. The fact that disruption was detectable despite small per-group sample sizes (*n* = 17 lake, *n* = 14 stream) suggests our simulation findings represent a lower bound on the risks of pooling at the sample sizes typical of published WGCNA studies.

Consensus WGCNA substantially mitigates these effects. Whereas merged analysis recovered only 7 of 14 T1 modules strongly (median *Z*_summary_ = 9.99), consensus WGCNA recovered all 14 modules from both populations with strong preservation (median *Z* = 24.74 for T1 and 21.80 for T2)—performance comparable to the Scenario 1 baseline. This improvement arises because consensus WGCNA requires co-expression relationships to be present in both populations, filtering out spurious between-population correlations.

Both mean expression differences and co-expression structure differences are empirically well documented in the contexts where WGCNA is most commonly applied [Robinson et al., 2010, Love et al., 2014, Law et al., 2014, Oldham et al., 2006, Horvath and Dong, 2008, Langfelder et al., 2011, Zhang et al., 2018]; the assumption that two biologically distinct groups share the same co-expression architecture is therefore unlikely to hold in most practical applications (see Supplementary Material for a fuller review of the supporting literature).

### 7.3 Prevalence of the merged approach in published WGCNA studies

To situate our simulation findings in the context of current practice, we surveyed 100 randomly selected publications applying WGCNA to multi-group expression data published between 2018 and 2025 (see Supplementary Material). Of the 75 papers with sufficient methodological reporting to permit classification, 59 (78.7%, 95% CI 69.4–87.9%) pooled all samples into a single merged WGCNA pipeline; 11 (14.7%) applied WGCNA separately to each group; and 5 (6.7%) used consensus WGCNA [Langfelder and Horvath, 2008]. No paper used a mixed or unclear approach among those with sufficient reporting. It is concerning, however, that roughly one in seven of the randomly sampled papers did not describe their analyses clearly enough for us to know which approach they took.

The dominance of the merged approach was consistent across biological contexts, spanning human disease studies (tumour versus normal tissue, disease versus healthy controls), agricultural and plant biology (distinct genotypes, treatment conditions), and animal models (pre-defined phenotypic or genetic subgroups). In the majority of merged-approach papers, the disease or group status was used as a downstream trait in module–eigengene correlation analyses after a single network was constructed from all pooled samples—the precise design that our simulations show is most susceptible to the confounding effects of population structure.

A common and concerning pattern in the surveyed literature was the pre-filtering of input genes by differential expression between groups before WGCNA construction, followed by correlation of the resulting modules with the same group variable used for pre-filtering. This design is circular by construction: the modules identified are enriched for genes that differ between groups—the same genes a simpler differential expression analysis would have identified—so the network framework adds little independent biological information [Langfelder and Horvath, 2020]. For example, Chai et al. [2022] used consensus WGCNA to compare co-expression networks in gray and white matter lesions in multiple sclerosis, concluding high overall similarity based on eigengene network density (D = 0.79) and subsequently focusing on modules enriched for differentially expressed genes– - an approach that may overlook substantial local reorganisation and introduce circularity. This circularity compounds the distortions documented in our simulations: not only does the merged network misrepresent the co-expression architecture of both populations, but the pre-filtering step further ensures that identified modules recapitulate differential expression rather than genuine co-regulation.

Among the 11 papers using population-specific networks, the most common approach was a split-then-intersect design that identifies hub genes shared across networks via Venn diagram overlap, without formally testing whether module *structures* are preserved—a weaker standard than the modulePreservation framework [Langfelder et al., 2011]. The 6.7% prevalence of consensus WGCNA is a small but encouraging minority; its low adoption likely reflects greater computational complexity and less prominent documentation, though sample-size constraints should not be used as justification for pooling (see Supplementary Material for further discussion).

### 7.4 Limitations

Several limitations of the present study should be noted.First, the simulations used a fixed network size of 500 genes and 150 samples per population. Second, the network modification procedure (randomly rewiring 30% of edges) does not correspond to a specific biological mechanism and may not capture the structural differences observed in real populations arising from genetic variation, developmental stage, or tissue type.

A sensitivity analysis spanning network sizes of 250, 500, and 1,000 genes and rewiring rates of 10%, 30%, and 50% confirmed that the qualitative pattern of results is robust to both parameter choices (Figure S7). Module disruption increased with rewiring rate at all network sizes, and Scenario 4 showed consistently similar or lower preservation than Scenario 3 across the full grid. Disruption was more severe in smaller networks, suggesting the risks of pooling are most acute for smaller gene sets. Per-group sample size was held constant at *n* = 150 and was not varied in the sensitivity grid; at smaller per-group *n*, correlation estimates become increasingly unreliable [Langfelder and Horvath, 2020] and *Z*_summary_ statistics are attenuated [Langfelder et al., 2011].

Third, the block-diagonal covariance assigns uniform within-community correlation (*ρ* = 0.85) and zero between-community correlation, which does not capture the graded correlation decay or hub-driven within-community heterogeneity present in real co-expression data. Fourth, because we used a fixed *β* = 6 across all scenarios, our results may not generalise to settings where the optimal soft-thresholding differs substantially between populations. Finally, only pairwise population comparisons were considered; the behaviour under merging of three or more populations with heterogeneous structures remains an open question (see Supplementary Material).

Our literature survey carries its own limitations. It is based on a random sample of 100 papers (2018–2025), of which 25 were excluded for insufficient methodological reporting, so the 78.7% prevalence applies only to papers with adequate reporting. The search query requires multi-group language in the title or abstract, likely under-representing studies that describe their multi-group design only in the Methods. Full survey methodology and the complete coding sheet are provided in the Supplementary Material.

Together, these simulation and empirical findings carry direct practical consequences for the large fraction of published WGCNA studies that pool samples from biologically distinct groups, and motivate the concrete recommendations that follow.

## 8 Recommendations for Practitioners

Based on the findings of this study, we offer the following practical recommendations for researchers applying WGCNA to datasets comprising samples from multiple populations, genotypes, treatments, or experimental conditions.

### 8.1 Test for between-group differences before merging

Before pooling samples from multiple groups into a single WGCNA pipeline, researchers should assess whether the groups differ in mean expression levels and, where possible, in co-expression structure. Differential expression analysis (e.g. using DESeq2 [Love et al., 2014] or edgeR [Robinson et al., 2010]) can identify genes with significantly different mean expression between groups. A large number of differentially expressed genes is a strong indication that pooling will induce spurious correlations in the merged network, as demonstrated in Toy Example 2 and Scenarios 2 and 4. Similarly, a preliminary comparison of pairwise gene–gene correlations between groups—even for a subset of genes—can reveal whether co-expression structure is conserved across groups before committing to a merged analysis. The importance of assessing between-group differences before network construction is not unique to transcriptomics: NetCoMi [Peschel et al., 2021], developed for microbiome association networks, explicitly builds between-group differential testing into its recommended workflow as a precondition for network comparison, reflecting a broader consensus that group differences must be characterised before any network inference is attempted.

### 8.2 Check for confounders before and after WGCNA

Expression data should be checked for batch effects and population structure using principal component analysis or similar methods [Leek et al., 2024] before WGCNA is applied. Variance stabilisation and normalisation methods that account for group differences in mean expression (e.g. variance-stabilising transformation in DESeq2) can also reduce the leverage effects described in the toy examples, though they do not eliminate them entirely. The term *leverage* here has a precise regression-theoretic meaning: observations far from the mean of the predictor variable exert disproportionate influence on the estimated regression coefficient. In the pooled WGCNA setting, samples from different populations occupy distinct regions of gene expression space, separated by ***δ*** = ***µ***_1_ − ***µ***_2_, placing both population clouds far from the grand mean of the merged dataset. The between-group mean difference therefore acts as a high-leverage configuration: the slope of the merged gene–gene regression is dominated by the between-population term *δ_A_δ_B_* rather than by the within-population covariance structure, as shown in Equation 1. Variance stabilisation reduces ∥***δ***∥ by bringing population means closer together, thereby reducing this leverage—but unless mean differences are eliminated entirely, the bias term 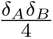 remains non-zero and the merged correlation continues to reflect between-population structure to some degree.

If pooling is unavoidable (e.g. due to limited sample sizes within individual groups), population membership should be included as a covariate in downstream analyses of module–trait relationships. Including population as a covariate removes the between-group contrast from the module–eigengene association estimates, partially mitigating the confounding documented in our simulations, though it does not correct the upstream distortion of the correlation matrix from which modules were originally derived.

### 8.3 Apply WGCNA separately to each group before merging

Where sample sizes permit, we recommend running WGCNA independently on each group and comparing the resulting modules before any merged analysis is attempted. Population-specific modules provide a baseline against which the modules from a merged network can be evaluated. If modules identified in the individual groups are largely recapitulated in the merged network (assessed using Fisher’s exact test or WGCNA’s modulePreservation function), the merged analysis can be interpreted with greater confidence. However, if population-specific modules are substantially reorganised in the merged network, the merged modules should be interpreted with caution.

This diagnostic step is not merely precautionary—it has practical precedent: Lohman et al. [2017] independently adopted this approach in their stickleback analysis, generating population-specific networks alongside the merged network to verify that module–trait correlations were not confounded by population origin. As reported in the Discussion, our reanalysis confirmed that merged analysis collapsed the module structure relative to the population-specific networks, illustrating the kind of disruption that a merged-only analysis would have missed. For a more granular alternative, CoDiNA [Morselli Gysi et al., 2020] classifies edges and nodes as common, group-specific, or different across networks, complementing module overlap analysis.

### 8.4 Use consensus WGCNA for multi-group analyses

For datasets from multiple groups that are intended to yield a single, shared set of modules, we recommend consensus WGCNA [Langfelder and Horvath, 2008] rather than a simple merging of samples. Consensus WGCNA identifies modules that are consistently present across groups by constructing a consensus topological overlap matrix from the group-specific TOMs, effectively requiring that co-expression relationships be preserved across groups for genes to be assigned to the same module. This approach is more robust to between-group differences in mean expression and network structure than pooling, because it down-weights gene–gene relationships that are present in only one group.

Our simulations provide direct empirical support for this recommendation: as reported above, consensus WGCNA applied to the Scenario 4 data recovered all 14 modules from both populations with strong preservation, compared to only 7 of 14 under merged analysis. By requiring co-expression relationships to be present in both populations simultaneously, consensus WGCNA avoids the spurious correlations introduced by pooling and recovers modules that genuinely represent the shared transcriptional architecture of both groups.

However, consensus modules may overlook biologically interesting differences in network structure among populations, since modules specific to one group are by design excluded from the consensus. Researchers whose primary interest is in population-specific co-expression patterns should therefore apply WGCNA separately to each group (Recommendation 3) rather than relying on consensus analysis alone.

### 8.5 Use module preservation statistics as a diagnostic

After running WGCNA on a merged dataset, WGCNA’s `modulePreservation` function [Langfelder et al., 2011] provides a routine check on whether merged modules are preserved in the individual group networks. A *Z*_summary_ *<* 2 for a module indicates that it is not preserved in the reference network and therefore likely reflects a statistical artefact of the pooling process; such modules should not be interpreted biologically without further validation. As shown in Scenarios 3 and 4, even modules with moderate preservation scores (*Z*_summary_ between 2 and 10) can mask substantially reorganised gene membership relative to the corresponding modules in the individual networks.

Computing preservation using each constituent population as the reference network in turn—not only the population of primary biological interest—guards against missing asymmetric disruption. In most multi-group comparisons with standard normalisation, differential expression between groups is bidirectional and disruption will be approximately symmetric across populations, as demonstrated in Scenario 4. Computing preservation in both directions confirms this symmetry and ensures that disruption affecting either population is not overlooked.

The T1-versus-T2 comparison provides a further useful diagnostic: when individual-versus-merged preservation is substantially lower than individual-versus-individual preservation, confounding is likely attributable to the pooling procedure rather than to inherent biological differences. As a practical heuristic, a drop of more than 5 units in median *Z*_summary_ between the individual-versus-individual and individual-versus-merged comparisons suggests that the merged network is distorted by pooling (as in Scenario 2, where the T1-versus-T2 median was 21.0 versus the merged median of 15.2). However, when individual-versus-individual preservation is itself low (*<* 10, as in Scenarios 3 and 4), low merged preservation more likely reflects genuine biological divergence, and the two sources cannot be cleanly separated without additional validation.

Finally, while modulePreservation summarises preservation at the module level, CoDiNA [Morselli Gysi et al., 2020] complements it at the edge and node level, pinpointing which gene–gene relationships differ between the merged and group-specific networks.

### 8.6 Report network diagnostics at the module level, not just globally

Our results demonstrate that commonly reported global network metrics can be misleading indicators of merged network quality. Modularity and clustering coefficient remained stable across all scenarios even when module membership was substantially disrupted, providing no assurance that merged modules are biologically meaningful. Edge density dropped substantially in Scenarios 2–4 but is non-specific: it cannot distinguish expression-driven density loss (Scenario 2, mostly preserved modules) from structure-driven module disruption (Scenarios 3–4). Researchers should therefore report module-level diagnostics—module size distributions, module density, and inter-network overlap statistics—rather than relying on global topology metrics to validate WGCNA results.

## Data Availability Statement

Our simulation code is available on GitHub (https://github.com/rogini98/wgcna-pooling-bias). All empirical data used from Lohman et al. [2017] are available from the Dryad repository https://doi.org/10.5061/dryad.mk8ns. All other data were simulated.

## Authors’ contributions

This work started while RR was in TE-R’s research group. Both RR and DIB designed this study. RR contributed to the code for both the simulations and the analysis. RR wrote the initial draft of the paper. DIB and TE-R contributed to the editing of the current manuscript. All authors contributed to the framing of the manuscript.

## Acknowledgements

We used Claude Code (April 2026 version) to proofread the manuscript and implemented some of its suggestions.

## Funding

RR, TE-R, and DIB acknowledge funding from the National Science Foundation (FAIN-2133740). RR acknowledges the support of The Roux Institute and the Harold Alfond Foundation.

## Conflicts of interest

All authors declare that they have no conflicts of interest.

## Supplementary Material

### Additional methodological observations from the literature survey

#### Split-then-intersect designs

Among the 11 papers in our survey that applied WGCNA separately per group, the most common approach was a split-then-intersect design, in which independent networks were built for each dataset or cohort and hub genes common to both were identified via Venn diagram overlap. While this approach avoids the direct confounding of pooling, the intersection step does not formally test whether the underlying module *structures* are preserved across populations—it selects genes that appear in corresponding modules in both networks without quantifying the statistical evidence for structural similarity. The most methodologically rigorous papers in our survey used the `modulePreservation` framework [Langfelder et al., 2011] to formally test which modules from a training network replicate in independent validation cohorts, retaining only preserved modules for downstream analysis. We recommend this approach as the minimum standard for multi-cohort WGCNA studies.

The 6.7% prevalence of consensus WGCNA represents a small but encouraging minority. Consensus WGCNA [Langfelder and Horvath, 2008] explicitly acknowledges the multi-population structure of the data and identifies modules that are robust across groups, directly addressing the problem our simulations document. Its low adoption likely reflects greater computational complexity, less prominent documentation relative to the standard single-network workflow, and the absence of widely adopted guidelines recommending its use for multi-group data. Researchers may also avoid this approach if they have low per-group sample sizes. The WGCNA developers recommend a minimum of 15 samples per network, and ideally at least 20 [Langfelder and Horvath, 2020]. Such constraints should not be taken as justification for pooling, however; an underpowered within-group network is preferable to a merged network that misrepresents the co-expression architecture of both populations.

#### Empirical context: ubiquity of between-group differences

Both mean expression differences and co-expression structure differences are empirically well documented in the contexts where WGCNA is most commonly applied. Differences in mean expression between biological groups are nearly ubiquitous in transcriptomic studies: comparisons of tumour versus normal tissue, distinct genotypes, developmental stages, or treatment conditions routinely identify hundreds to thousands of differentially expressed genes, often with large effect sizes [Robinson et al., 2010, Love et al., 2014, Law et al., 2014]. Differences in co-expression structure between groups have received less systematic attention, but the evidence where it has been examined is consistent: co-expression networks constructed separately from distinct biological groups frequently differ in module composition, hub gene identity, and inter-module connectivity [Oldham et al., 2006, Horvath and Dong, 2008, Langfelder et al., 2011, Zhang et al., 2018, Liu et al., 2019, Zhou et al., 2021]. The assumption that two biologically distinct groups share the same co-expression architecture—the implicit assumption of any merged WGCNA pipeline—is therefore unlikely to hold in most practical applications.

#### Behaviour under merging of three or more populations

Only pairwise population comparisons were considered in the main simulations. The behaviour of WGCNA under merging of three or more populations with heterogeneous structures remains an open question. We expect that adding more populations creates additional opportunities for among-group covariance differences, mean-expression shifts, and covariance among population means—all of which could further skew the merged network beyond what our two-population simulations document.

### Systematic review of WGCNA multi-group analytical practice

#### Search strategy

We searched PubMed on April 8, 2026 using the `rentrez` R package [Winter, 2017] with the following query:

~~~
WGCNA[tiab] AND ("multiple populations"[tiab] OR "two populations"[tiab] OR "multiple
groups"[tiab] OR "two groups"[tiab] OR "multiple genotypes"[tiab] OR "two genotypes"[tiab]
OR "case and control"[tiab] OR "cases and controls"[tiab] OR "disease and control"[tiab]
OR "treated and untreated"[tiab] OR "multiple tissues"[tiab] OR "multiple timepoints"[tiab]
OR "multiple breeds"[tiab] OR "multiple cohorts"[tiab] OR "multiple datasets"[tiab]
OR "multiple conditions"[tiab] OR "multiple species"[tiab] OR "two conditions"[tiab]
OR "two tissues"[tiab] OR "two cohorts"[tiab]) AND ("2018"[pdat] : "2025"[pdat])
~~~

The `[tiab]` field tag restricts matching to titles and abstracts; the date filter restricts results to papers published between 1 January 2018 and 31 December 2025. This query was designed to identify papers where the presence of multiple biological groups was signalled in the title or abstract. The query requires multi-group language to appear in the title or abstract, which may under-represent studies where the multi-group structure of the data is described only in the Methods section. Our survey should therefore be interpreted as characterising WGCNA practice in studies that explicitly foreground their multi-group design, rather than as a complete census of all multi-group WGCNA applications. A random sample of 100 was drawn from these records using a fixed random seed (seed = 42) in R for reproducibility.

#### Text retrieval

For each sampled paper, metadata (title, authors, journal, year, DOI) and the abstract were retrieved from PubMed via XML using `entrez fetch`. For papers with a PubMed Central (PMC) identifier, the full-text XML was additionally retrieved from the PMC database and parsed to extract the Methods and Results sections. Section extraction targeted `<sec>` elements whose `<title>` contained any of the following terms: *method*, *material*, *result*, *statistical*, *data analysis*, *bioinformatic*, *network analysis*, *wgcna*, *co-expression*. For papers without PMC full text, only the abstract was used and the paper was automatically flagged for manual review. Of the 100 sampled papers, 90 had Methods/Results text available from PMC and 10 were coded from abstract only.

#### Inclusion and exclusion criteria

Papers were included if two criteria were both satisfied: (i) WGCNA was applied empirically (not merely cited), evidenced by the presence of at least one of the following terms in the title or retrieved text: *“WGCNA was performed/applied/used/conducted/run”*, *“weighted gene co-expression network”*, *“soft-thresholding power”*, *“blockwiseModules”*, *“module eigengene”*, *“TOM-based dissimilarity”*, *“dynamic tree cut”*, or *“topological overlap matrix”*; and (ii) the dataset comprised two or more distinct biological groups, evidenced by at least one of the following patterns in the title or retrieved text: *two/multiple groups*, *populations*, *conditions*, *tissues*, *cohorts*, *genotypes*, *breeds*, or *species*; *case(s) and control(s)*; *disease versus control/healthy/normal* ; *patient versus healthy/control* ; *wild-type versus mutant* ; *tumour versus normal/adjacent* ; *pre/post treatment* ; or *comparison between groups/populations/conditions*. Papers were additionally excluded if WGCNA was applied to non-transcriptomic data (e.g. metabolomics or miRNA abundance matrices), if the WGCNA design could not be determined from available text, or if the biological groups present did not constitute distinct pre-defined populations in the sense relevant to this study (e.g. data-derived subgroups identified post hoc within a single homogeneous population). Of the 100 sampled papers, 75 met both inclusion criteria and 25 were excluded.

#### Coding scheme

Each included paper was assigned one of three codes based on keyword detection in the title and retrieved text (Methods/Results where available, abstract otherwise):

1. *Merged* —all samples from all groups were combined into a single expression matrix and WGCNA was run once on the pooled data, with group membership subsequently correlated with module eigengenes as a trait. Detected by phrases such as: *“all samples were combined/pooled/merged”*, *“WGCNA was performed on all/the combined/pooled data”*, *“module–trait correlation/association”*, *“module eigengenes were correlated with disease/treatment/-group”*, *“modules significantly associated with clinical/disease/prognosis”*, or *“differentially expressed genes between [groups] were selected as input for WGCNA”*.
2. *Split* —WGCNA was applied independently to each group, yielding group-specific networks and module sets, which were then compared across groups. Detected by phrases such as: *“WGCNA was performed separately/independently for each group/dataset/cohort”*, *“separate network for each”*, *“modulePreservation”*, *“module preservation between/across”*, *“population-specific network/module”*, or *“Venn diagram to identify common/shared modules/hub genes across datasets”*.
3. *Consensus*—consensus WGCNA [Langfelder and Horvath, 2008] was used to identify modules consistent across groups. Detected by: *“consensus WGCNA/network/module/analysis”*, *“blockwiseConsensusModules”*, *“consensus topological overlap”*, or *“shared modules across”*.

The coding priority was: Consensus *>* Split *>* Merged. Papers coded from abstract only were automatically flagged for manual review regardless of their automated code, since WGCNA methodology is rarely described in sufficient detail in abstracts. Papers with mixed or conflicting signals were also flagged and manually reviewed by RR; the Methods section of the full paper was read where accessible and a final code assigned based on explicit description of whether samples were pooled, separated, or processed using consensus WGCNA. Of the 100 sampled papers, 76 required manual review. Manual examination resolved every ambiguous case to one of Codes 1–3, with the remainder excluded.

#### Coding reliability

All manual review was conducted by a single coder (RR). The absence of independent double-coding is acknowledged as a limitation; however, the coding scheme was applied to explicit methodological language—the presence or absence of pooling, separate networks, or consensus modules—rather than subjective interpretation. Borderline cases and the evidence phrases used to justify each code are documented in the coding sheet (Supplementary Table).

#### Statistical analysis

Proportions are reported with 95% confidence intervals computed using the normal approximation:

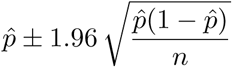

where *p̂* is the observed proportion and *n* = 75 is the number of included papers. For small proportions (e.g. *p̂* ≈ 0.067), the normal approximation is on the boundary of its validity range; results should be interpreted with this in mind. The complete coding sheet, including PMIDs, extracted evidence phrases, and final codes for all 100 sampled papers, is provided in Supplementary Table. The R code implementing the full pipeline is available at https://github.com/rogini98/wgcna-pooling-bias.

**Table S1:** Summary of WGCNA multi-group analytical practice. Results from a random sample of 100 publications (2018–2025), of which 75 met inclusion criteria. Confidence intervals computed using the normal approximation 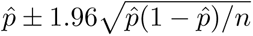, where *n* = 75.

| Classification | *N* | % | 95% CI lower | 95% CI upper |
| --- | --- | --- | --- | --- |
| Merged | 59 | 78.7% | 69.4% | 87.9% |
| Split | 11 | 14.7% | 6.7% | 22.7% |
| Consensus | 5 | 6.7% | 1.0% | 12.3% |
| **Total included** | **75** | **100.0%** |  |  |
| *Excluded* | *25* |  |  |  |

**Table S2:**
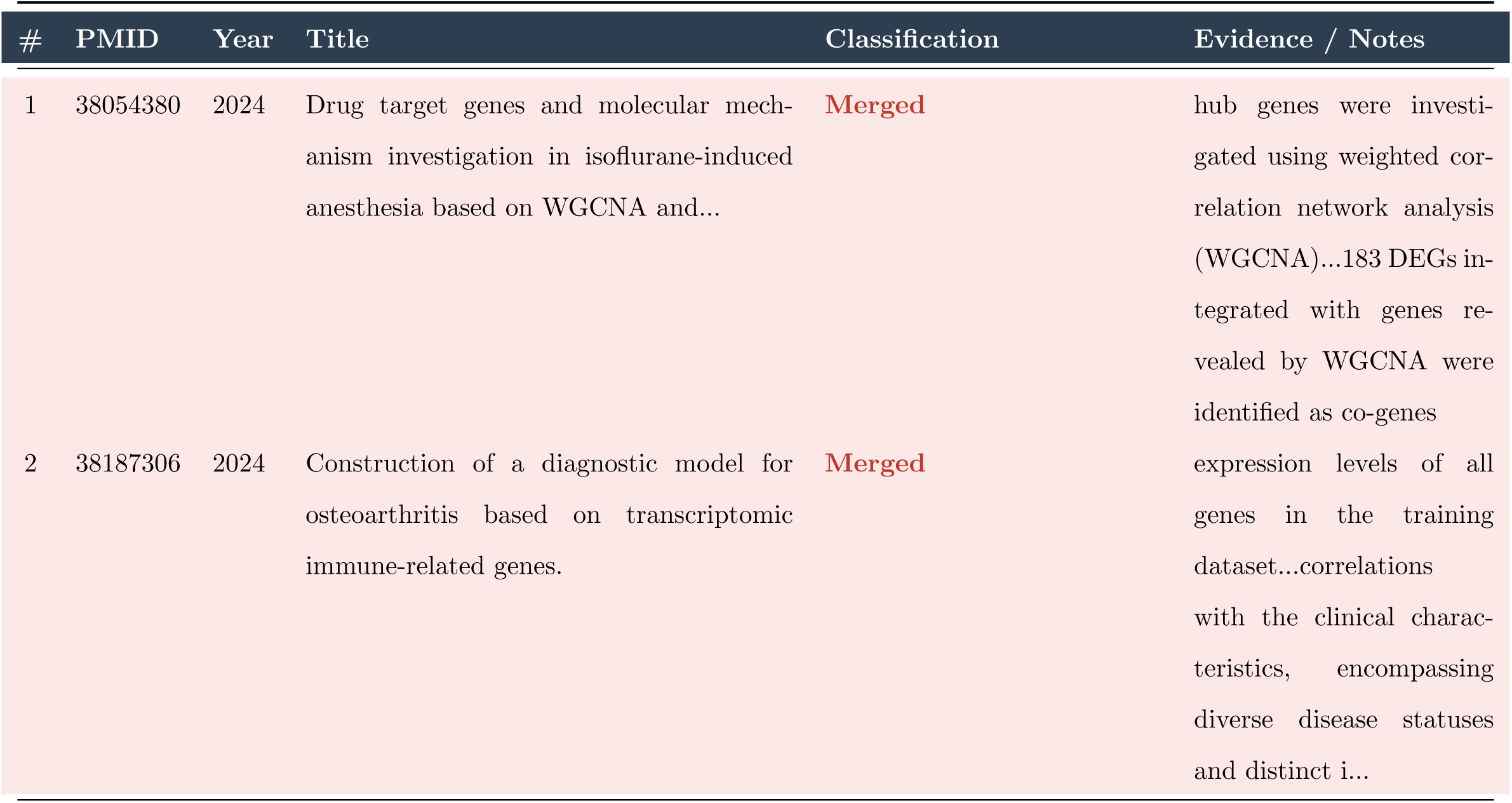
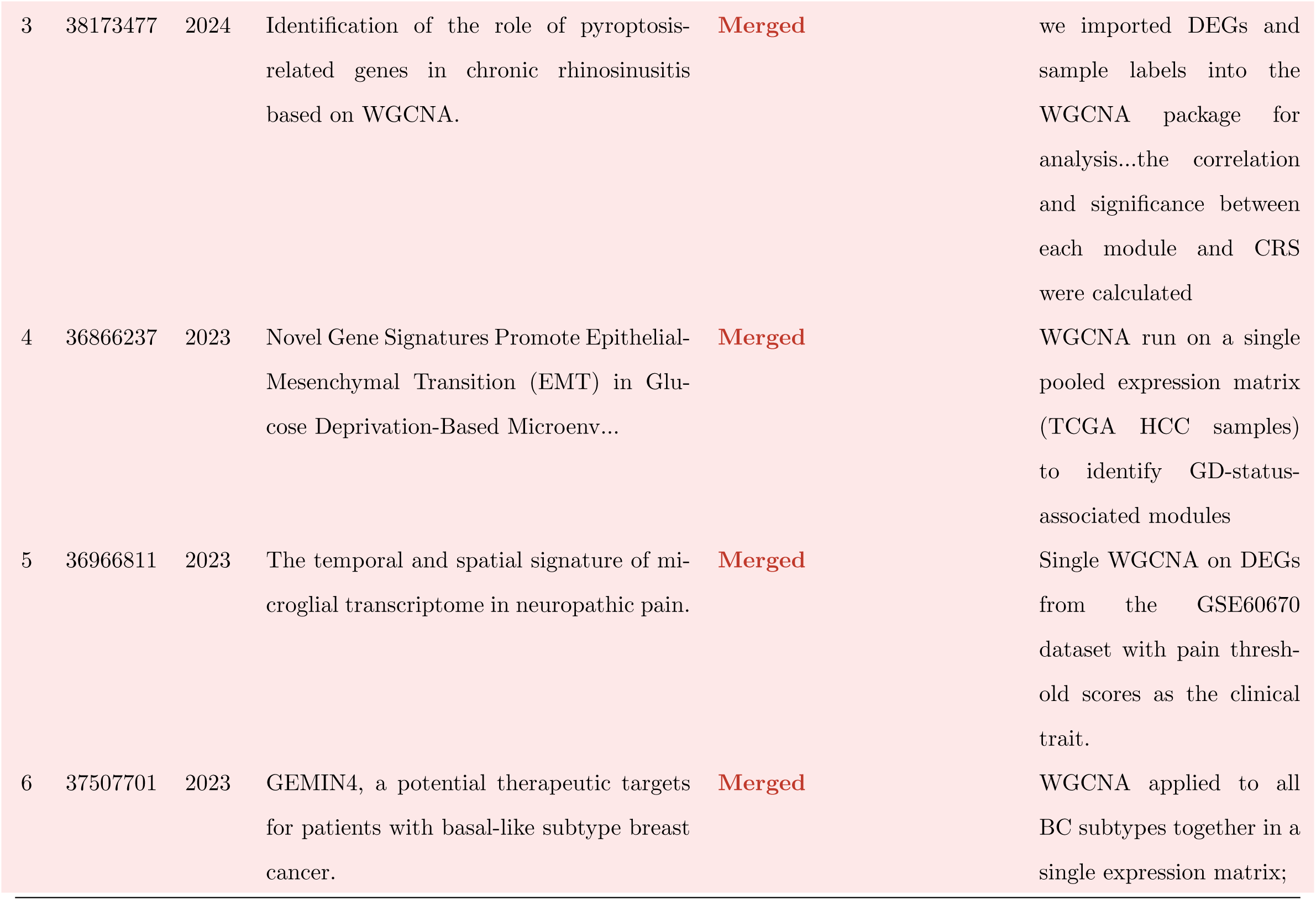
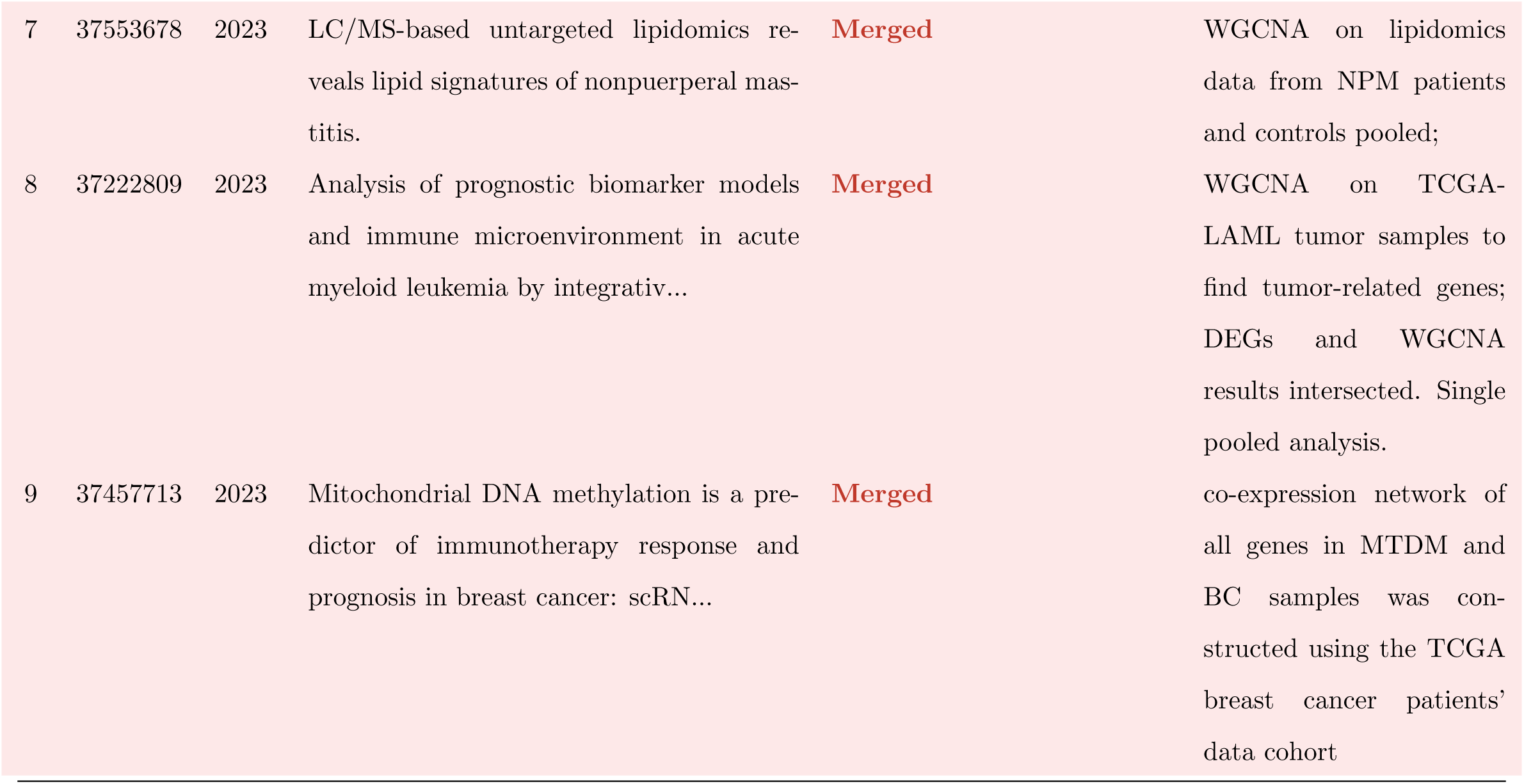
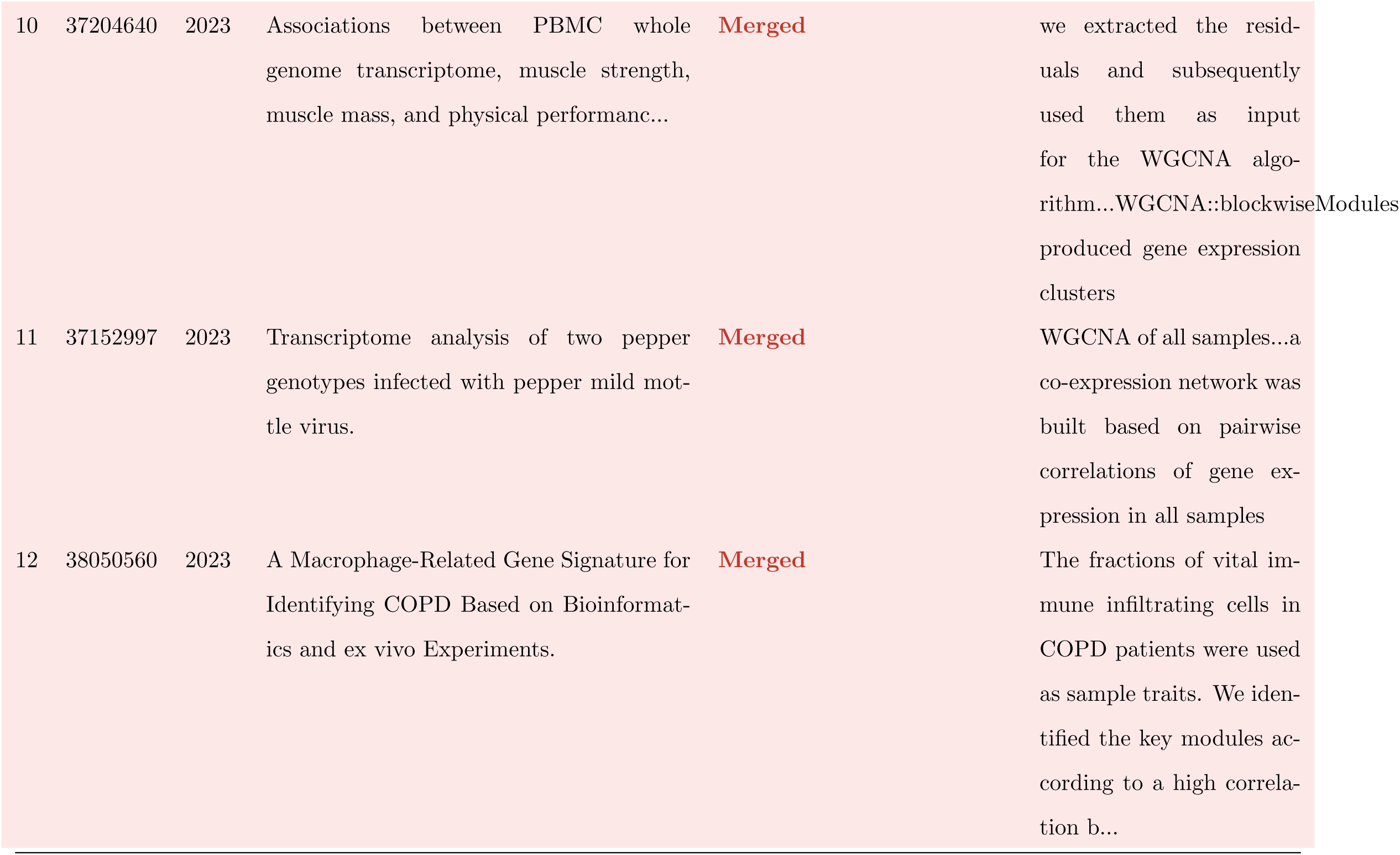
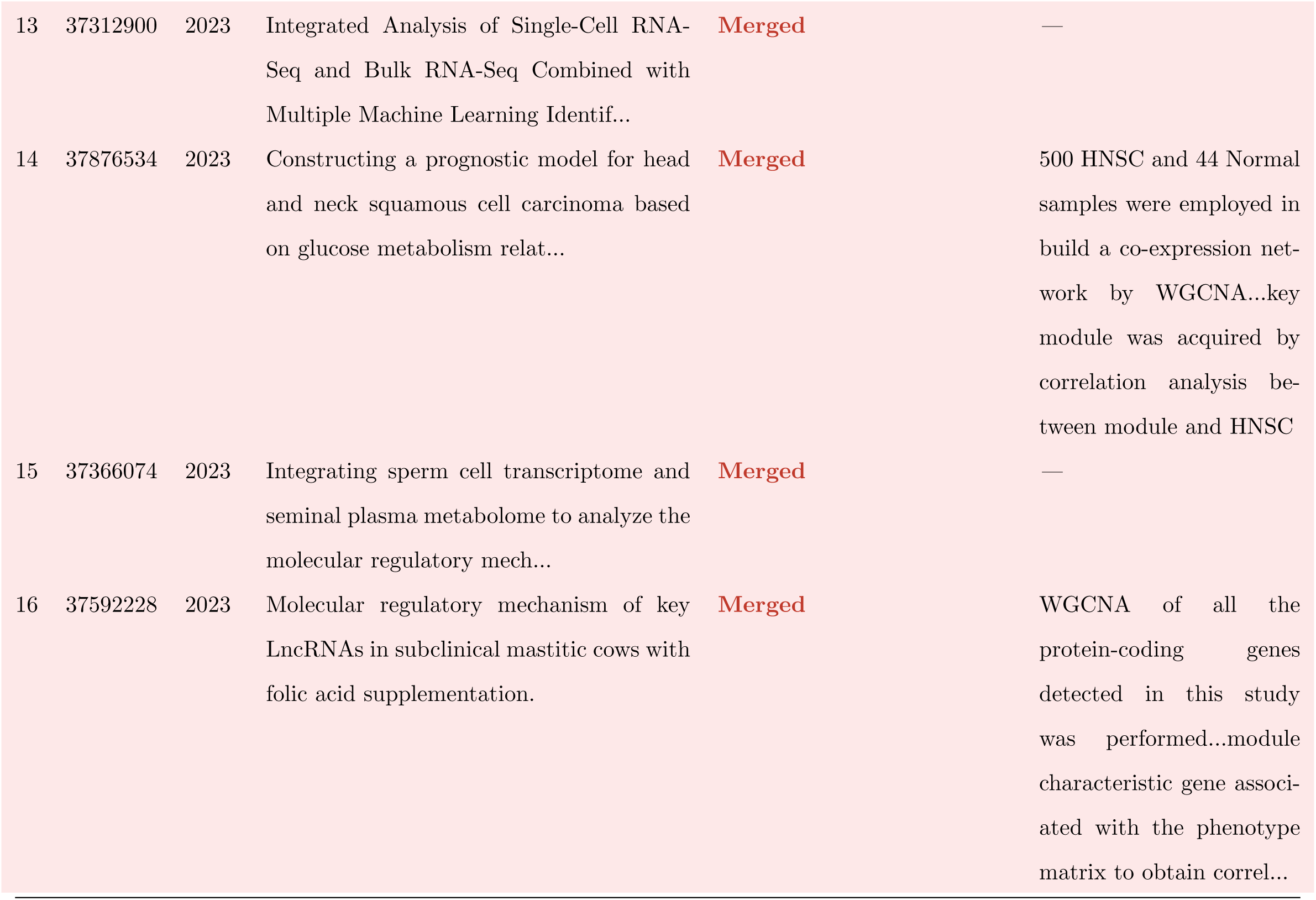
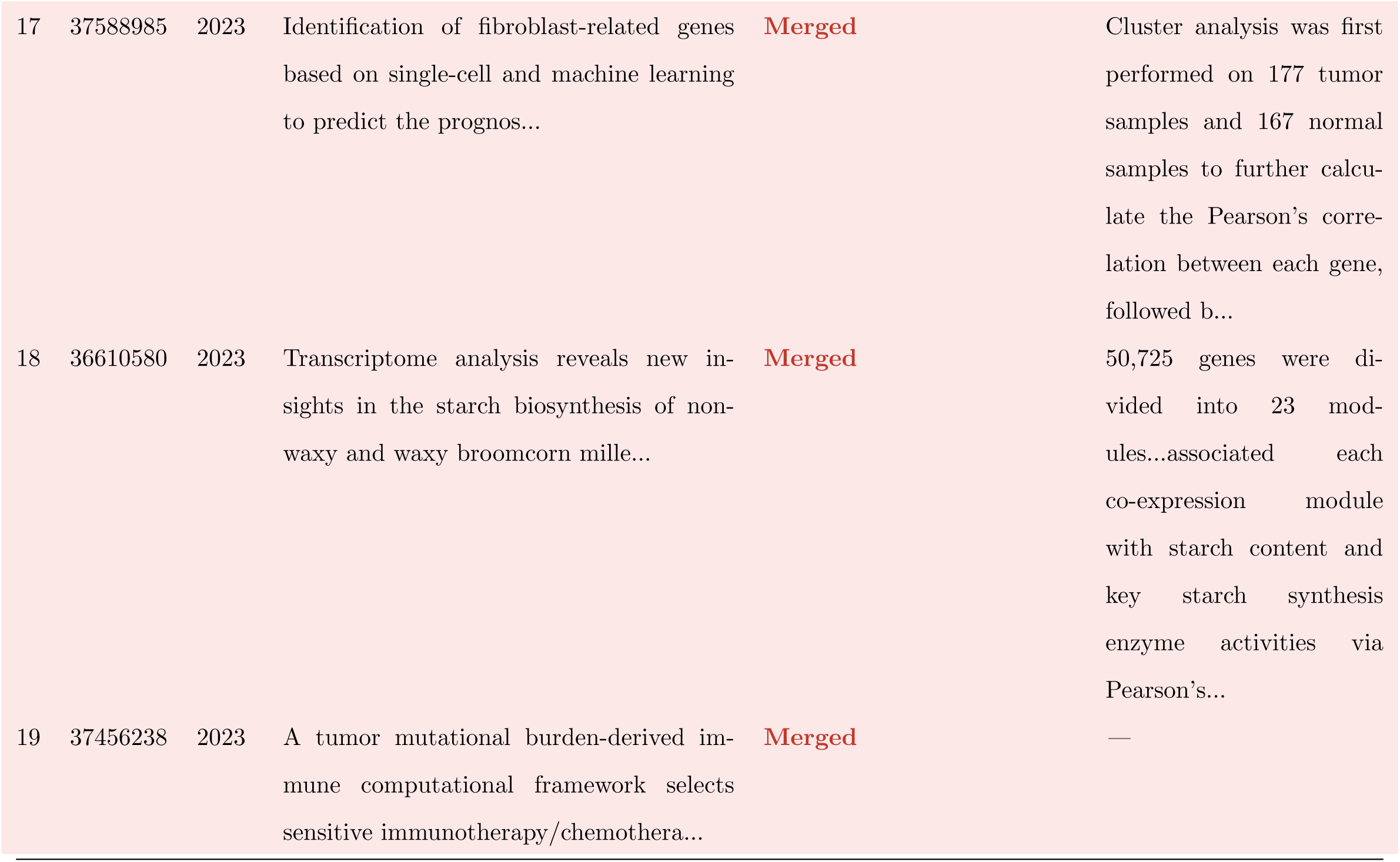
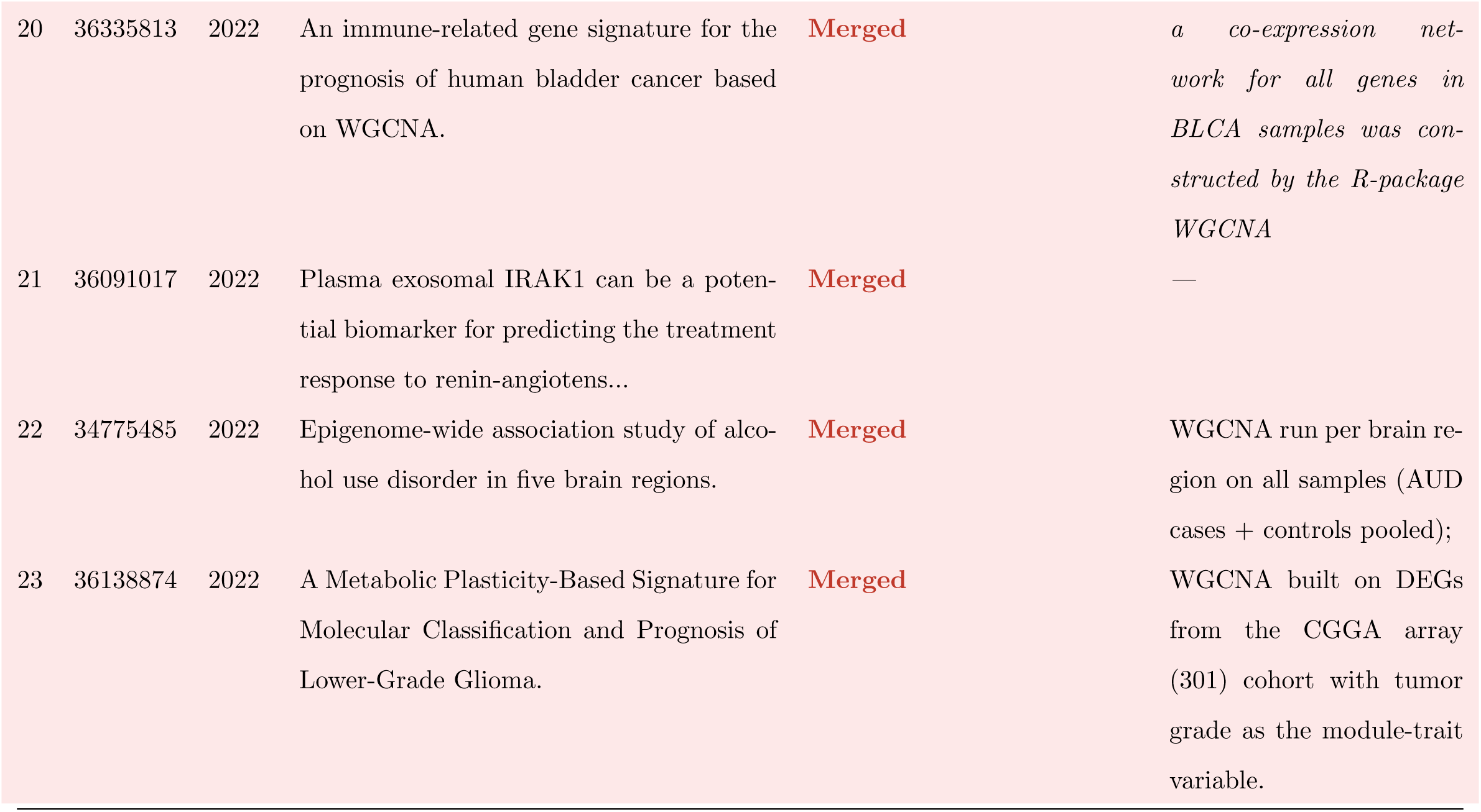
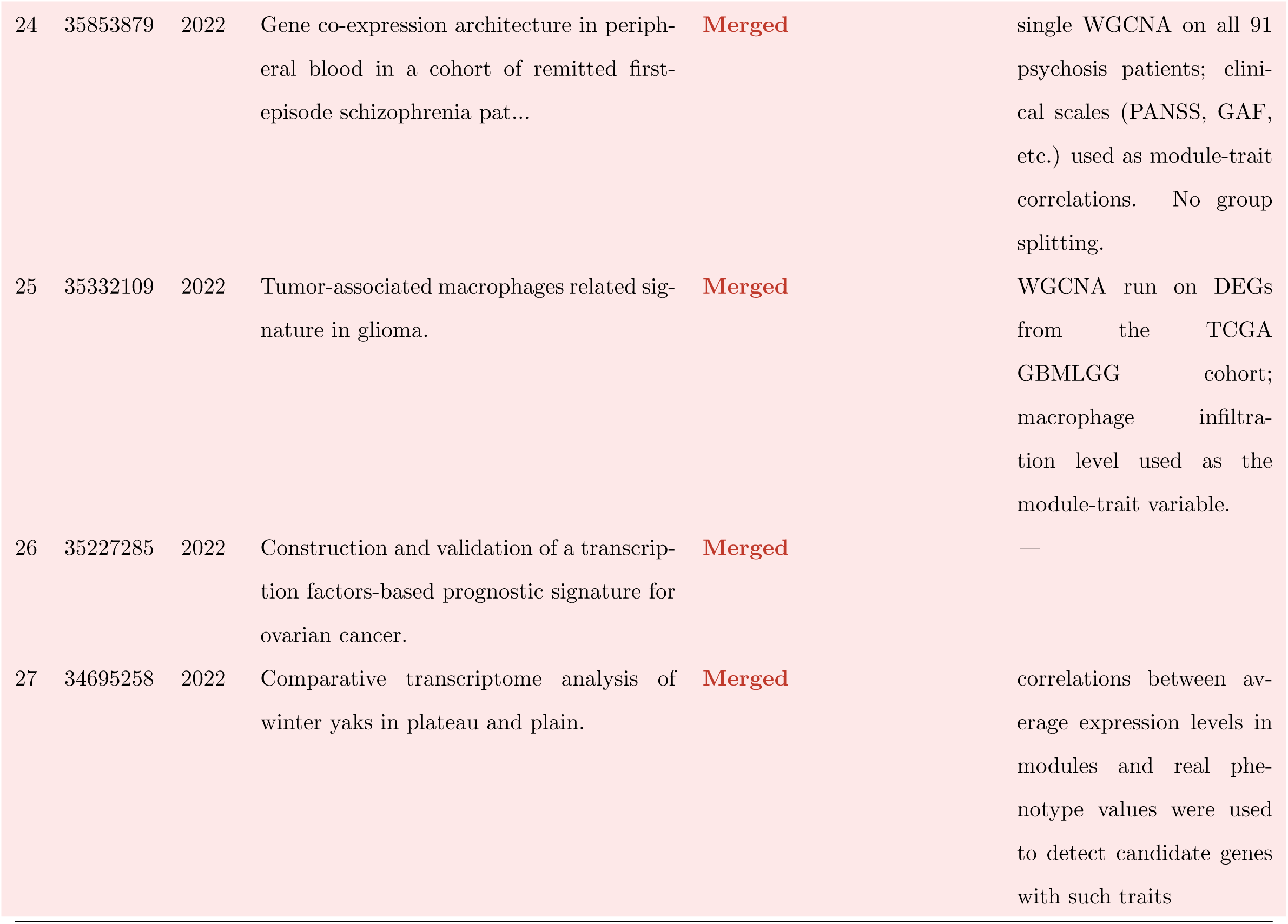
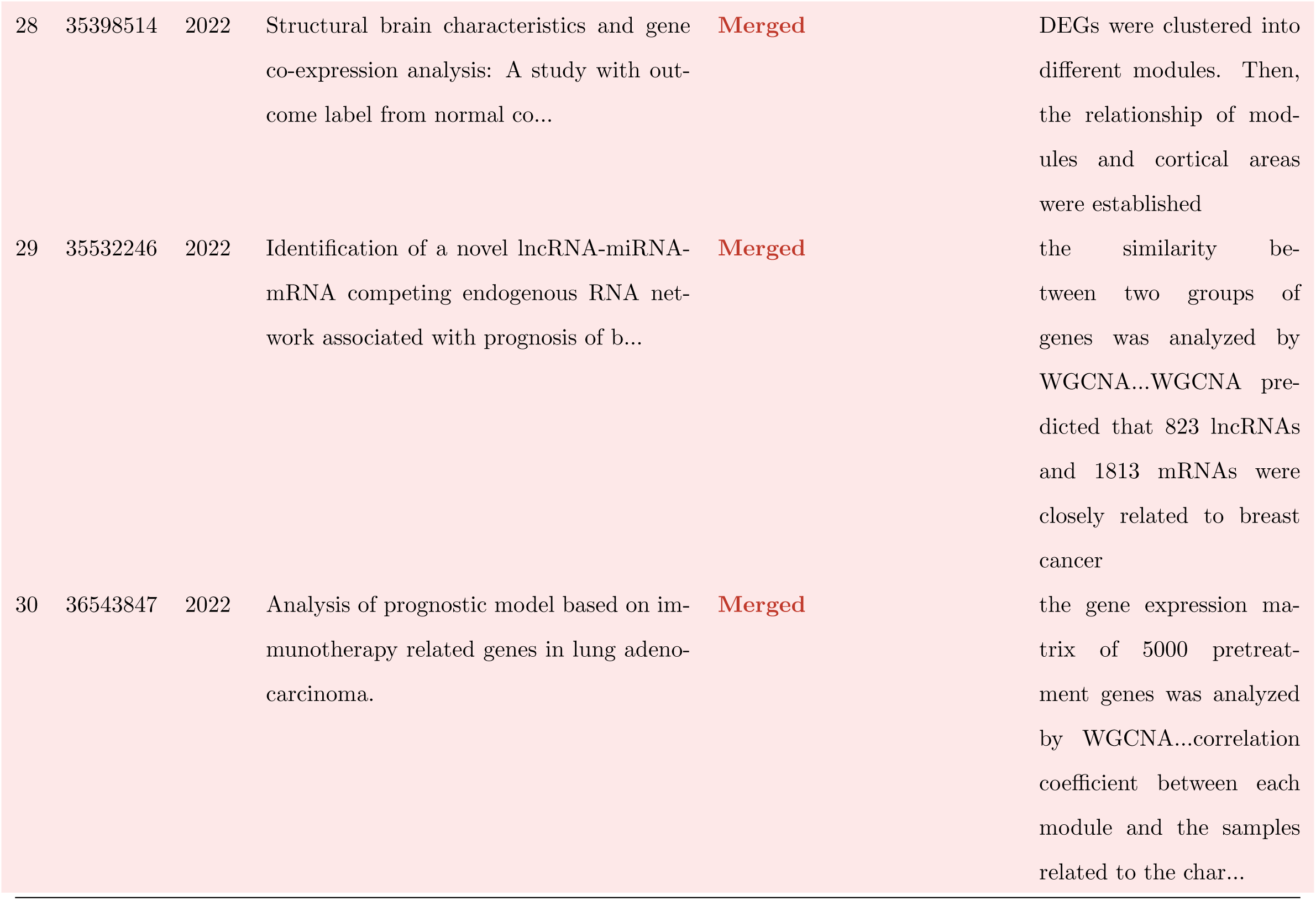
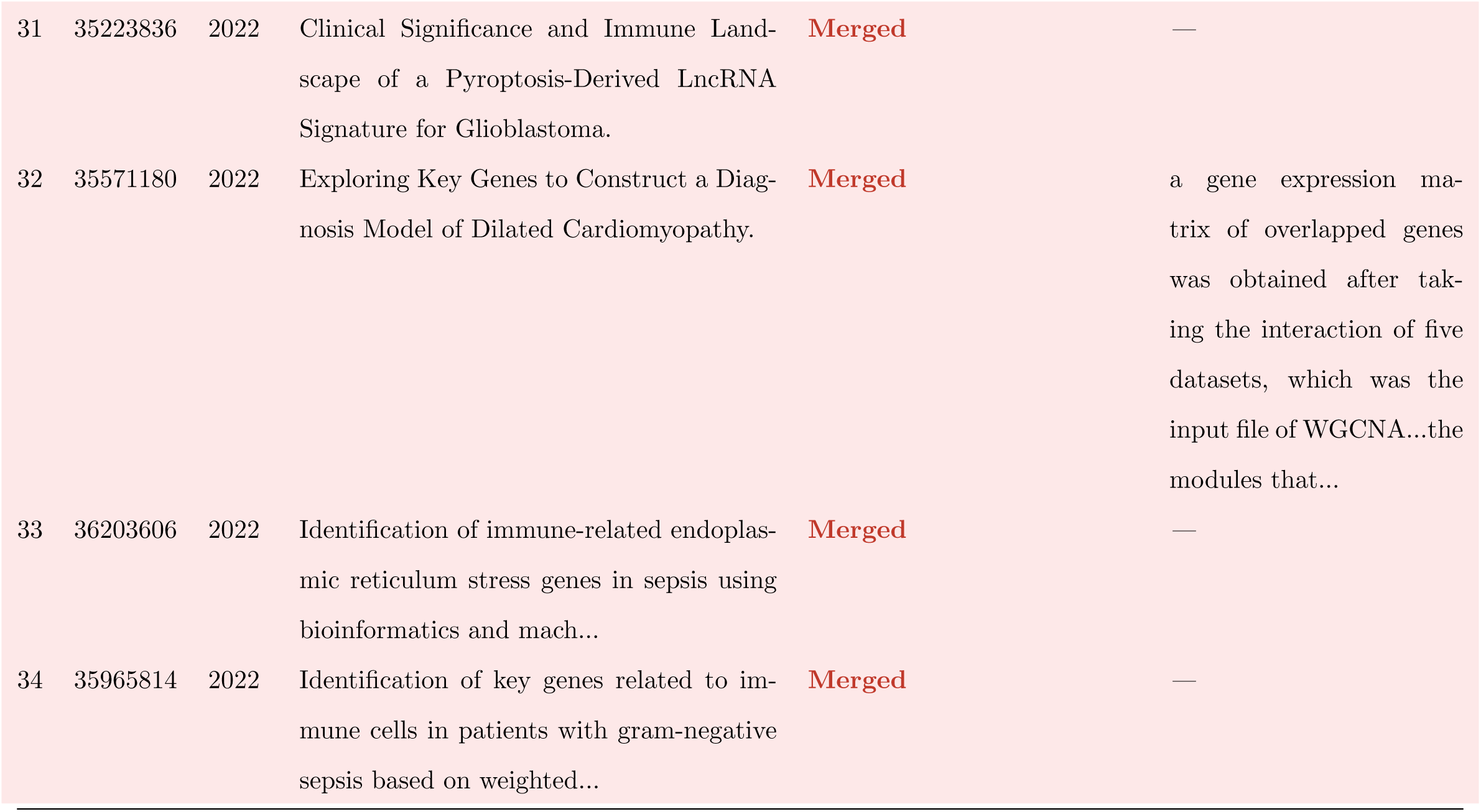
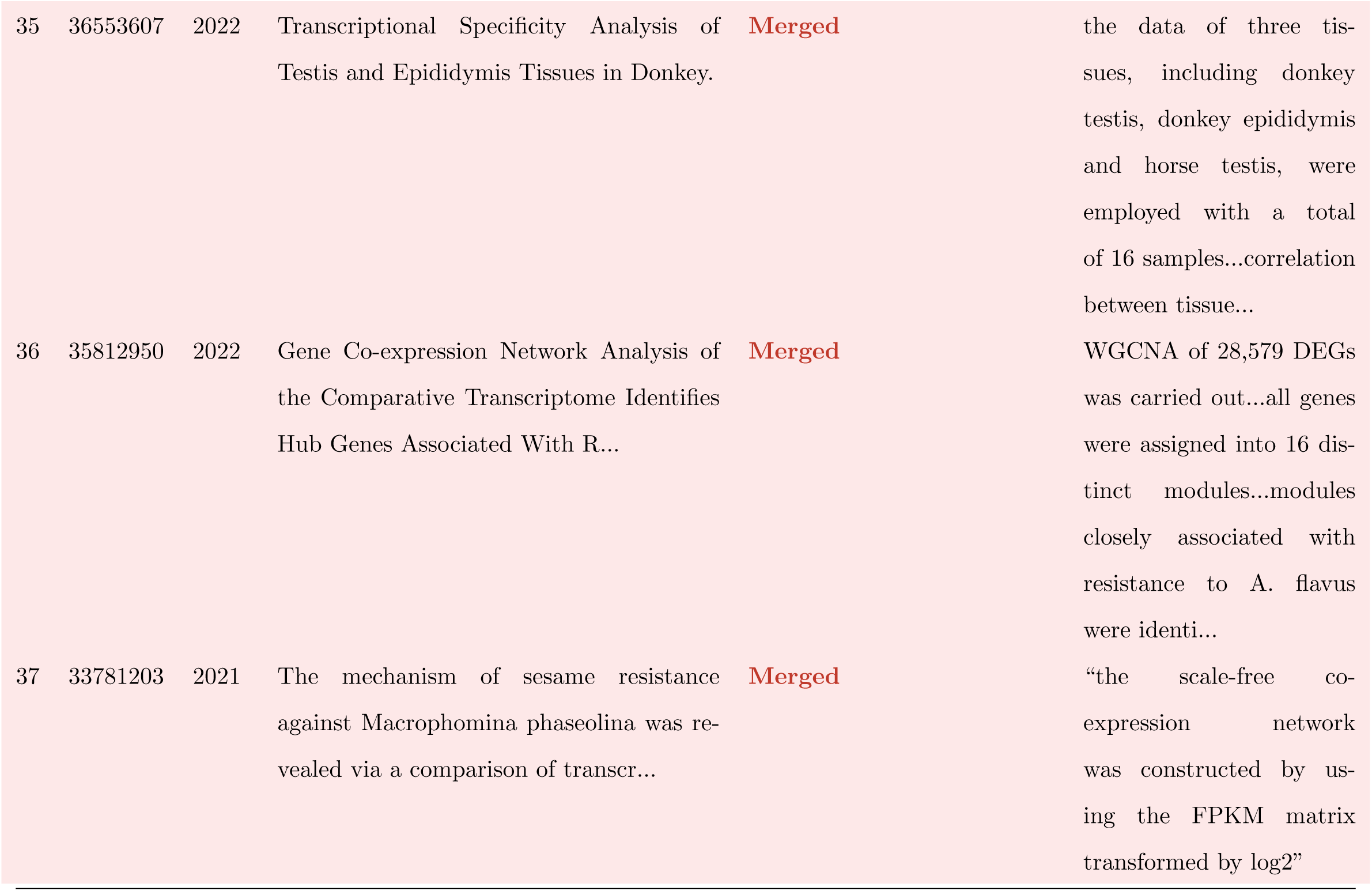
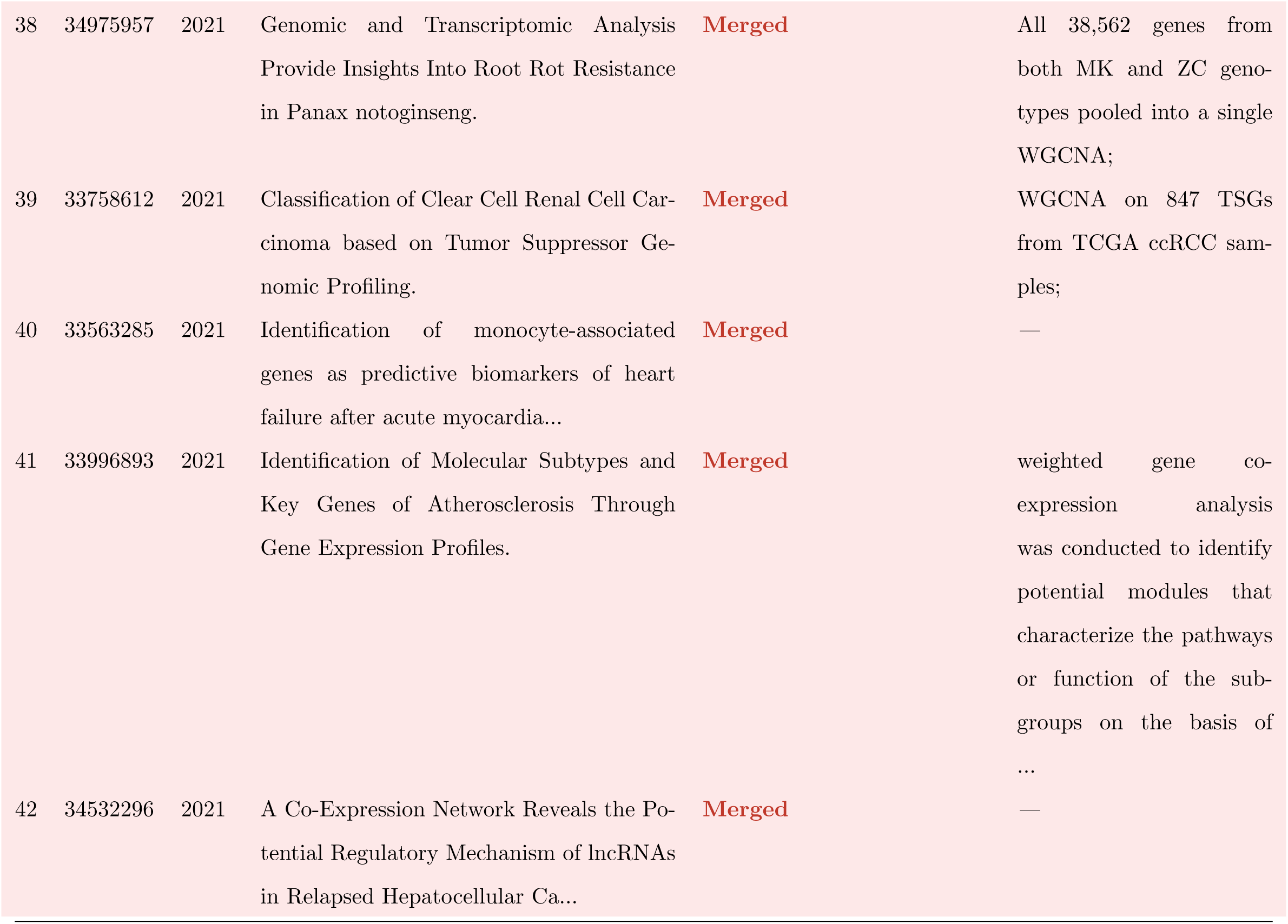
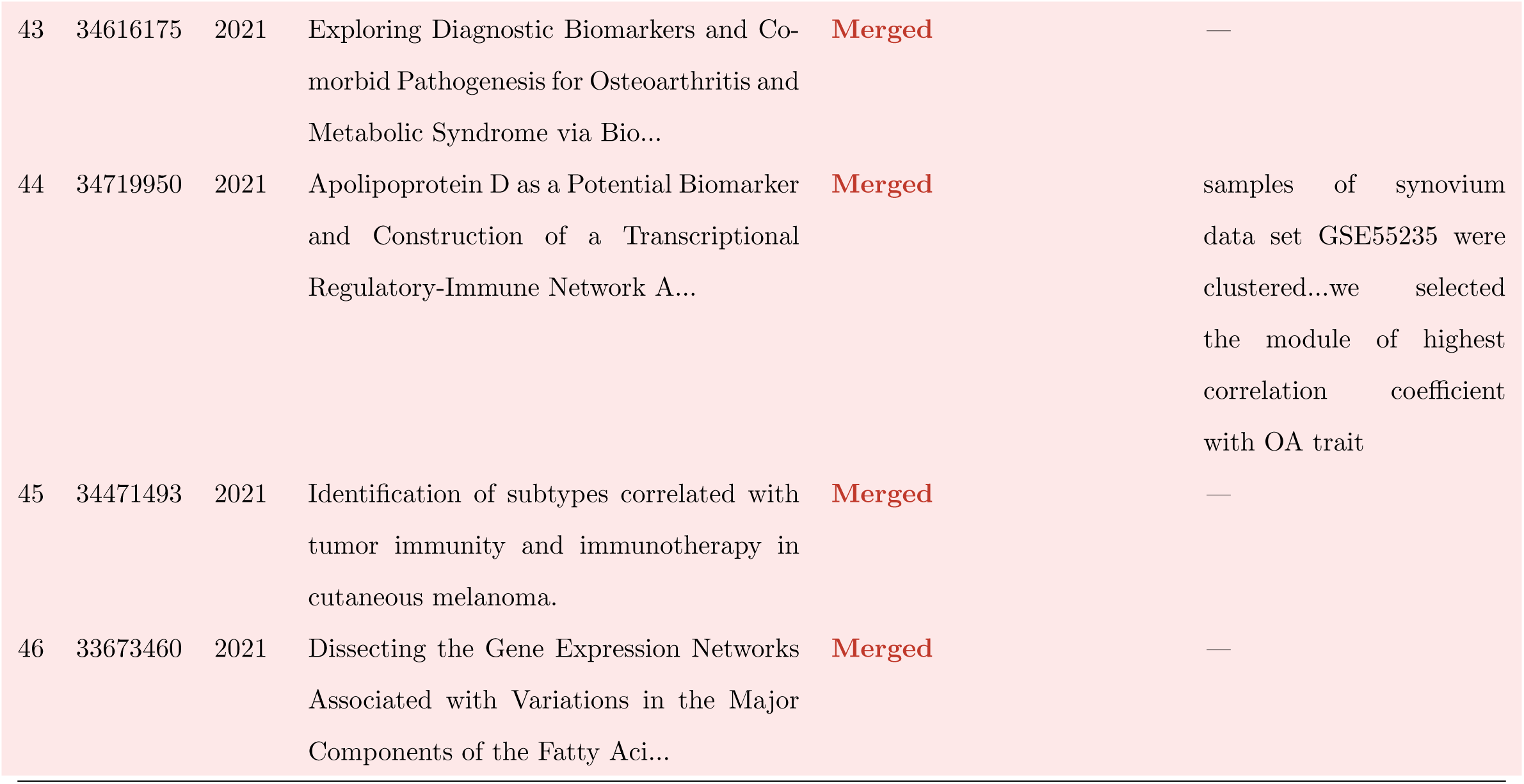
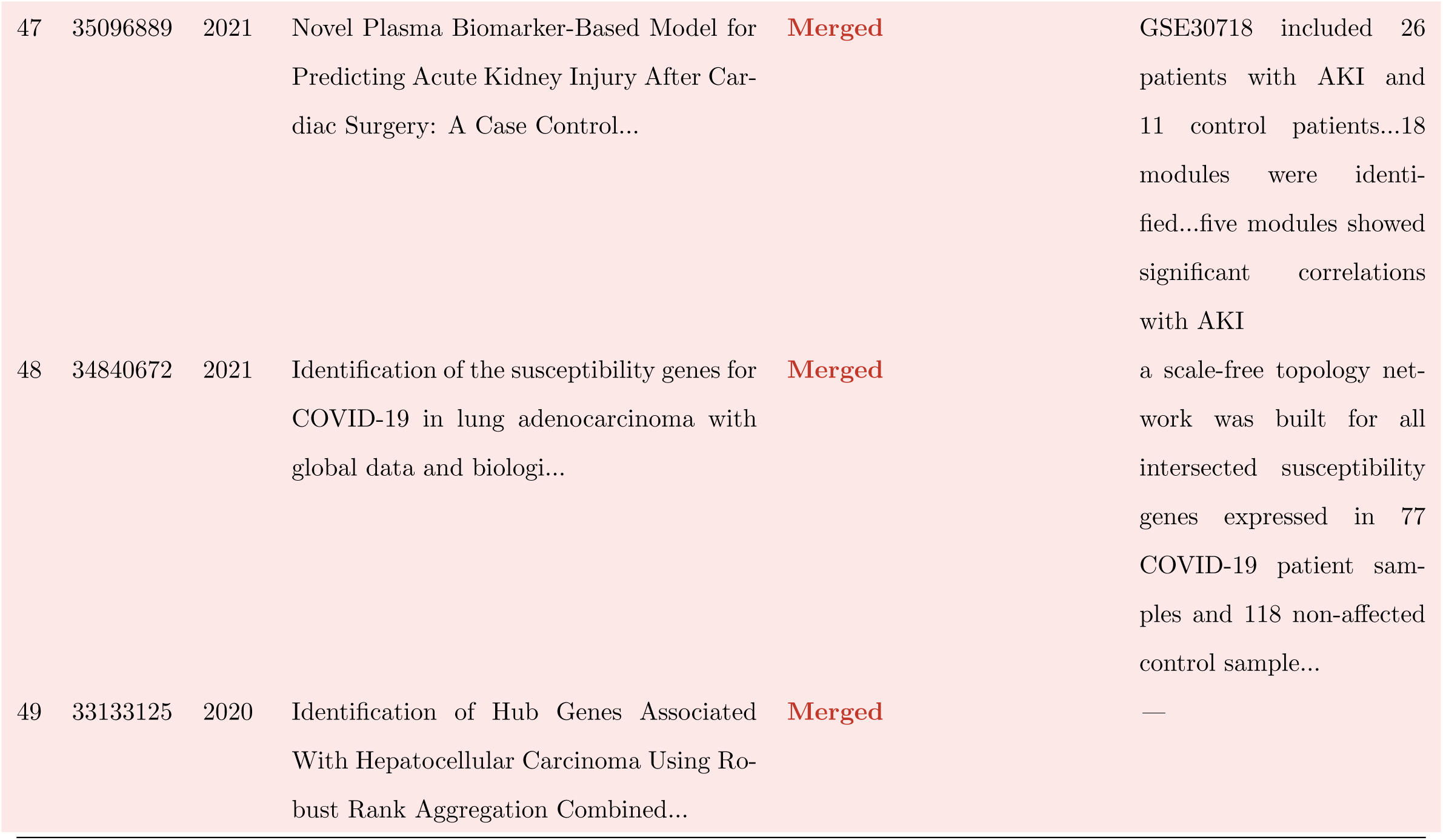
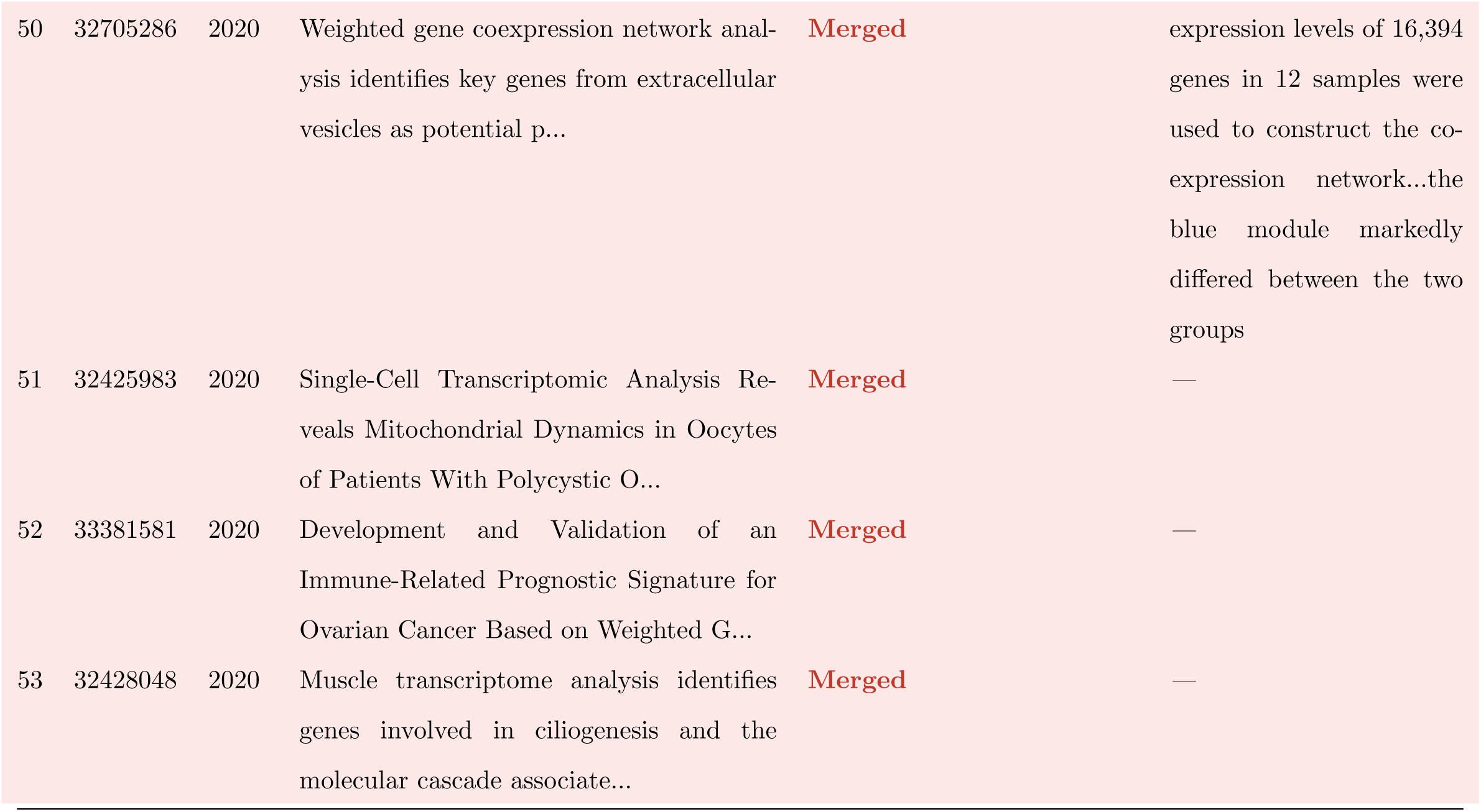
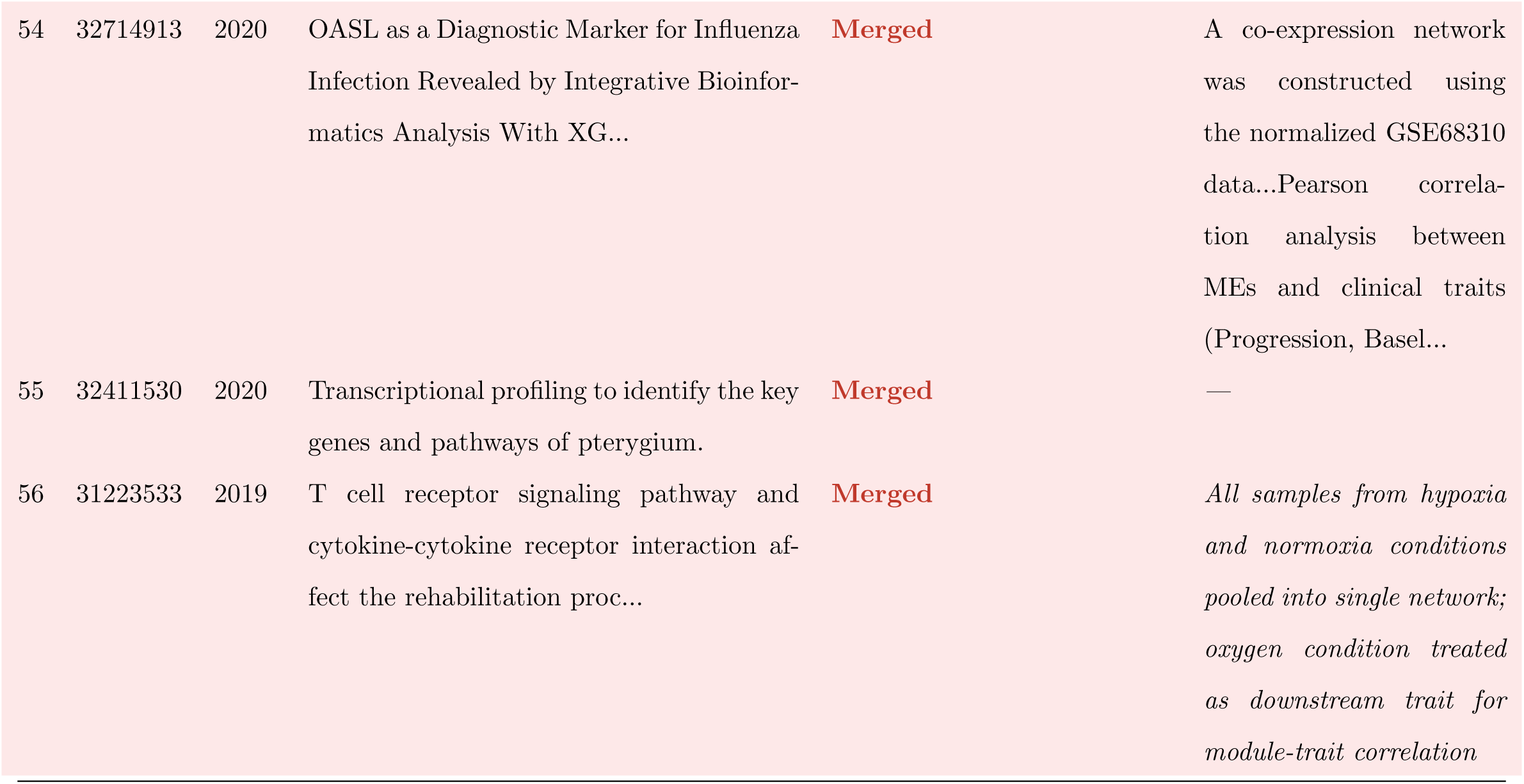
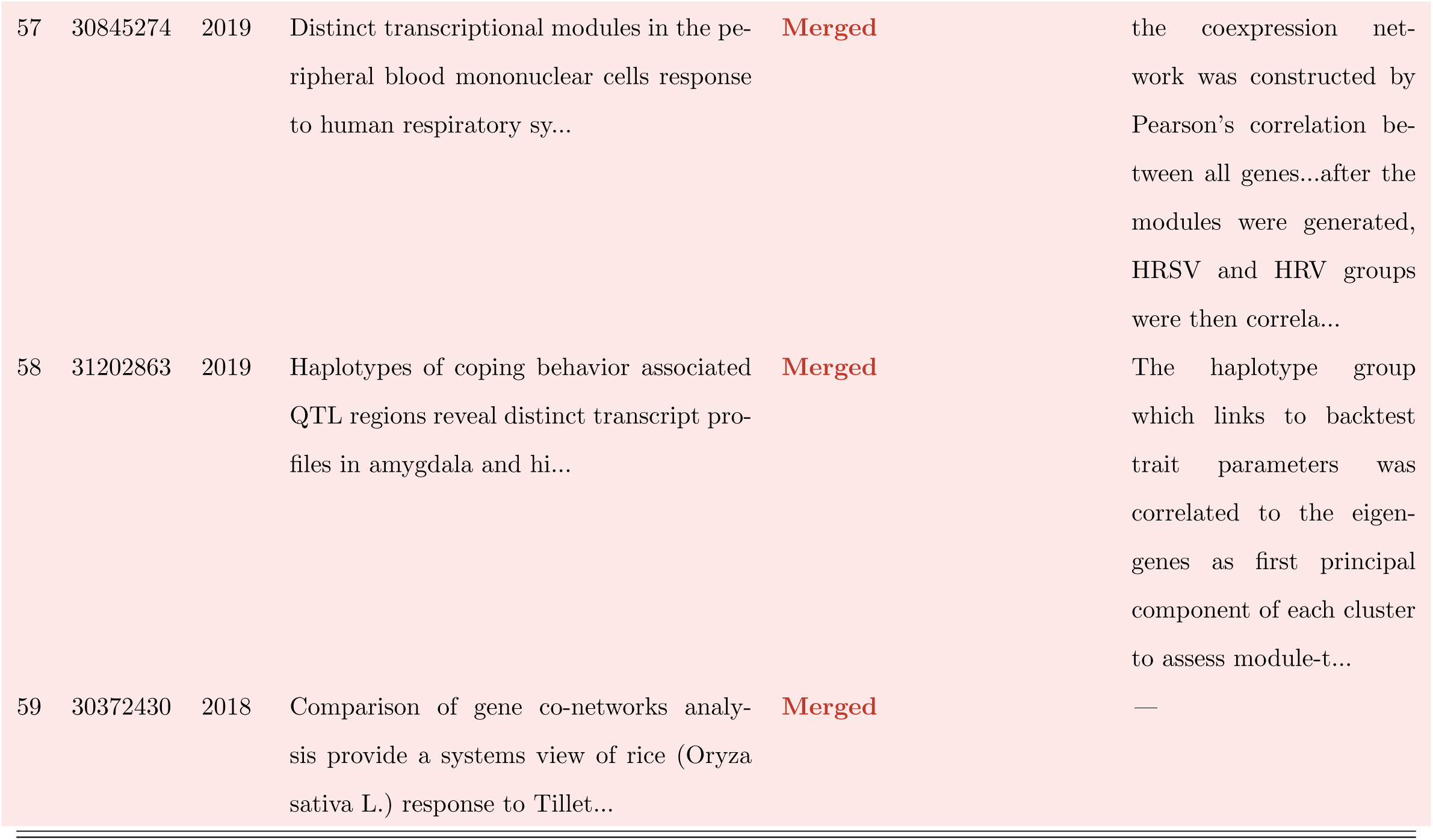
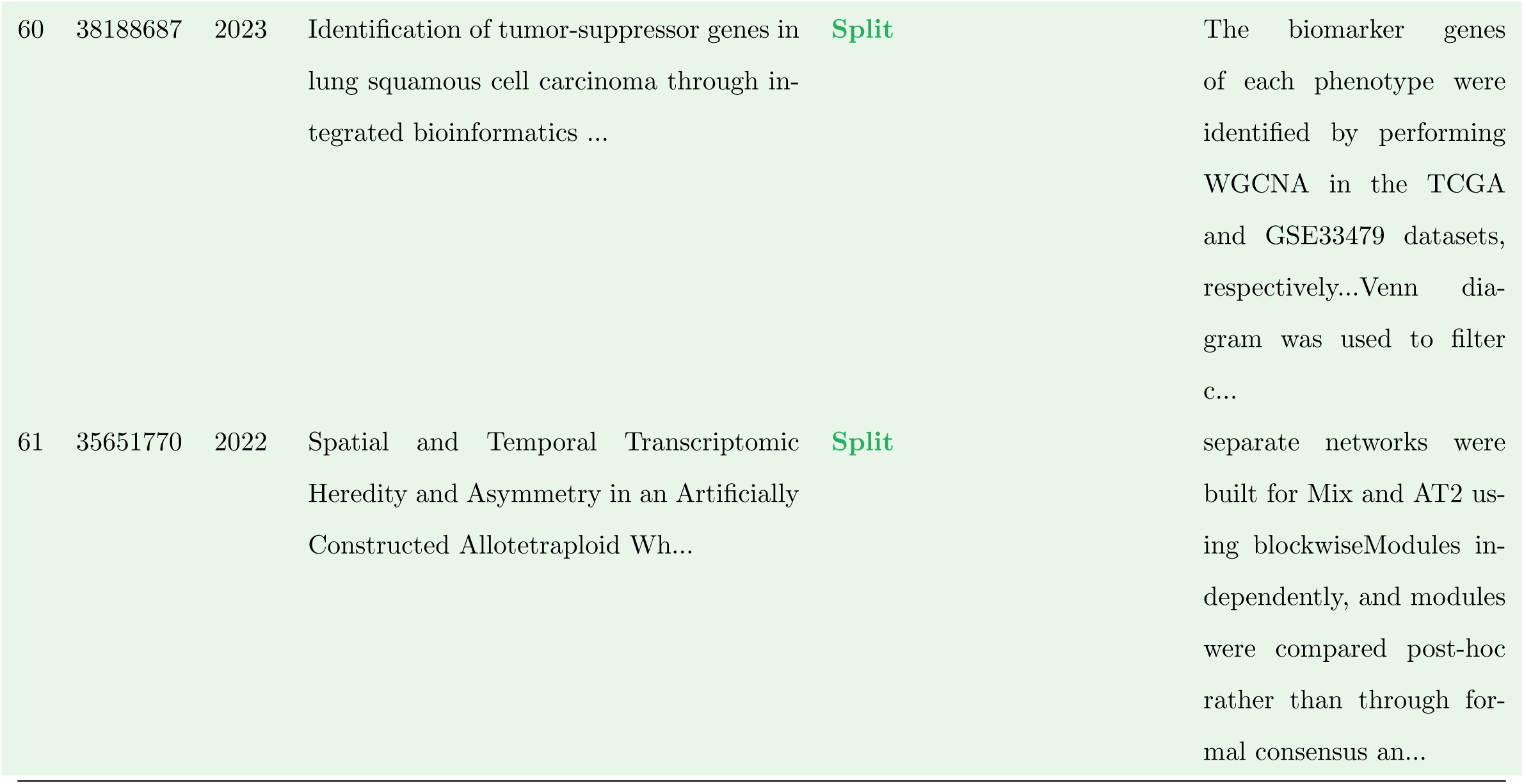
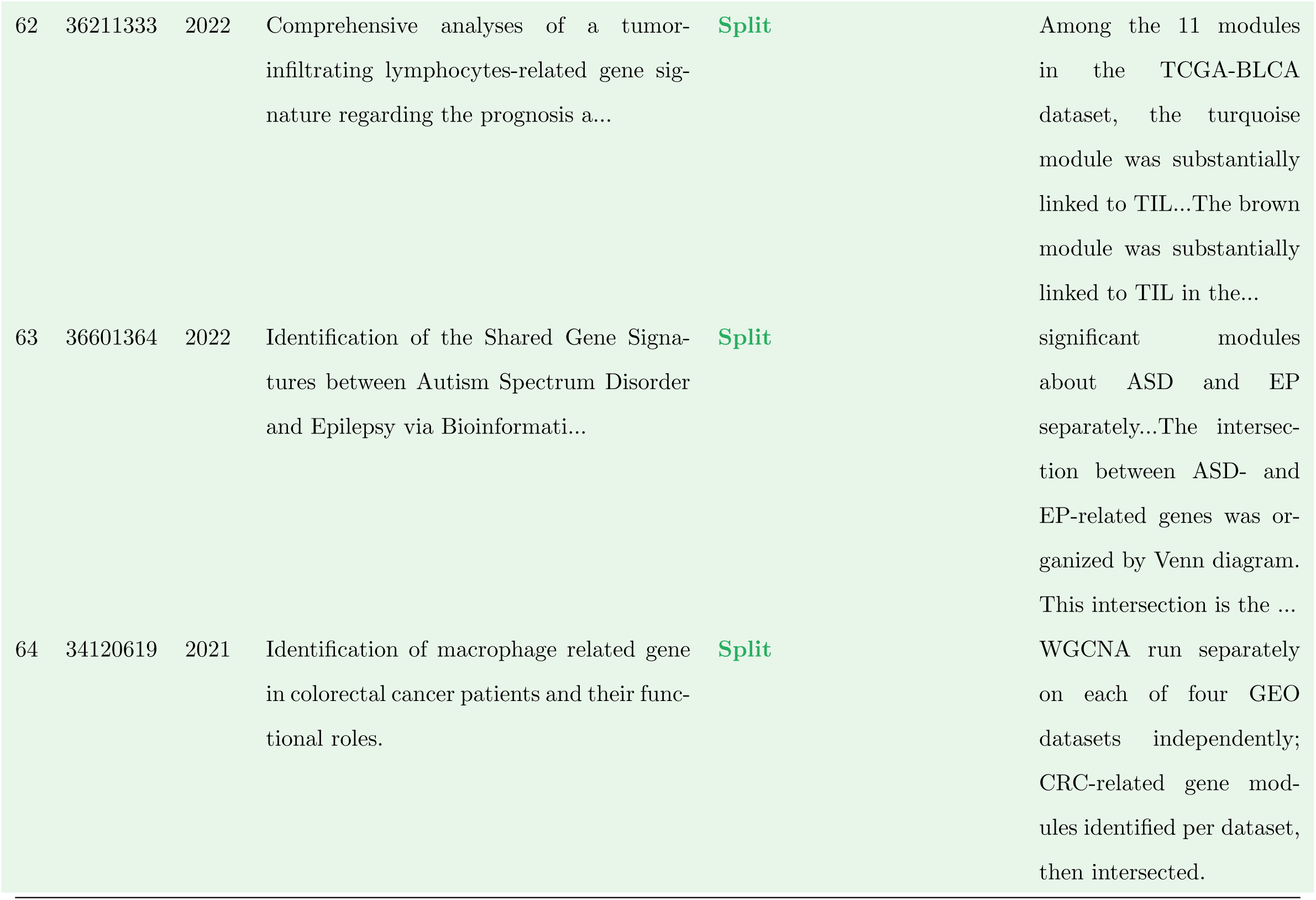
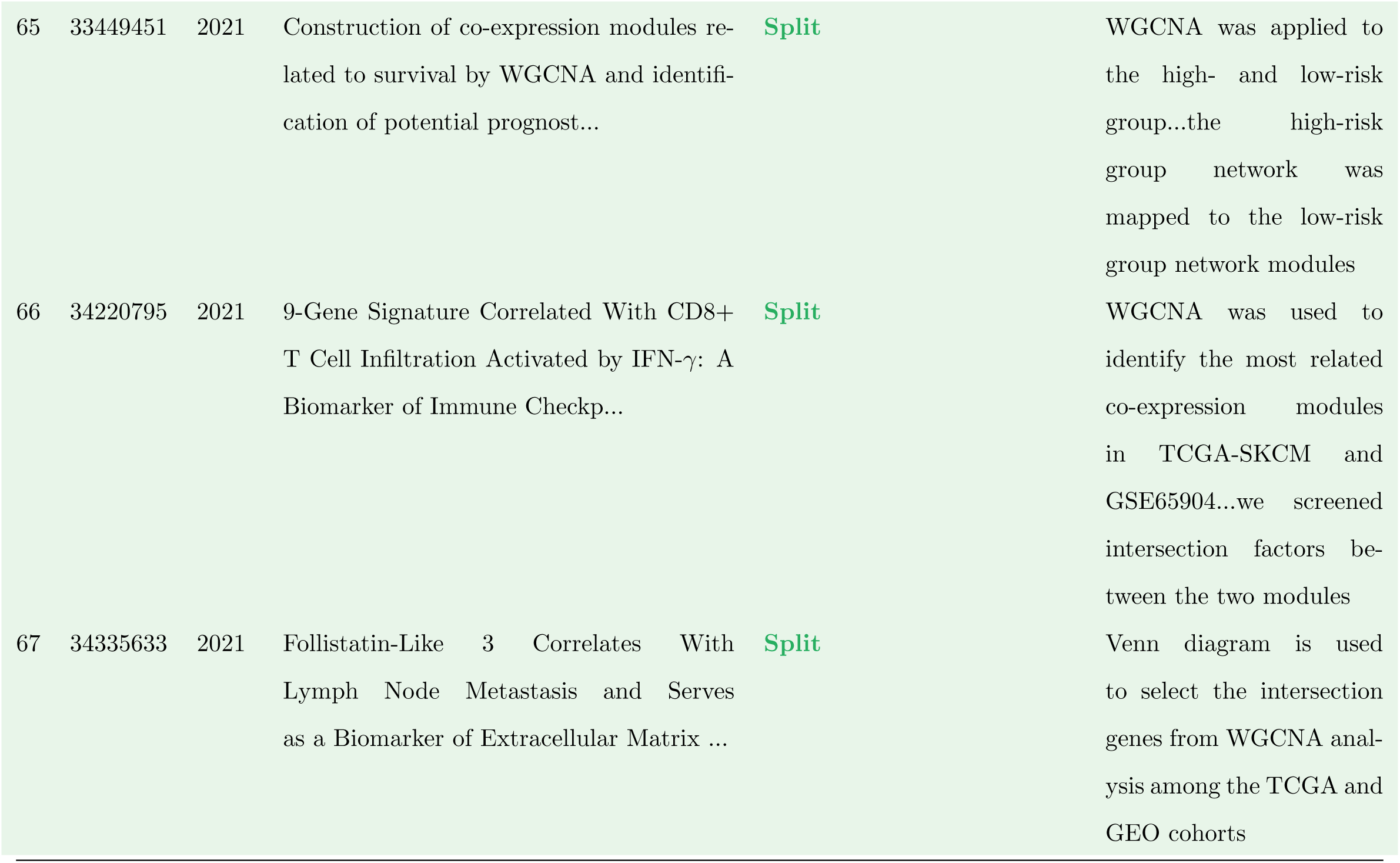
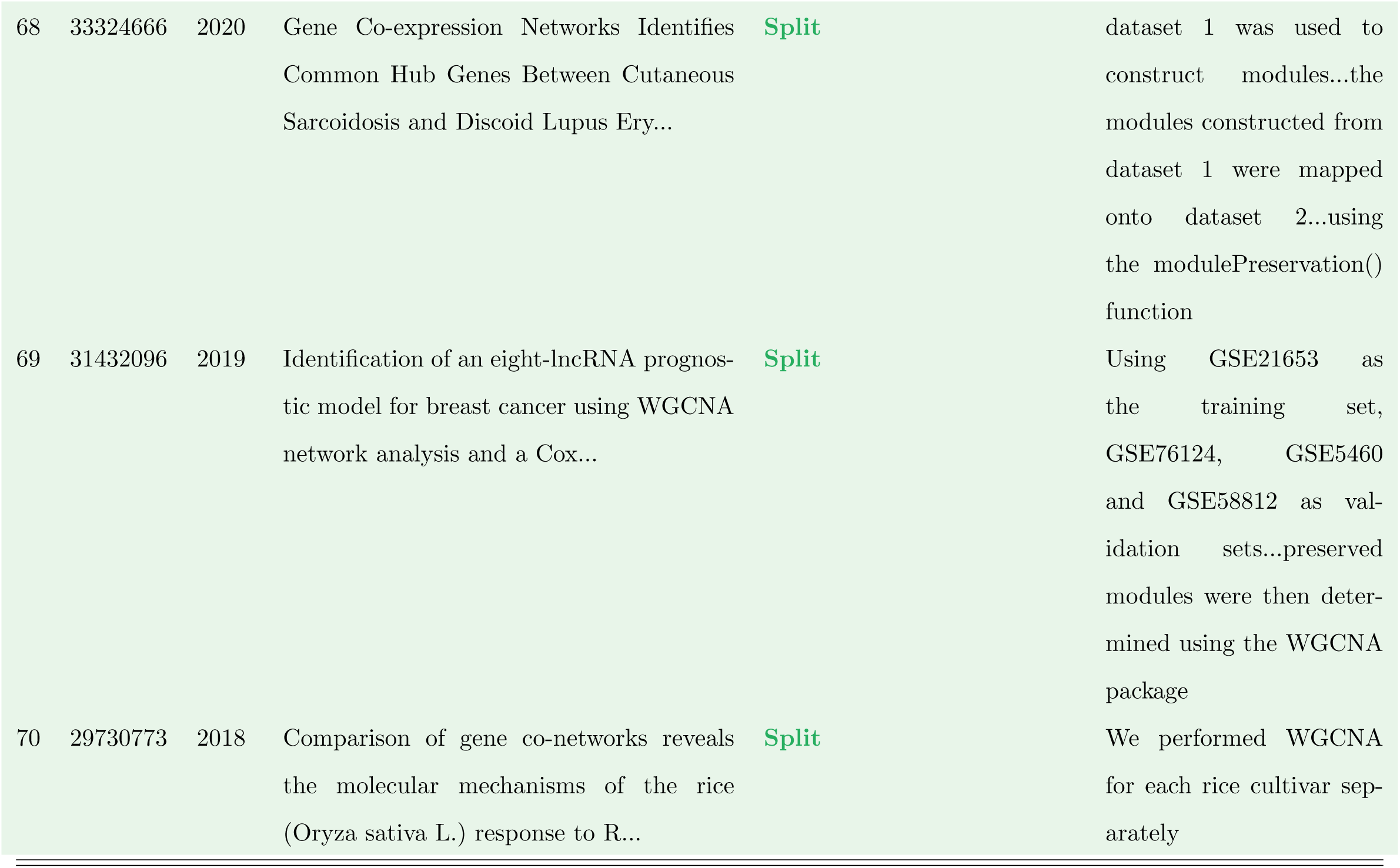
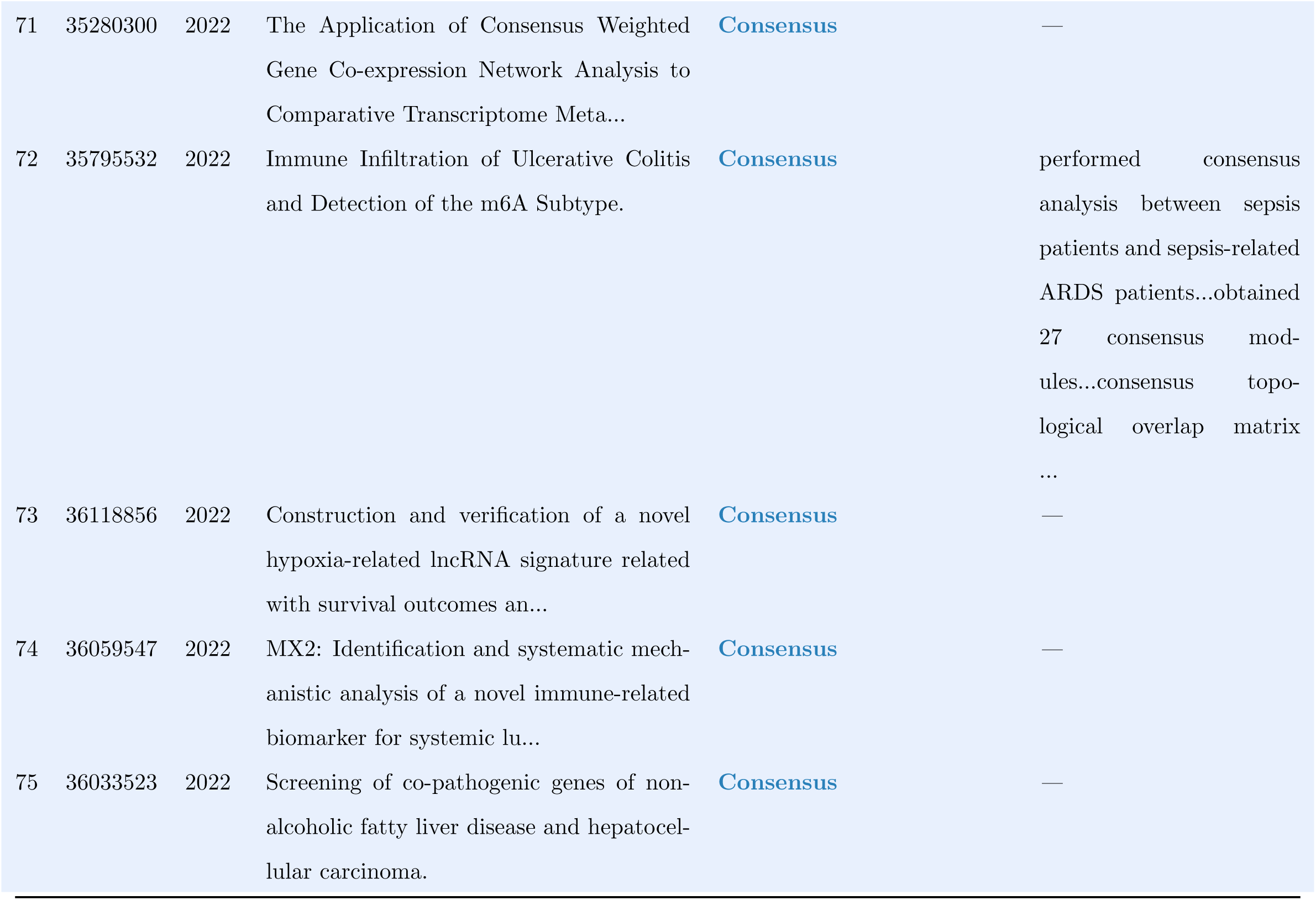
Complete coding sheet for included papers (. *N* = 75). Colour coding: Merged, Split, Consensus.

| # | PMID | Year | Title | Classification | Evidence / Notes |
| --- | --- | --- | --- | --- | --- |
| 1 | 38054380 | 2024 | Drug target genes and molecular mechanism investigation in isoflurane-induced anesthesia based on WGCNA and... | Merged | hub genes were investigated using weighted correlation network analysis (WGCNA)...183 DEGs integrated with genes revealed by WGCNA were identified as co-genes |
| 2 | 38187306 | 2024 | Construction of a diagnostic model for osteoarthritis based on transcriptomic immune-related genes. | Merged | expression levels of all genes in the training dataset...correlations with the clinical characteristics, encompassing diverse disease statuses and distinct i... |

Table S2 – *continued*
| # | PMID | Year | Title | Classification | Evidence / Notes |
| --- | --- | --- | --- | --- | --- |
| 3 | 38173477 | 2024 | Identification of the role of pyroptosis-related genes in chronic rhinosinusitis based on WGCNA. | Merged | we imported DEGs and sample labels into the WGCNA package for analysis...the correlation and significance between each module and CRS were calculated |
| 4 | 36866237 | 2023 | Novel Gene Signatures Promote Epithelial-Mesenchymal Transition (EMT) in Glucose Deprivation-Based Microenv... | Merged | WGCNA run on a single pooled expression matrix (TCGA HCC samples) to identify GD-status-associated modules |
| 5 | 36966811 | 2023 | The temporal and spatial signature of microglial transcriptome in neuropathic pain. | Merged | Single WGCNA on DEGs from the GSE60670 dataset with pain threshold scores as the clinical trait. |
| 6 | 37507701 | 2023 | GEMIN4, a potential therapeutic targets for patients with basal-like subtype breast cancer. | Merged | WGCNA applied to all BC subtypes together in a single expression matrix; |

Table S2 – continued
| # | PMID | Year | Title | Classification | Evidence / Notes |
| --- | --- | --- | --- | --- | --- |
| 7 | 37553678 | 2023 | LC/MS-based untargeted lipidomics reveals lipid signatures of nonpuerperal mastitis. | Merged | WGCNA on lipidomics data from NPM patients and controls pooled; |
| 8 | 37222809 | 2023 | Analysis of prognostic biomarker models and immune microenvironment in acute myeloid leukemia by integrativ... | Merged | WGCNA on TCGA-LAML tumor samples to find tumor-related genes; DEGs and WGCNA results intersected. Single pooled analysis. |
| 9 | 37457713 | 2023 | Mitochondrial DNA methylation is a predictor of immunotherapy response and prognosis in breast cancer: scRN... | Merged | co-expression network of all genes in MTDM and BC samples was constructed using the TCGA breast cancer patients' data cohort |

Table S2 – continued
| # | PMID | Year | Title | Classification | Evidence / Notes |
| --- | --- | --- | --- | --- | --- |
| 10 | 37204640 | 2023 | Associations between PBMC whole genome transcriptome, muscle strength, muscle mass, and physical performanc... | Merged | we extracted the residuals and subsequently used them as input for the WGCNA algorithm...WGCNA::blockwiseModules produced gene expression clusters |
| 11 | 37152997 | 2023 | Transcriptome analysis of two pepper genotypes infected with pepper mild motile virus. | Merged | WGCNA of all samples...a co-expression network was built based on pairwise correlations of gene expression in all samples |
| 12 | 38050560 | 2023 | A Macrophage-Related Gene Signature for Identifying COPD Based on Bioinformatics and ex vivo Experiments. | Merged | The fractions of vital immune infiltrating cells in COPD patients were used as sample traits. We identified the key modules according to a high correlation b... |

Table S2 – *continued*
| # | PMID | Year | Title | Classification | Evidence / Notes |
| --- | --- | --- | --- | --- | --- |
| 13 | 37312900 | 2023 | Integrated Analysis of Single-Cell RNA-Seq and Bulk RNA-Seq Combined with Multiple Machine Learning Identif... | **Merged** | — |
| 14 | 37876534 | 2023 | Constructing a prognostic model for head and neck squamous cell carcinoma based on glucose metabolism relat... | **Merged** | 500 HNSC and 44 Normal samples were employed in build a co-expression network by WGCNA...key module was acquired by correlation analysis between module and HNSC |
| 15 | 37366074 | 2023 | Integrating sperm cell transcriptome and seminal plasma metabolome to analyze the molecular regulatory mech... | **Merged** | — |
| 16 | 37592228 | 2023 | Molecular regulatory mechanism of key LncRNAs in subclinical mastitic cows with folic acid supplementation. | **Merged** | WGCNA of all the protein-coding genes detected in this study was performed...module characteristic gene associated with the phenotype matrix to obtain correl... |

Table S2 – continued
| # | PMID | Year | Title | Classification | Evidence / Notes |
| --- | --- | --- | --- | --- | --- |
| 17 | 37588985 | 2023 | Identification of fibroblast-related genes based on single-cell and machine learning to predict the prognos... | Merged | Cluster analysis was first performed on 177 tumor samples and 167 normal samples to further calculate the Pearson's correlation between each gene, followed b... |
| 18 | 36610580 | 2023 | Transcriptome analysis reveals new insights in the starch biosynthesis of non-waxy and waxy broomcorn mille... | Merged | 50,725 genes were divided into 23 modules...associated each co-expression module with starch content and key starch synthesis enzyme activities via Pearson's... |
| 19 | 37456238 | 2023 | A tumor mutational burden-derived immune computational framework selects sensitive immunotherapy/chemothera... | Merged | — |

Table S2 – continued
| # | PMID | Year | Title | Classification | Evidence / Notes |
| --- | --- | --- | --- | --- | --- |
| 20 | 36335813 | 2022 | An immune-related gene signature for the prognosis of human bladder cancer based on WGCNA. | Merged | a co-expression network for all genes in BLCA samples was constructed by the R-package WGCNA |
| 21 | 36091017 | 2022 | Plasma exosomal IRAK1 can be a potential biomarker for predicting the treatment response to renin-angiotens... | Merged | — |
| 22 | 34775485 | 2022 | Epigenome-wide association study of alcohol use disorder in five brain regions. | Merged | WGCNA run per brain region on all samples (AUD cases + controls pooled); |
| 23 | 36138874 | 2022 | A Metabolic Plasticity-Based Signature for Molecular Classification and Prognosis of Lower-Grade Glioma. | Merged | WGCNA built on DEGs from the CGGA array (301) cohort with tumor grade as the module-trait variable. |

Table S2 – *continued*
| # | PMID | Year | Title | Classification | Evidence / Notes |
| --- | --- | --- | --- | --- | --- |
| 24 | 35853879 | 2022 | Gene co-expression architecture in peripheral blood in a cohort of remitted first-episode schizophrenia pat... | Merged | single WGCNA on all 91 psychosis patients; clinical scales (PANSS, GAF, etc.) used as module-trait correlations. No group splitting. |
| 25 | 35332109 | 2022 | Tumor-associated macrophages related signature in glioma. | Merged | WGCNA run on DEGs from the TCGA GBMLGG cohort; macrophage infiltration level used as the module-trait variable. |
| 26 | 35227285 | 2022 | Construction and validation of a transcription factors-based prognostic signature for ovarian cancer. | Merged | — |
| 27 | 34695258 | 2022 | Comparative transcriptome analysis of winter yaks in plateau and plain. | Merged | correlations between average expression levels in modules and real phenotype values were used to detect candidate genes with such traits |

Table S2 – *continued*
| # | PMID | Year | Title | Classification | Evidence / Notes |
| --- | --- | --- | --- | --- | --- |
| 28 | 35398514 | 2022 | Structural brain characteristics and gene co-expression analysis: A study with outcome label from normal co... | Merged | DEGs were clustered into different modules. Then, the relationship of modules and cortical areas were established |
| 29 | 35532246 | 2022 | Identification of a novel lncRNA-miRNA-mRNA competing endogenous RNA network associated with prognosis of b... | Merged | the similarity between two groups of genes was analyzed by WGCNA...WGCNA predicted that 823 lncRNAs and 1813 mRNAs were closely related to breast cancer |
| 30 | 36543847 | 2022 | Analysis of prognostic model based on immunotherapy related genes in lung adenocarcinoma. | Merged | the gene expression matrix of 5000 pretreatment genes was analyzed by WGCNA...correlation coefficient between each module and the samples related to the char... |

Table S2 – *continued*
| # | PMID | Year | Title | Classification | Evidence / Notes |
| --- | --- | --- | --- | --- | --- |
| 31 | 35223836 | 2022 | Clinical Significance and Immune Landscape of a Pyroptosis-Derived LncRNA Signature for Glioblastoma. | **Merged** | — |
| 32 | 35571180 | 2022 | Exploring Key Genes to Construct a Diagnosis Model of Dilated Cardiomyopathy. | **Merged** | a gene expression matrix of overlapped genes was obtained after taking the interaction of five datasets, which was the input file of WGCNA...the modules that... |
| 33 | 36203606 | 2022 | Identification of immune-related endoplasmic reticulum stress genes in sepsis using bioinformatics and mach... | **Merged** | — |
| 34 | 35965814 | 2022 | Identification of key genes related to immune cells in patients with gram-negative sepsis based on weighted... | **Merged** | — |

Table S2 – *continued*
| # | PMID | Year | Title | Classification | Evidence / Notes |
| --- | --- | --- | --- | --- | --- |
| 35 | 36553607 | 2022 | Transcriptional Specificity Analysis of Testis and Epididymis Tissues in Donkey. | Merged | the data of three tissues, including donkey testis, donkey epididymis and horse testis, were employed with a total of 16 samples...correlation between tissue... |
| 36 | 35812950 | 2022 | Gene Co-expression Network Analysis of the Comparative Transcriptome Identifies Hub Genes Associated With R... | Merged | WGCNA of 28,579 DEGs was carried out...all genes were assigned into 16 distinct modules...modules closely associated with resistance to A. flavus were identi... |
| 37 | 33781203 | 2021 | The mechanism of sesame resistance against Macrophomina phaseolina was revealed via a comparison of transcr... | Merged | “the scale-free co-expression network was constructed by using the FPKM matrix transformed by log2” |

Table S2 – continued
| # | PMID | Year | Title | Classification | Evidence / Notes |
| --- | --- | --- | --- | --- | --- |
| 38 | 34975957 | 2021 | Genomic and Transcriptomic Analysis Provide Insights Into Root Rot Resistance in Panax notoginseng. | Merged | All 38,562 genes from both MK and ZC genotypes pooled into a single WGCNA; |
| 39 | 33758612 | 2021 | Classification of Clear Cell Renal Cell Carcinoma based on Tumor Suppressor Genomic Profiling. | Merged | WGCNA on 847 TSGs from TCGA ccRCC samples; |
| 40 | 33563285 | 2021 | Identification of monocyte-associated genes as predictive biomarkers of heart failure after acute myocardia... | Merged | — |
| 41 | 33996893 | 2021 | Identification of Molecular Subtypes and Key Genes of Atherosclerosis Through Gene Expression Profiles. | Merged | weighted gene co-expression analysis was conducted to identify potential modules that characterize the pathways or function of the subgroups on the basis of ... |
| 42 | 34532296 | 2021 | A Co-Expression Network Reveals the Potential Regulatory Mechanism of lncRNAs in Relapsed Hepatocellular Ca... | Merged | — |

Table S2 – *continued*
| # | PMID | Year | Title | Classification | Evidence / Notes |
| --- | --- | --- | --- | --- | --- |
| 43 | 34616175 | 2021 | Exploring Diagnostic Biomarkers and Co-morbid Pathogenesis for Osteoarthritis and Metabolic Syndrome via Bio... | **Merged** | — |
| 44 | 34719950 | 2021 | Apolipoprotein D as a Potential Biomarker and Construction of a Transcriptional Regulatory-Immune Network A... | **Merged** | samples of synovium data set GSE55235 were clustered...we selected the module of highest correlation coefficient with OA trait |
| 45 | 34471493 | 2021 | Identification of subtypes correlated with tumor immunity and immunotherapy in cutaneous melanoma. | **Merged** | — |
| 46 | 33673460 | 2021 | Dissecting the Gene Expression Networks Associated with Variations in the Major Components of the Fatty Aci... | **Merged** | — |

Table S2 – continued
| # | PMID | Year | Title | Classification | Evidence / Notes |
| --- | --- | --- | --- | --- | --- |
| 47 | 35096889 | 2021 | Novel Plasma Biomarker-Based Model for Predicting Acute Kidney Injury After Cardiac Surgery: A Case Control... | Merged | GSE30718 included 26 patients with AKI and 11 control patients...18 modules were identified...five modules showed significant correlations with AKI |
| 48 | 34840672 | 2021 | Identification of the susceptibility genes for COVID-19 in lung adenocarcinoma with global data and biologi... | Merged | a scale-free topology network was built for all intersected susceptibility genes expressed in 77 COVID-19 patient samples and 118 non-affected control sample... |
| 49 | 33133125 | 2020 | Identification of Hub Genes Associated With Hepatocellular Carcinoma Using Robust Rank Aggregation Combined... | Merged | — |

Table S2 – continued
| # | PMID | Year | Title | Classification | Evidence / Notes |
| --- | --- | --- | --- | --- | --- |
| 50 | 32705286 | 2020 | Weighted gene coexpression network analysis identifies key genes from extracellular vesicles as potential p... | Merged | expression levels of 16,394 genes in 12 samples were used to construct the co-expression network...the blue module markedly differed between the two groups |
| 51 | 32425983 | 2020 | Single-Cell Transcriptomic Analysis Reveals Mitochondrial Dynamics in Oocytes of Patients With Polycystic O... | Merged | — |
| 52 | 33381581 | 2020 | Development and Validation of an Immune-Related Prognostic Signature for Ovarian Cancer Based on Weighted G... | Merged | — |
| 53 | 32428048 | 2020 | Muscle transcriptome analysis identifies genes involved in ciliogenesis and the molecular cascade associate... | Merged | — |

Table S2 – continued
| # | PMID | Year | Title | Classification | Evidence / Notes |
| --- | --- | --- | --- | --- | --- |
| 54 | 32714913 | 2020 | OASL as a Diagnostic Marker for Influenza Infection Revealed by Integrative Bioinformatics Analysis With XG... | Merged | A co-expression network was constructed using the normalized GSE68310 data...Pearson correlation analysis between MEs and clinical traits (Progression, Basel... |
| 55 | 32411530 | 2020 | Transcriptional profiling to identify the key genes and pathways of pterygium. | Merged | — |
| 56 | 31223533 | 2019 | T cell receptor signaling pathway and cytokine-cytokine receptor interaction affect the rehabilitation proc... | Merged | All samples from hypoxia and normoxia conditions pooled into single network; oxygen condition treated as downstream trait for module-trait correlation |

Table S2 – continued
| # | PMID | Year | Title | Classification | Evidence / Notes |
| --- | --- | --- | --- | --- | --- |
| 57 | 30845274 | 2019 | Distinct transcriptional modules in the peripheral blood mononuclear cells response to human respiratory sy... | Merged | the coexpression network was constructed by Pearson's correlation between all genes...after the modules were generated, HRSV and HRV groups were then correla... |
| 58 | 31202863 | 2019 | Haplotypes of coping behavior associated QTL regions reveal distinct transcript profiles in amygdala and hi... | Merged | The haplotype group which links to backtest trait parameters was correlated to the eigen-genes as first principal component of each cluster to assess module-t... |
| 59 | 30372430 | 2018 | Comparison of gene co-networks analysis provide a systems view of rice (Oryza sativa L.) response to Tillet... | Merged | — |

Table S2 – continued
| # | PMID | Year | Title | Classification | Evidence / Notes |
| --- | --- | --- | --- | --- | --- |
| 60 | 38188687 | 2023 | Identification of tumor-suppressor genes in lung squamous cell carcinoma through integrated bioinformatics ... | Split | The biomarker genes of each phenotype were identified by performing WGCNA in the TCGA and GSE33479 datasets, respectively...Venn diagram was used to filter c... |
| 61 | 35651770 | 2022 | Spatial and Temporal Transcriptomic Heredity and Asymmetry in an Artificially Constructed Allotetraploid Wh... | Split | separate networks were built for Mix and AT2 using blockwiseModules independently, and modules were compared post-hoc rather than through formal consensus an... |

Table S2 – *continued*
| # | PMID | Year | Title | Classification | Evidence / Notes |
| --- | --- | --- | --- | --- | --- |
| 62 | 36211333 | 2022 | Comprehensive analyses of a tumor-infiltrating lymphocytes-related gene signature regarding the prognosis a... | Split | Among the 11 modules in the TCGA-BLCA dataset, the turquoise module was substantially linked to TIL...The brown module was substantially linked to TIL in the... |
| 63 | 36601364 | 2022 | Identification of the Shared Gene Signatures between Autism Spectrum Disorder and Epilepsy via Bioinformati... | Split | significant modules about ASD and EP separately...The intersection between ASD- and EP-related genes was organized by Venn diagram. This intersection is the ... |
| 64 | 34120619 | 2021 | Identification of macrophage related gene in colorectal cancer patients and their functional roles. | Split | WGCNA run separately on each of four GEO datasets independently; CRC-related gene modules identified per dataset, then intersected. |

Table S2 – *continued*
| # | PMID | Year | Title | Classification | Evidence / Notes |
| --- | --- | --- | --- | --- | --- |
| 65 | 33449451 | 2021 | Construction of co-expression modules related to survival by WGCNA and identification of potential prognost... | Split | WGCNA was applied to the high- and low-risk group...the high-risk group network was mapped to the low-risk group network modules |
| 66 | 34220795 | 2021 | 9-Gene Signature Correlated With CD8+ T Cell Infiltration Activated by IFN-γ: A Biomarker of Immune Checkp... | Split | WGCNA was used to identify the most related co-expression modules in TCGA-SKCM and GSE65904...we screened intersection factors between the two modules |
| 67 | 34335633 | 2021 | Follistatin-Like 3 Correlates With Lymph Node Metastasis and Serves as a Biomarker of Extracellular Matrix ... | Split | Venn diagram is used to select the intersection genes from WGCNA analysis among the TCGA and GEO cohorts |

Table S2 – *continued*
| # | PMID | Year | Title | Classification | Evidence / Notes |
| --- | --- | --- | --- | --- | --- |
| 68 | 33324666 | 2020 | Gene Co-expression Networks Identifies Common Hub Genes Between Cutaneous Sarcoidosis and Discoid Lupus Ery... | Split | dataset 1 was used to construct modules...the modules constructed from dataset 1 were mapped onto dataset 2...using the modulePreservation() function |
| 69 | 31432096 | 2019 | Identification of an eight-lncRNA prognostic model for breast cancer using WGCNA network analysis and a Cox... | Split | Using GSE21653 as the training set, GSE76124, GSE5460 and GSE58812 as validation sets...preserved modules were then determined using the WGCNA package |
| 70 | 29730773 | 2018 | Comparison of gene co-networks reveals the molecular mechanisms of the rice (Oryza sativa L.) response to R... | Split | We performed WGCNA for each rice cultivar separately |

Table S2 – *continued*
| # | PMID | Year | Title | Classification | Evidence / Notes |
| --- | --- | --- | --- | --- | --- |
| 71 | 35280300 | 2022 | The Application of Consensus Weighted Gene Co-expression Network Analysis to Comparative Transcriptome Meta... | Consensus | — |
| 72 | 35795532 | 2022 | Immune Infiltration of Ulcerative Colitis and Detection of the m6A Subtype. | Consensus | performed consensus analysis between sepsis patients and sepsis-related ARDS patients...obtained 27 consensus modules...consensus topological overlap matrix ... |
| 73 | 36118856 | 2022 | Construction and verification of a novel hypoxia-related lncRNA signature related with survival outcomes an... | Consensus | — |
| 74 | 36059547 | 2022 | MX2: Identification and systematic mechanistic analysis of a novel immune-related biomarker for systemic lu... | Consensus | — |
| 75 | 36033523 | 2022 | Screening of co-pathogenic genes of non-alcoholic fatty liver disease and hepatocellular carcinoma. | Consensus | — |

**Table S3:**
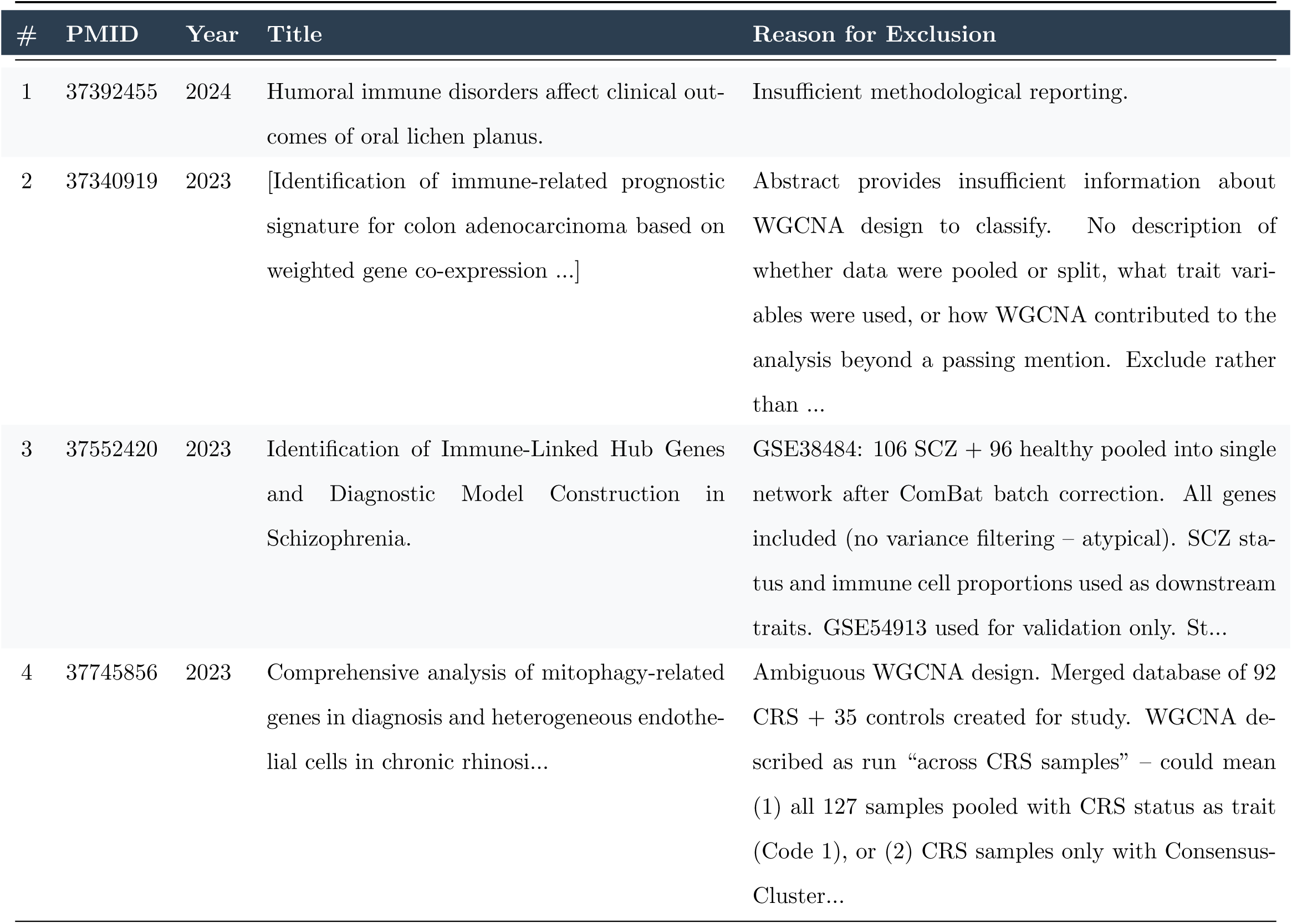
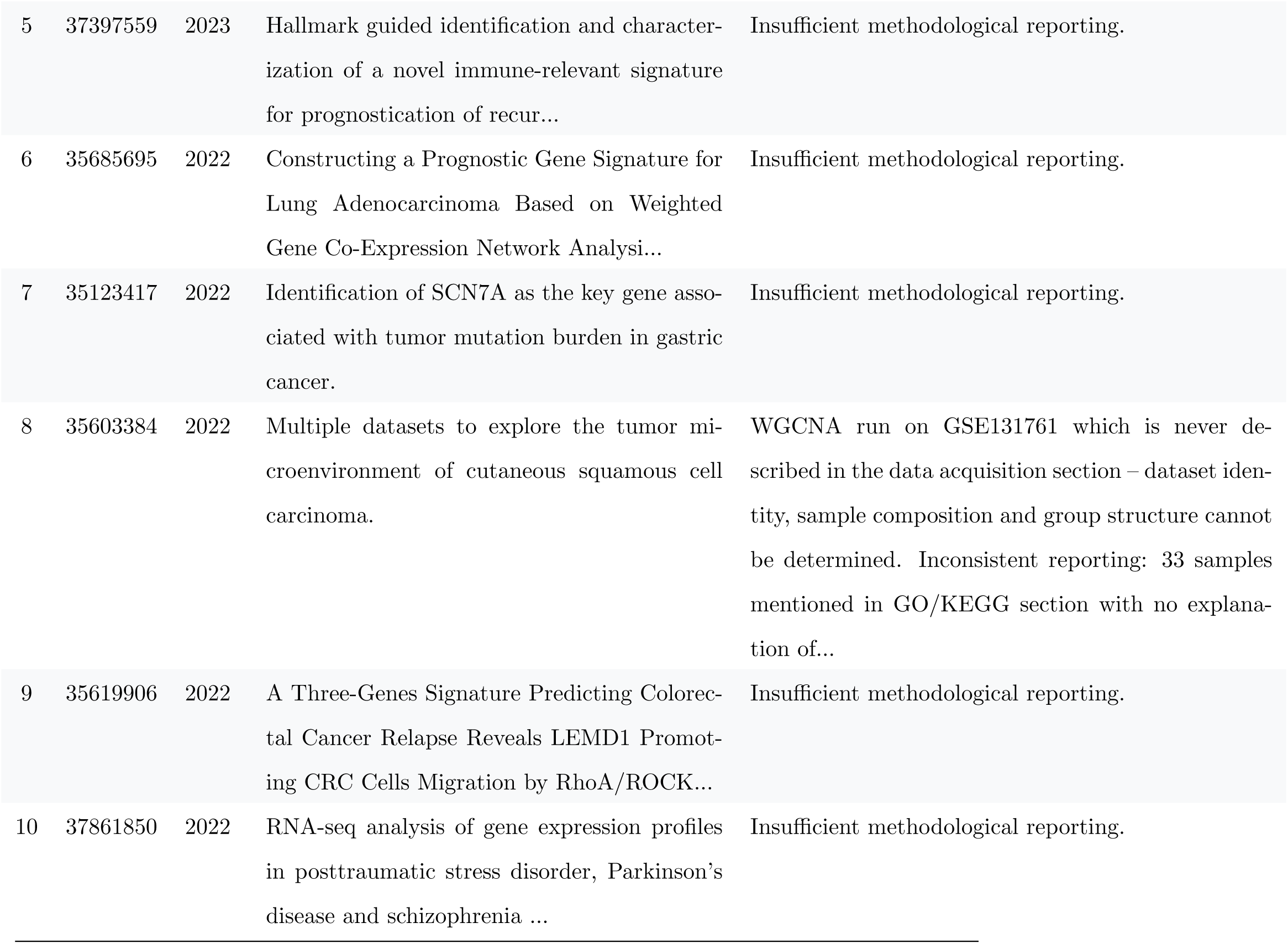
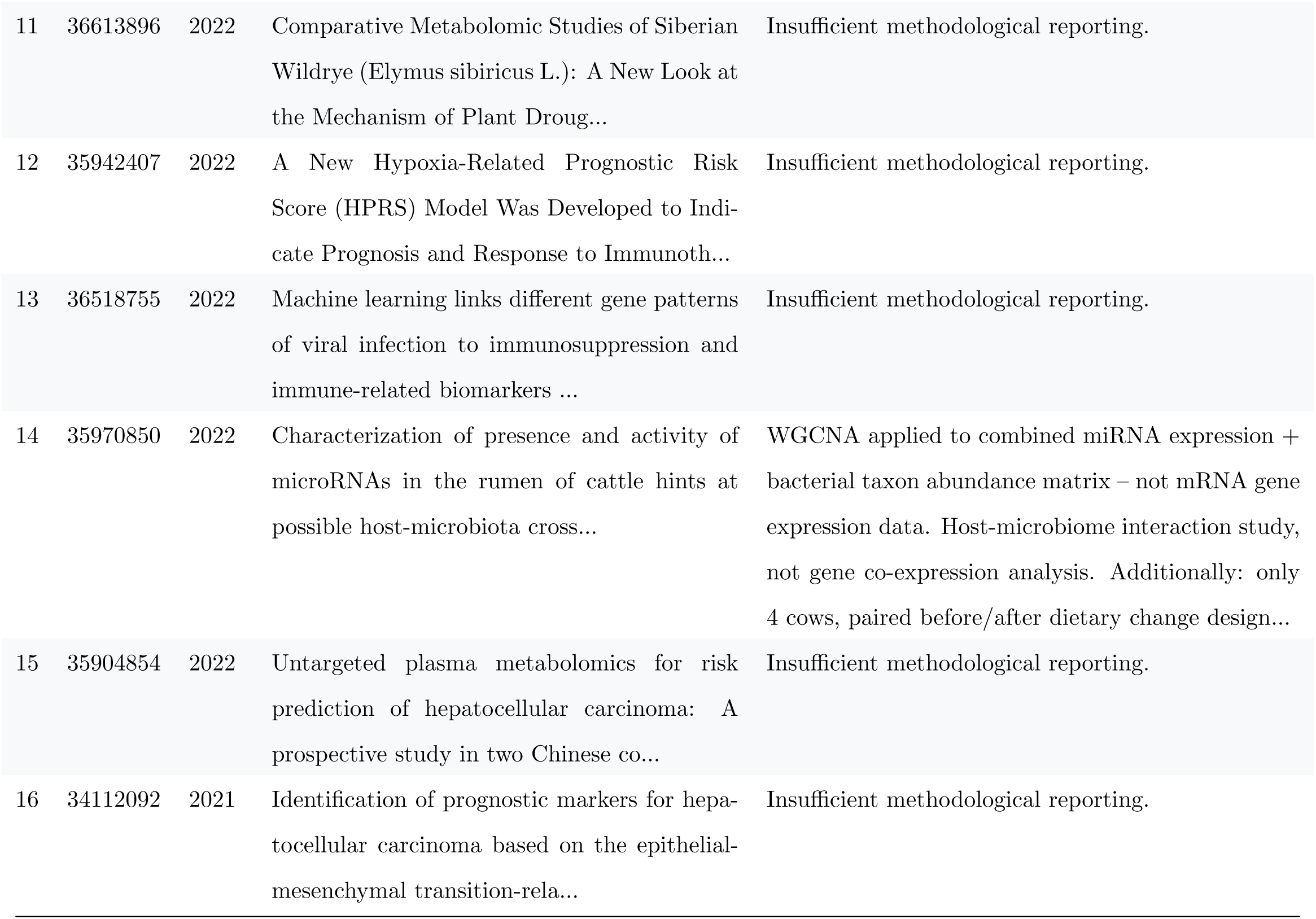
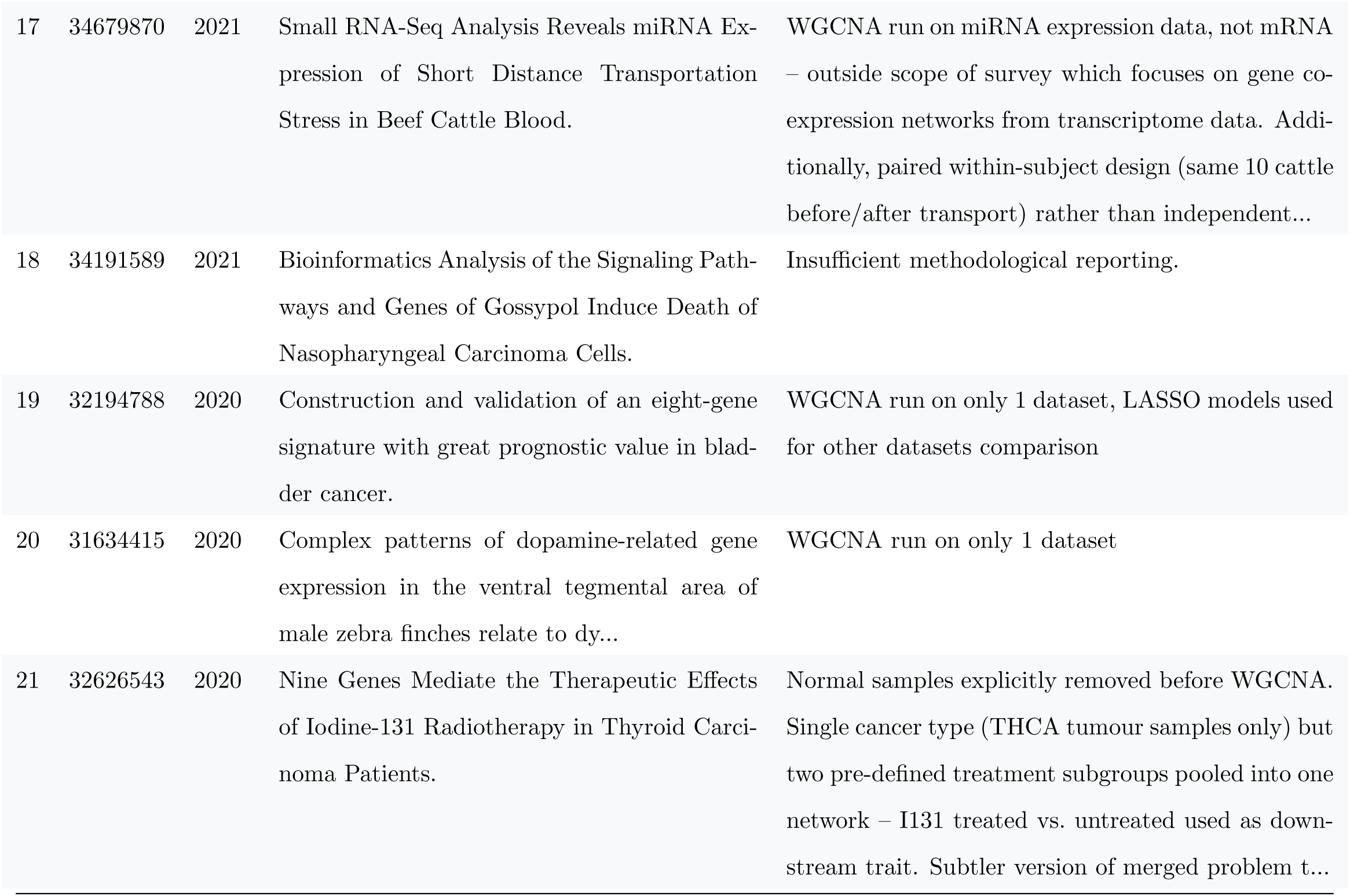
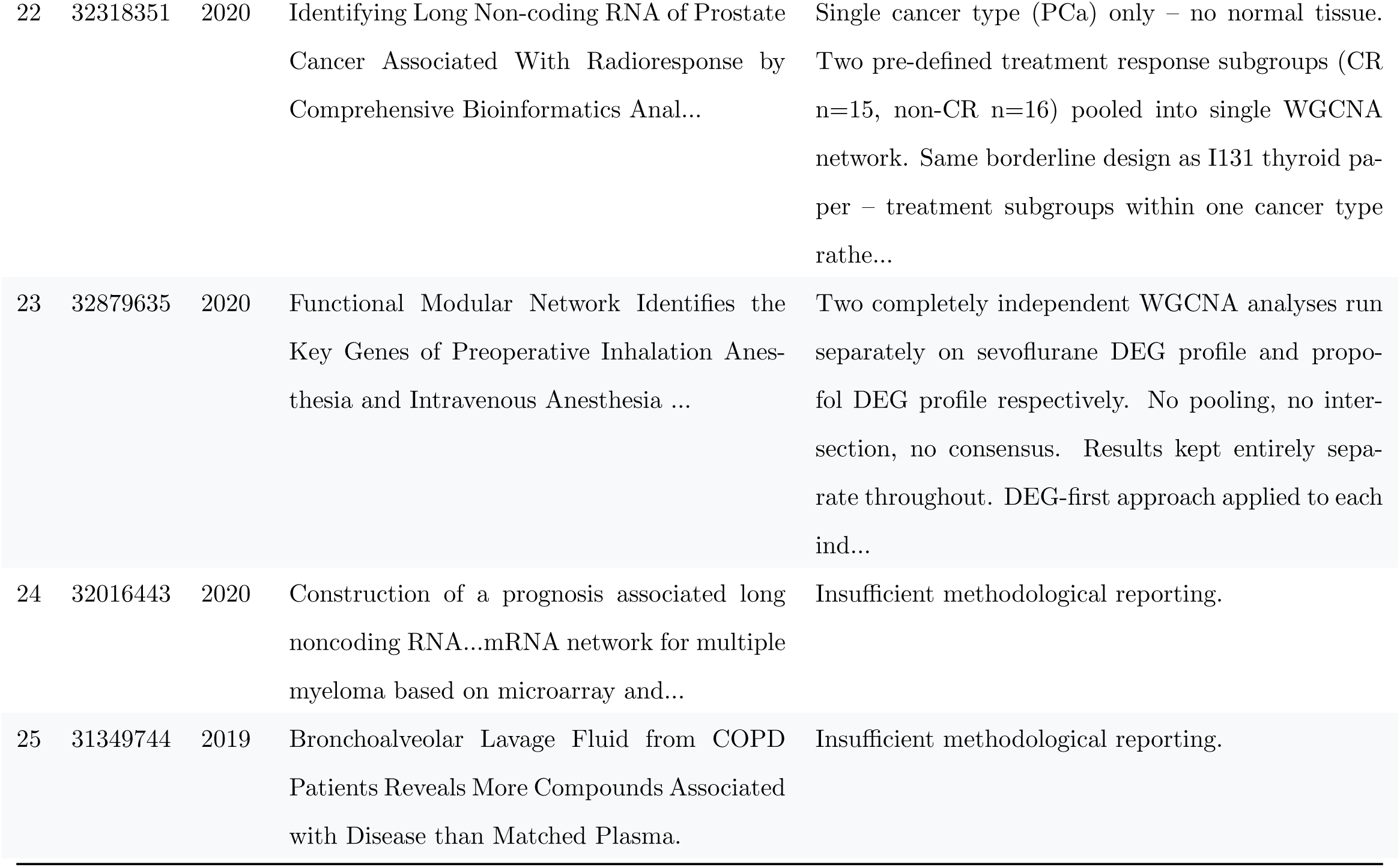
Excluded papers (N = 25) with reason for exclusion.

### Toy example figures

The nine-panel figures for Toy Examples 1 and 2 are presented in the main text (Figures 2 and 3 respectively). The supplemental figure below shows an additional mixed-mean scenario not covered in the main text.

**Figure S1:**
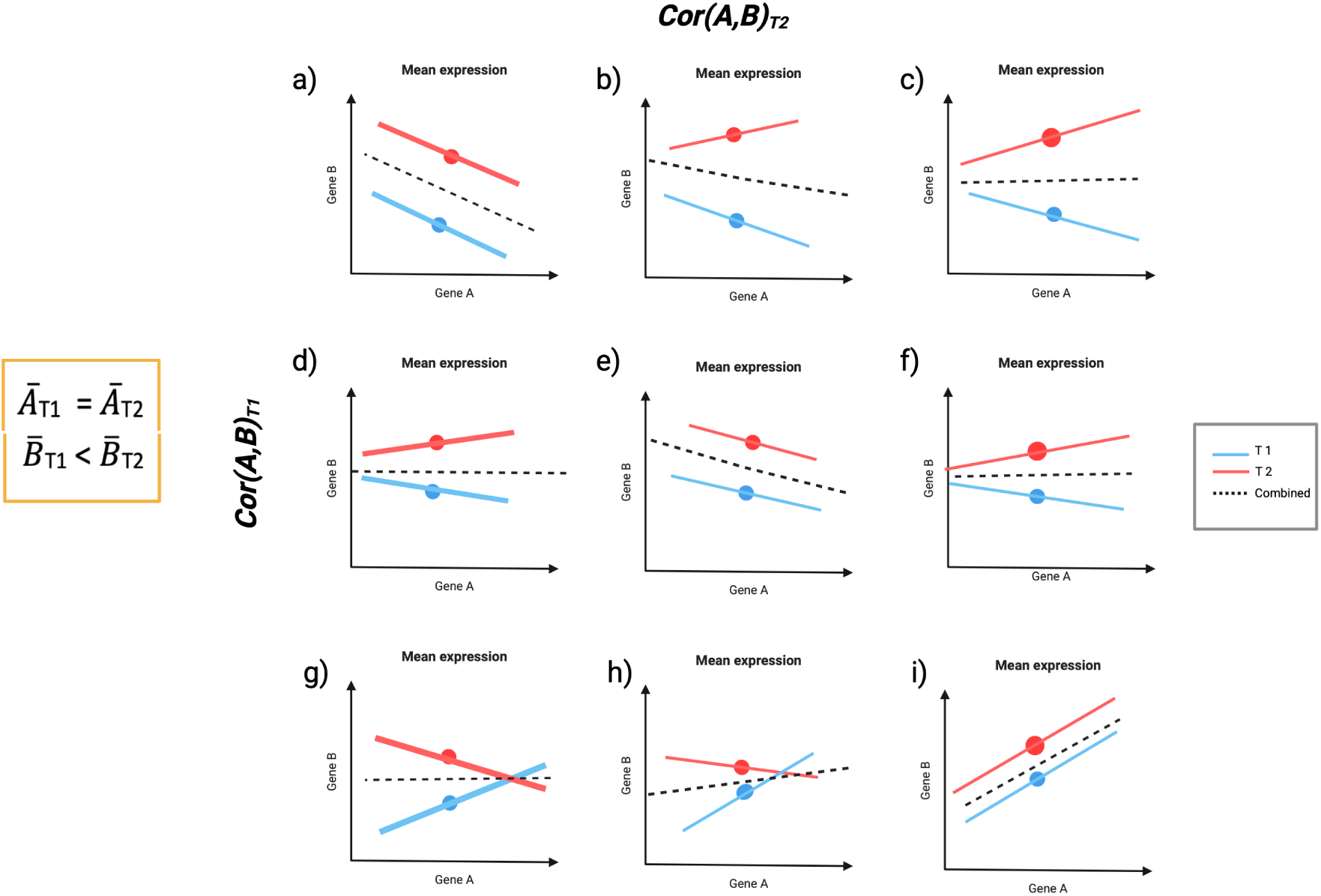
Impact of merging data from two populations with equal mean expression for Gene *A* and different mean expression for Gene *B*. Panels a)-i) show nine possible correlation scenarios between Gene *A* and *B*. Blue and red lines represent correlations in Population T1 and T2 respectively, while dotted lines show correlations in the pooled data. Diagonal panels a), e), and i) demonstrate accurate correlation detection when both populations share similar correlation patterns. Other panels illustrate how ignoring population structure can lead to misidentified correlations when populations exhibit different correlation patterns.

**Figure S2:**
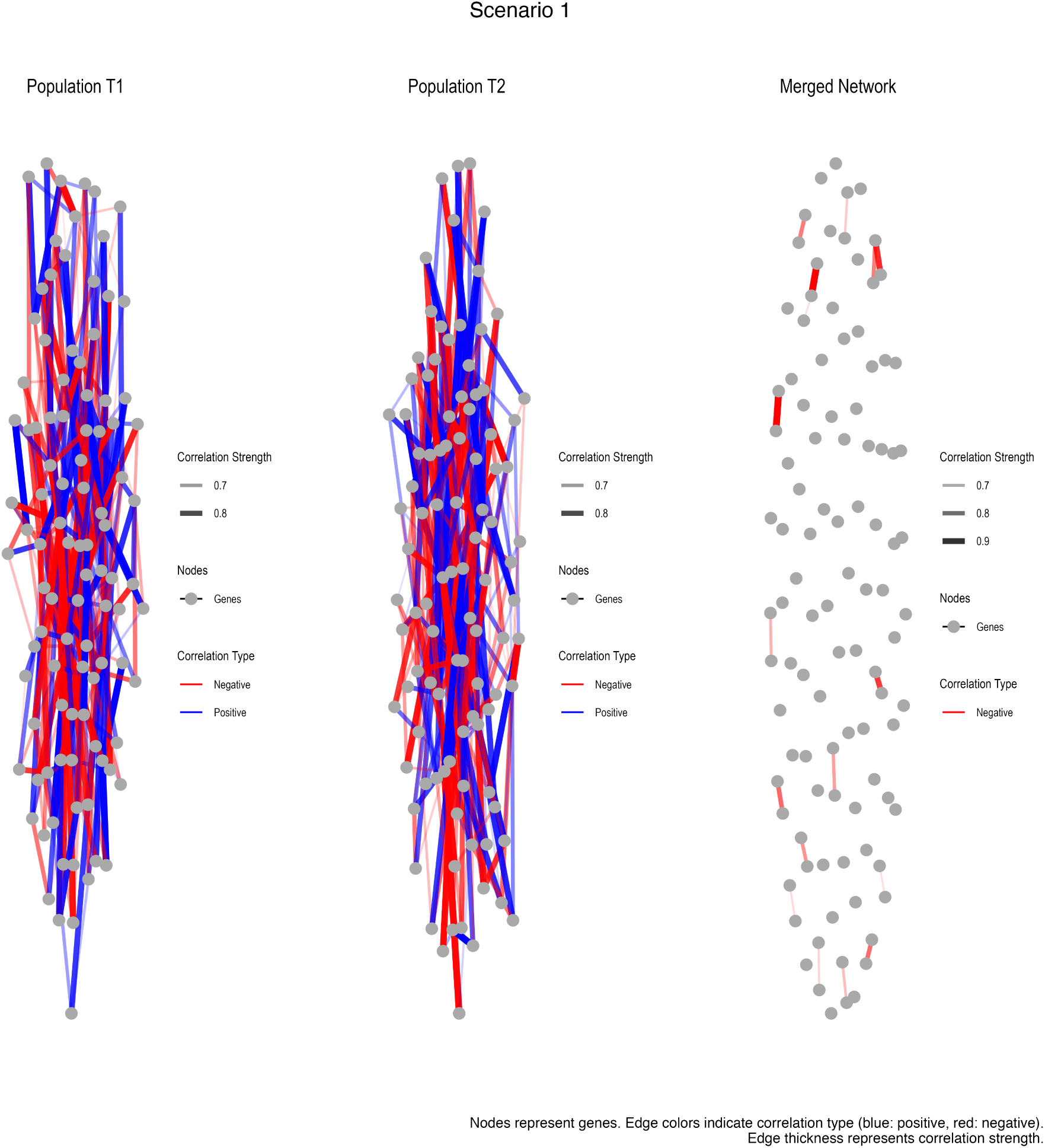
Simulated networks generated for Scenario 1, where Populations T1 and T2 have the same mean expression and network structure.

**Figure S3:**
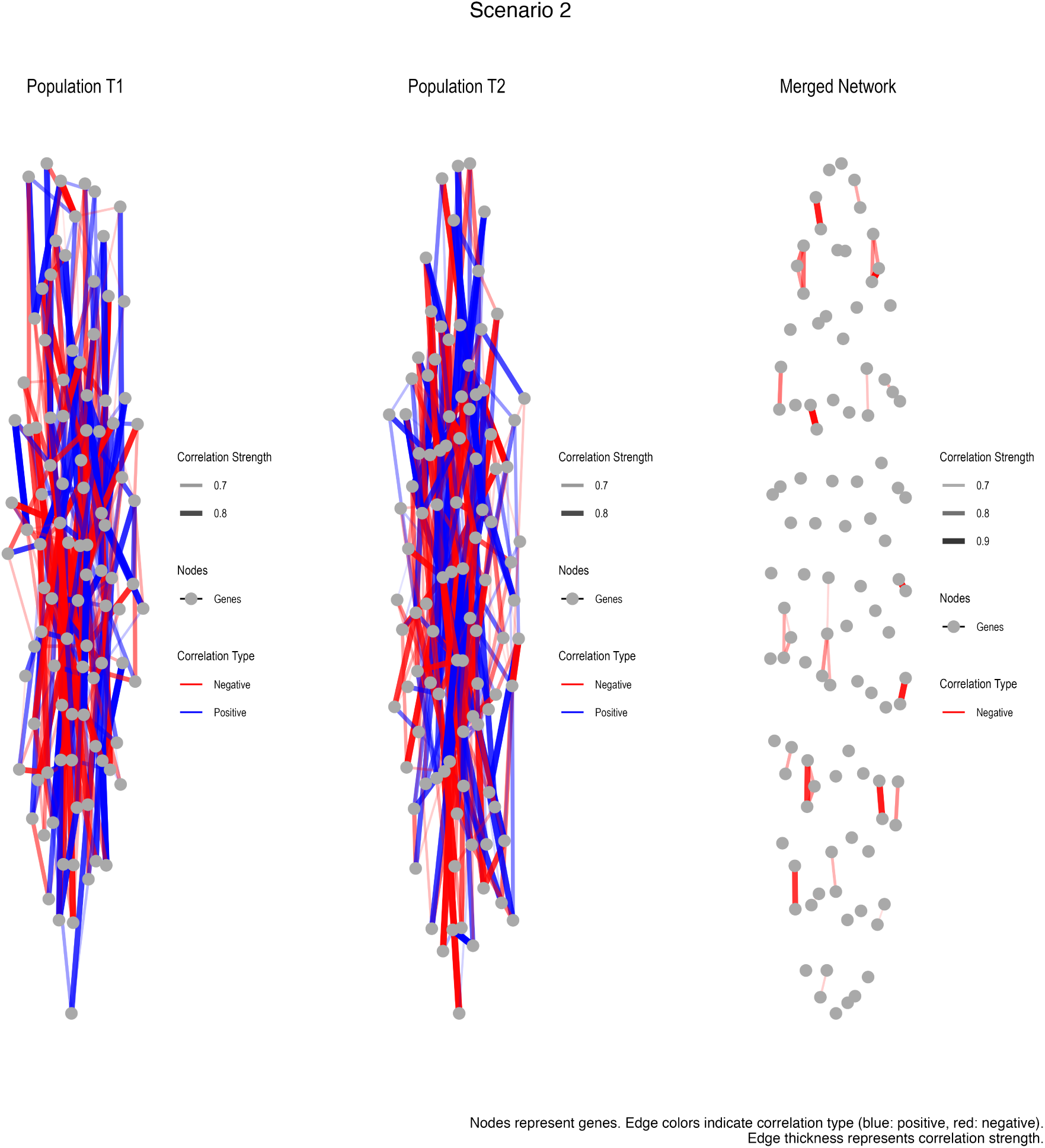
Simulated networks generated for Scenario 2, where Populations T1 and T2 have different mean expressions but similar network structures.

**Figure S4:**
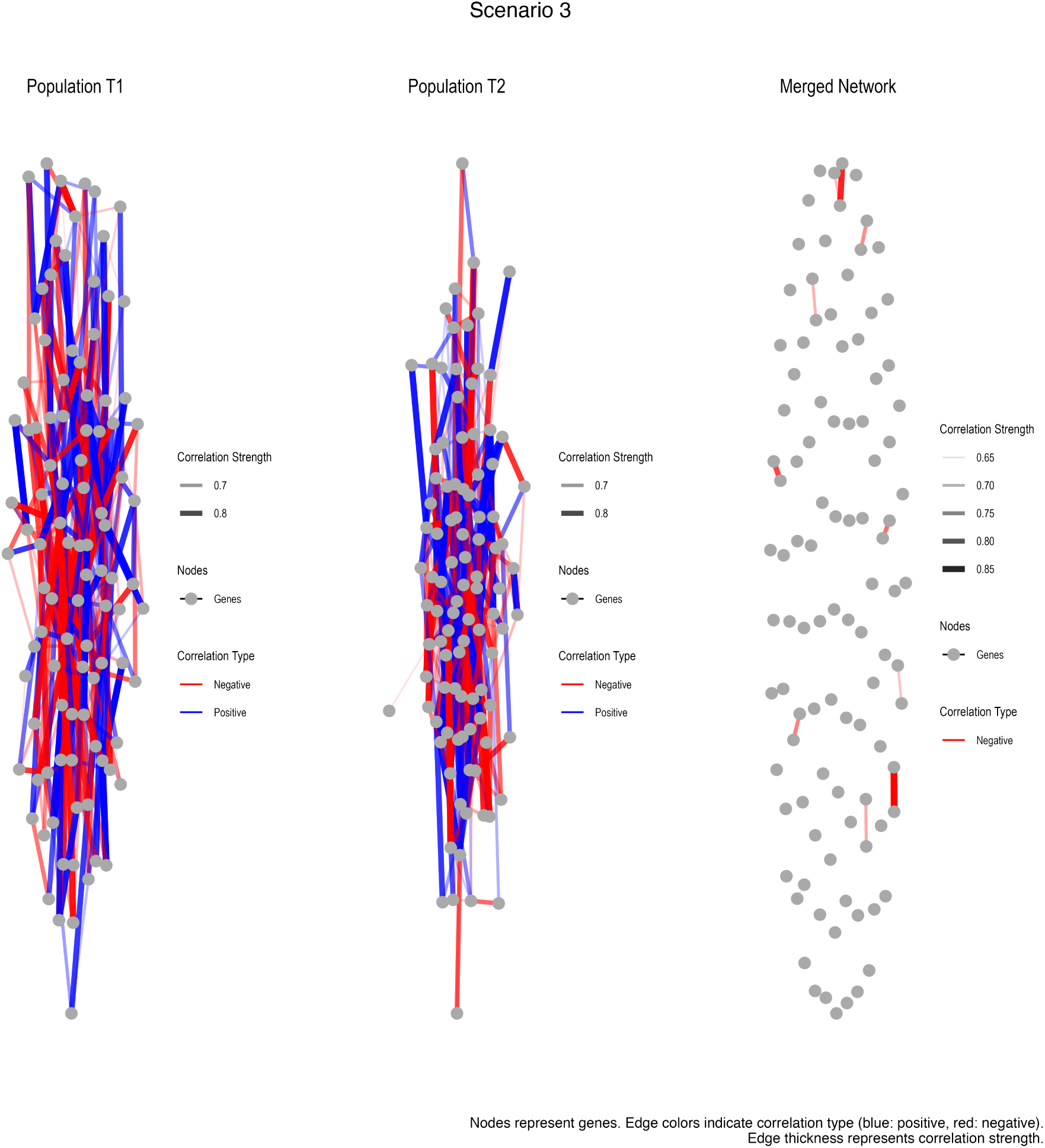
Simulated networks generated for Scenario 3, where Populations T1 and T2 have the same mean expression but different network structures.

**Figure S5:**
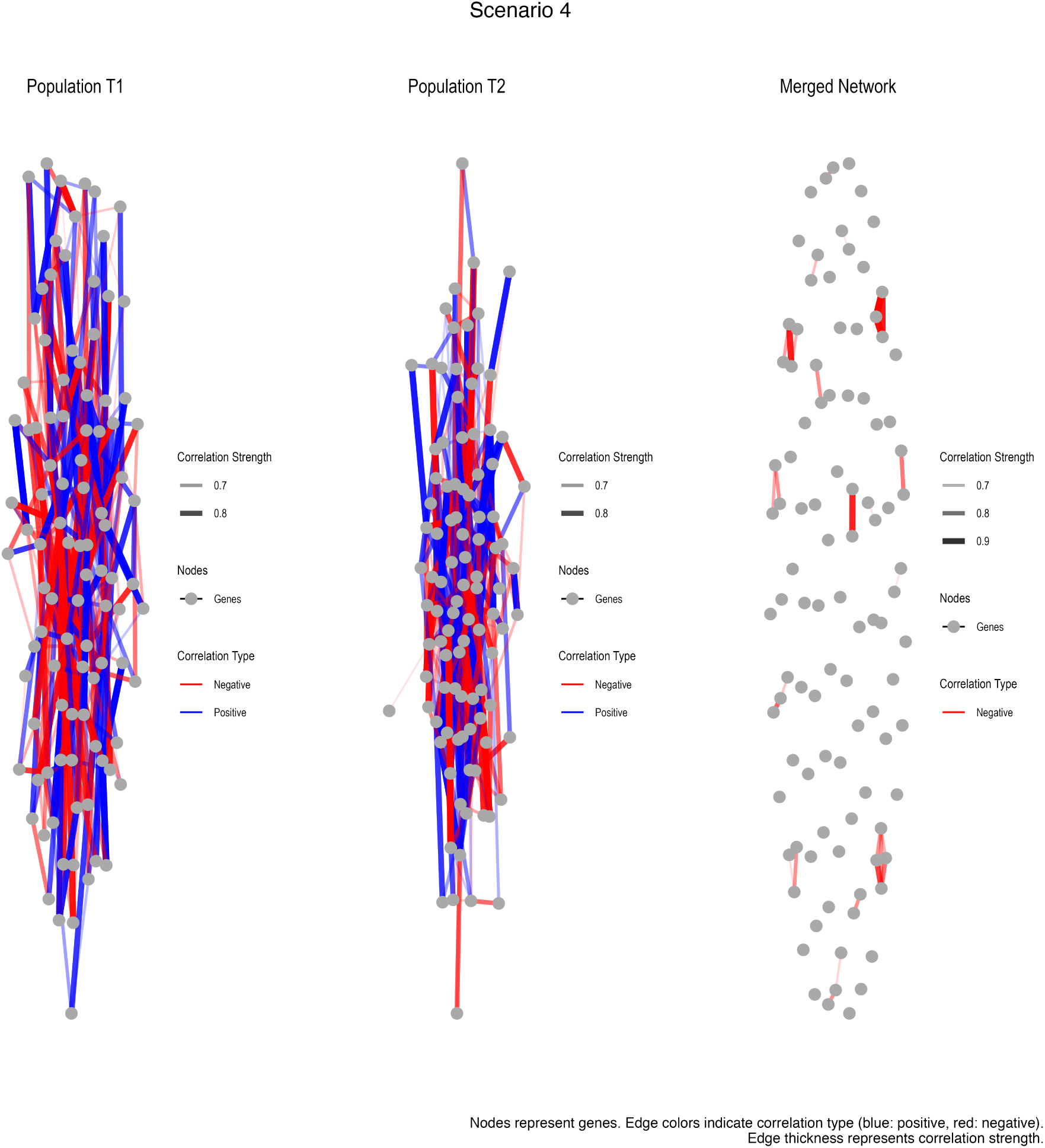
Simulated networks generated for Scenario 4, where Populations T1 and T2 have different mean expressions and network structures.

### Covariance matrix construction and regularisation

Gene expression data were simulated using a block-diagonal covariance matrix derived from the community structure of the Barabási–Albert network, rather than directly from the network’s edge weights. Specifically, gene communities were first identified in the BA graph using the Louvain algorithm [Blondel et al., 2008], treating edge weights as absolute values to satisfy the non-negativity requirement of the algorithm. A covariance matrix Σ was then constructed for the *pgenes* = 500 genes as follows. The diagonal was initialised to the base variance *σ*^2^ = 1.0. For each pair of genes (*i, j*) belonging to the same community, the off-diagonal entry was set to Σ*_ij_* = *ρ* · *σ*^2^, where *ρ* = 0.85 is the within-community correlation. Gene pairs in different communities were treated as independent, with Σ*_ij_* = 0.

The BA edge weights do not directly enter the covariance structure; the scale-free topology contributes only through the community size distribution implied by the Louvain partition. Within-community correlation is set uniformly to *ρ* = 0.85 regardless of edge weights in the underlying graph, and between-community correlation is exactly zero. This block-diagonal Gaussian formulation is a deliberate simplification that isolates the effect of group membership on module structure from the additional complexity of graded within-module correlation decay.

This block-diagonal construction guarantees positive definiteness without regularisation, since the minimum eigenvalue of a block-diagonal correlation matrix of this form is 1 − *ρ* = 0.15 *>* 0. Nevertheless, a regularisation safeguard was included in the implementation: if the minimum eigenvalue *λ*_min_ ≤ 0 (which did not occur in practice under the simulation parameters used), the matrix would be adjusted as

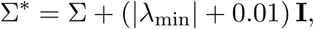

ensuring all eigenvalues are strictly positive.

For scenarios requiring structural differences between populations (Scenarios 3 and 4), two independent modifications were applied to the second population prior to covariance construction: 30% of edges in the BA graph were rewired (removed and replaced with randomly selected new edges, preserving edge count), and 30% of gene community memberships from the original Louvain partition were randomly reassigned to different communities. These two perturbations are independent—the perturbed memberships are not derived by re-running community detection on the modified graph. The covariance matrix for *T* 2 was constructed from the perturbed community assignments as described above.

### Edge threshold for network visualisation and edge preservation

Two distinct edge thresholds are used in the analysis and should not be conflated. For visualisation purposes, a separate correlation-threshold graph was constructed from the pooled expression data using the igraph package: edges were included between gene pairs whose absolute Pearson correlation exceeded 0.6 on the pooled 300 × 500 expression matrix. This graph was used solely for network visualisation (Figure 1) and was not used in any WGCNA module detection or preservation analysis.

For the WGCNA analyses, no edge threshold was applied prior to module detection. Adjacency matrices were computed directly from the expression matrices using soft-thresholding with power *β* = 6 and a signed hybrid network type, yielding fully dense 500 × 500 adjacency matrices in which every gene pair receives a non-zero weight. For the computation of topology metrics (modularity, clustering coefficient, and network density; Figure 5B) and edge preservation between individual and merged networks (Figure 4D, Figure 5A), these fully dense WGCNA adjacency matrices were binarised using a threshold of 0.1 prior to graph construction and edge counting. Without this threshold, edge density and clustering coefficient are trivially near 1.0 for any soft-thresholded WGCNA adjacency matrix, as every gene pair nominally constitutes an edge. The threshold of 0.1 was chosen to retain gene pairs with non-trivial co-expression signal after soft-thresholding. The sensitivity of the topology and edge preservation results to this choice can be assessed by varying the threshold between 0.05 and 0.3. To verify that the conclusions regarding global topology preservation are not an artefact of this choice, we repeated the topology and edge preservation calculations across a range of binarisation thresholds (*τ* ∈ {0.05, 0.10, 0.15, 0.20, 0.25, 0.30}) for all four scenarios across all 20 simulation seeds. Results are shown in Figure S6. Panel A demonstrates that modularity, edge density, and clustering coefficient remain closely aligned between T1, T2, and the merged network across all thresholds and all four scenarios—including Scenarios 3 and 4 where module membership is substantially disrupted. This confirms that the apparent preservation of global topology metrics is a robust property of soft-thresholded WGCNA adjacency matrices and not an artefact of the 0.1 threshold choice. Panel B shows that edge preservation is consistently high in Scenarios 1 and 2 but decreases with increasing threshold in Scenarios 3 and 4, particularly for the T1 vs T2 comparison, reflecting genuine structural divergence between populations. Together, these results confirm that global topology metrics are uninformative about module-level disruption regardless of the binarisation threshold chosen.

### Sensitivity analysis

To assess the robustness of the simulation findings to the choice of network size and structural divergence rate, we conducted a sensitivity analysis spanning a grid of network sizes (*pgenes* ∈ {250, 500, 1,000} genes) and rewiring rates (*r* ∈ {10%, 30%, 50%}) for Scenarios 3 and 4, yielding 18 conditions in total. All other parameters were held constant at the values used in the main analysis (*n* = 150 samples per population, within-community correlation *ρ* = 0.85, soft-thresholding power *β* = 6, minimum module size = max(5, ⌊*p/*50⌋)). Module preservation statistics were computed using *n* = 100 permutations per condition. Conditions were distributed across 7 parallel workers using a PSOCK socket cluster.

Results are shown in Figure S7. Module disruption generally increased with rewiring rate across all network sizes and both scenarios, confirming that the 30% rewiring rate used in the main analysis does not produce qualitatively atypical results. Notably, larger networks (*pgenes* = 1,000) showed substantially greater preservation than smaller networks (*pgenes* = 250) at equivalent rewiring rates, with median *Z*_summary_ values of 15.7–38.9 at 1,000 genes compared to 5.9–10.2 at 250 genes across the same conditions. This network size effect is consistent with more stable gene–gene correlation estimates in larger datasets, which produce more robust module detection. Scenario 4 showed consistently similar or lower preservation than Scenario 3 across the full parameter grid, confirming that the compounding effect of bidirectional expression differences on structural disruption is robust to the specific network size and rewiring rate used. The main conclusions of the paper—that merged WGCNA systematically disrupts module membership in the presence of structural divergence, and that this disruption is generally undetectable from global topology metrics—hold across all 18 conditions examined.

Sample size was excluded from the sensitivity grid because *Z*_summary_ statistics from `modulePreservation()` are sensitive to both module size and per-group sample size [Langfelder et al., 2011]. The WGCNA FAQ additionally recommends a minimum of 15 samples per dataset for reliable correlation estimates, with 20 or more preferred [Langfelder and Horvath, 2020]. At *n* = 150 per group, our simulations comfortably exceed this threshold.

**Table S4:** Mean median *Z*_summary_ for T1-versus-merged module preservation across 20 simulation replicates, by scenario. SD: standard deviation; 95% interval: empirical interval across replicates.

| **Scenario** | **Mean median $Z$** | **SD** | **95% interval** |
| --- | --- | --- | --- |
| 1 (baseline) | 25.2 | 3.32 | 19.0–30.7 |
| 2 (expression) | 15.1 | 1.24 | 12.8–17.2 |
| 3 (structure) | 16.8 | 1.48 | 13.8–19.0 |
| 4 (combined) | 11.9 | 1.01 | 10.2–13.5 |

**Figure S6:**
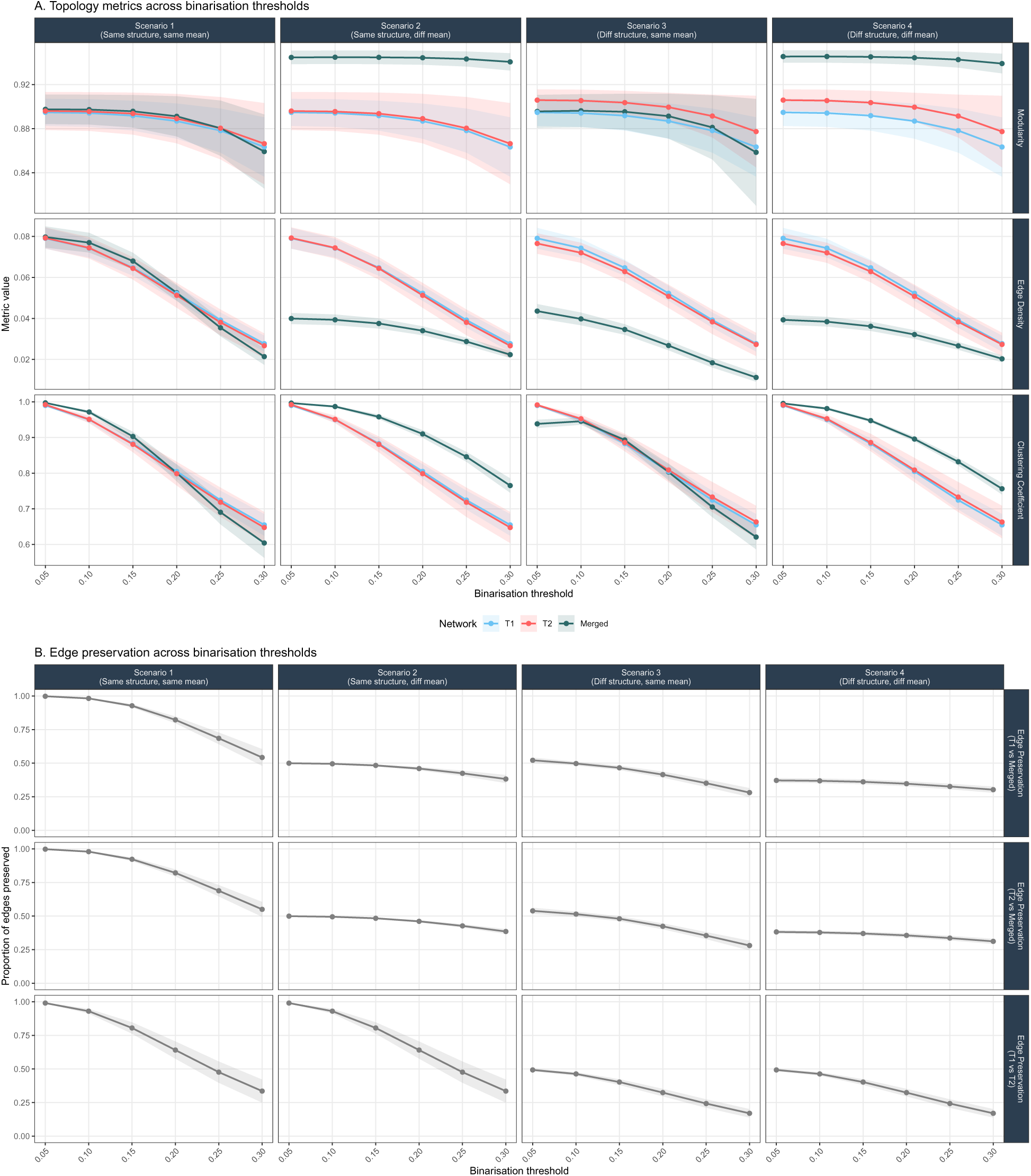
Topology metrics and edge preservation are robust to binarisation threshold choice. Topology metrics (Panel A) and edge preservation proportions (Panel B) computed across six binarisation thresholds (*τ* ∈ {0.05, 0.10, 0.15, 0.20, 0.25, 0.30}) applied to soft-thresholded WGCNA adjacency matrices, for all four scenarios across 20 simulation seeds. Ribbons show mean ± SD across seeds. Panel A: modularity, edge density, and clustering coefficient remain closely aligned between T1 (blue), T2 (red), and merged (teal) networks across all thresholds and scenarios, confirming that global topology metrics appear preserved regardless of threshold choice. Panel B: edge preservation is consistently high in Scenarios 1 and 2 (shared network structure) but decreases with increasing threshold in Scenarios 3 and 4, particularly for the T1 vs T2 comparison, reflecting genuine structural divergence between populations.

**Figure S7:**
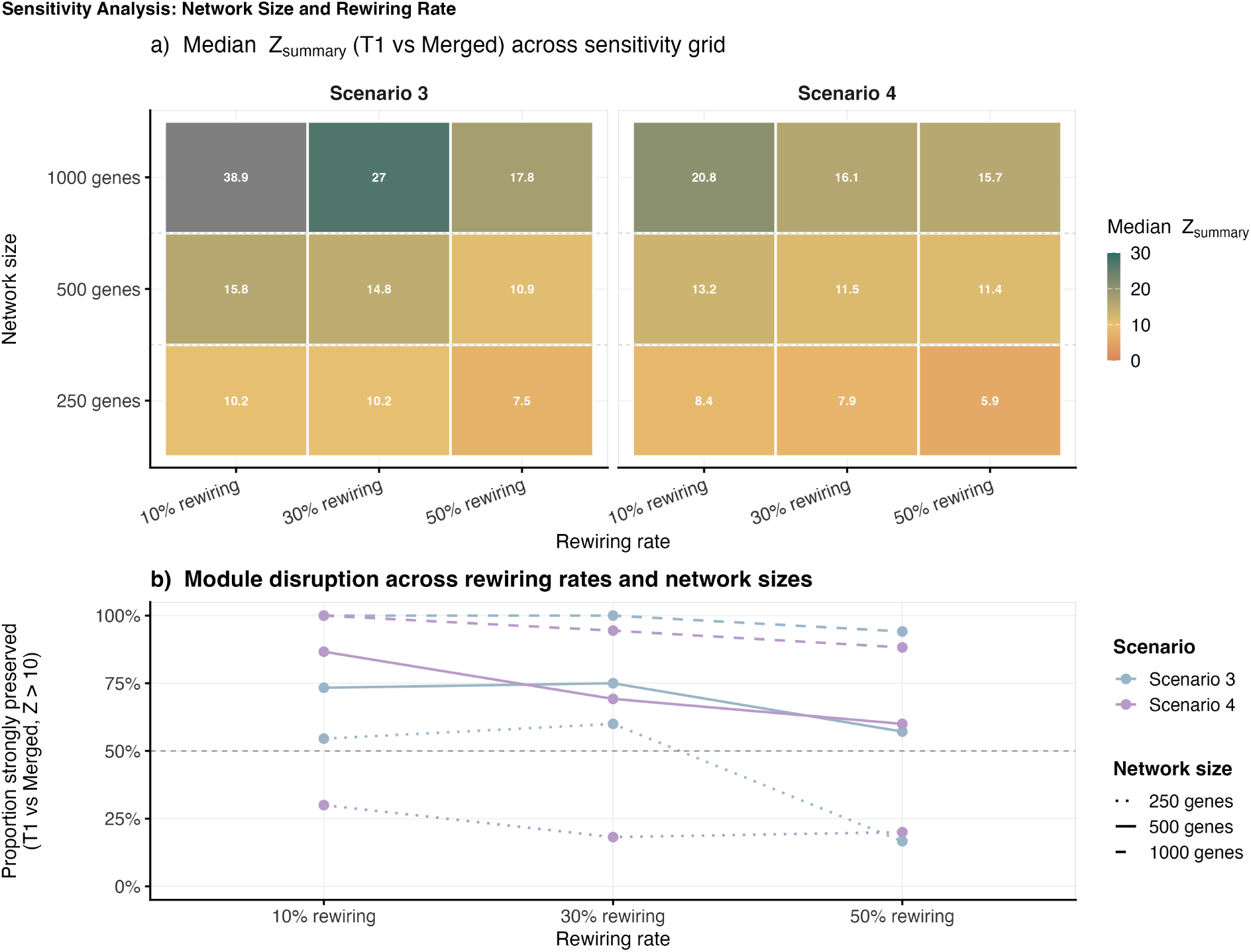
Sensitivitys analysis: module preservation across network sizes and rewiring rates. **a)** Median *Z*_summary_ (T1 versus merged) across network sizes of 250, 500, and 1,000 genes and rewiring rates of 10%, 30%, and 50% for Scenarios 3 and 4. Darker teal indicates stronger preservation; values below 10 fall below the strong preservation threshold. **b)** Proportion of T1 modules strongly preserved (*Z*_summary_ *>* 10) stratified by network size (line type) and scenario (colour); dashed horizontal line marks 50%. Disruption increases with rewiring rate, confirming 30% rewiring is a moderate rather than extreme case. Larger networks show greater preservation, consistent with more stable co-expression estimates. Scenario 4 shows similar or lower preservation than Scenario 3, confirming expression differences compound structural disruption regardless of network size. All conditions: *n* = 100 permutations.

**Figure S8:**
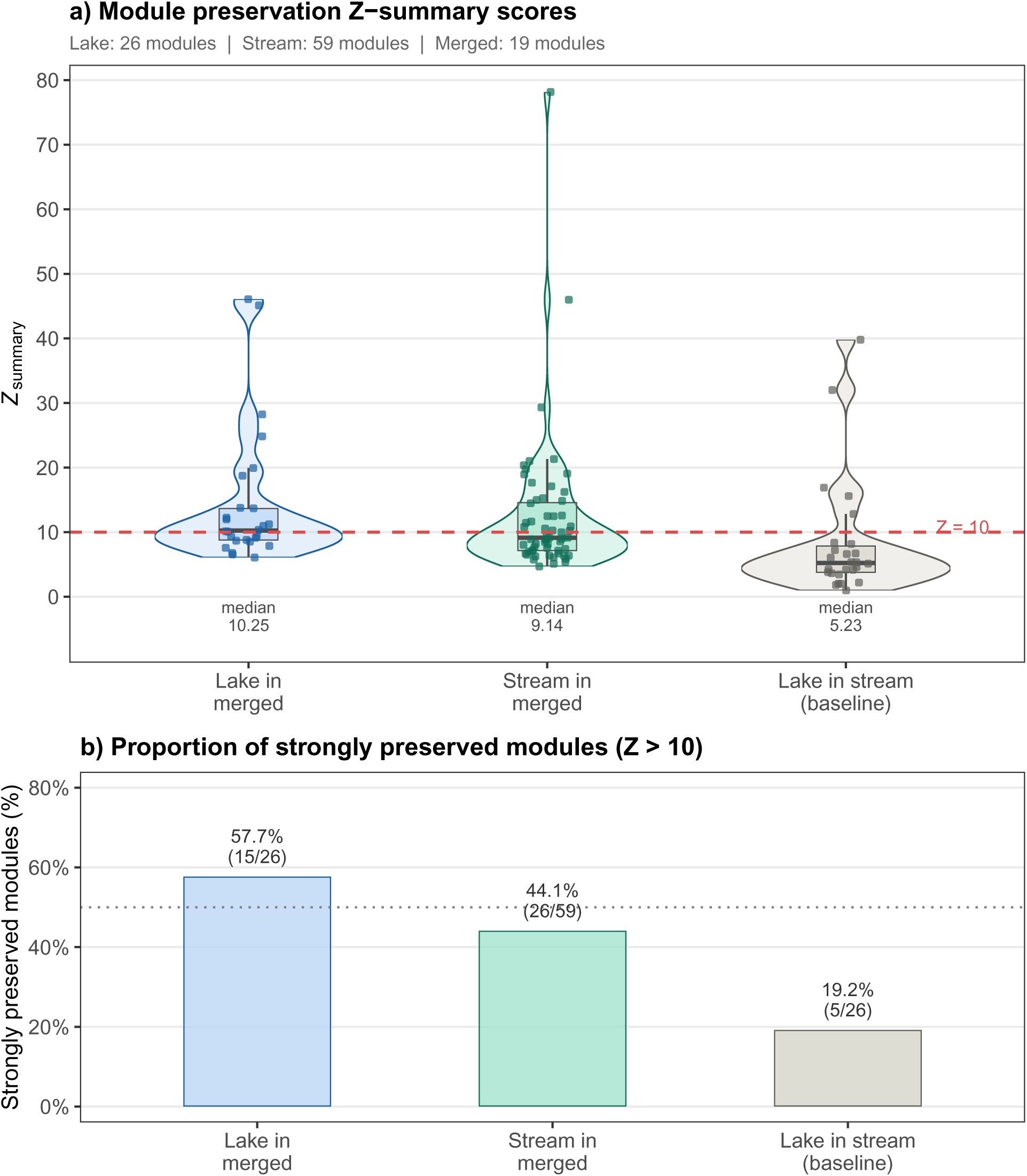
Module preservation in lake–stream stickleback co-expression networks. **a)** Distribution of *Z*_summary_ preservation scores for lake modules in the merged network (blue; *n* = 26), stream modules in the merged network (teal; *n* = 59), and lake modules in the stream network as a biological baseline (grey; *n* = 26). Median *Z*_summary_ values are annotated below each violin. The dashed red line indicates the *Z* = 10 threshold for strong preservation. **b)** Proportion of strongly preserved modules in each comparison: 58% of lake modules in the merged network, 44% of stream modules in the merged network, and 19% of lake modules in the stream network. Data are from wild uncaged stickleback from Roberts Lake and Stream [Lohman et al., 2017].

## Notes

### Competing Interest Statement

The authors have declared no competing interest.

